# Accurate action potential inference from a calcium sensor protein through biophysical modeling

**DOI:** 10.1101/479055

**Authors:** David S Greenberg, Damian J Wallace, Kay-Michael Voit, Silvia Wuertenberger, Uwe Czubayko, Arne Monsees, Takashi Handa, Joshua T Vogelstein, Reinhard Seifert, Yvonne Groemping, Jason ND Kerr

## Abstract

Multiphoton imaging of genetically encoded calcium indicators is routinely used to report activity from populations of spatially resolved neurons *in vivo*. However, since the relationship between fluorescence and action potentials (APs) is nonlinear and varies over neurons, quantitatively inferring AP discharge is problematic. To address this we developed a biophysical model of calcium binding kinetics for the indicator GCaMP6s that accurately describes AP-evoked fluorescence changes *in vivo*. The model’s physical interpretation allowed the same parameters to describe GCaMP6s binding kinetics for both *in vitro* binding assays and *in vivo* imaging. Using this model, we developed an algorithm to infer APs from fluorescence and measured its accuracy with cell-attached electrical recordings. This approach consistently inferred more accurate AP counts and times than alternative methods for firing rates from 0 to >20 Hz, while requiring less training data. These results demonstrate the utility of quantitative, biophysically grounded models for complex biological data.

## Introduction

Linking animal behavior to the underlying neuronal activity requires accurately measuring action potentials (APs) from large populations of neurons simultaneously. Extracellular electrical recordings [65, 84, 117] are widely used to monitor APs in large neuronal populations during behavior [118, 130, 136] but cannot unambiguously assign activity to individual neurons or spatial locations [12, 55]. Alternatively, AP-evoked calcium influx can be detected optically using multiphoton excitation [30] of fluorescent calcium sensors [129]. While this approach can be used to record from both active and silent neurons in anesthetized [70, 120], awake [32, 51] and freely moving animals [114, 143], quantitative readout of neuronal activity requires fluorescence signals to be converted into APs using an inference procedure. For synthetic sensors with a single calcium binding site [129, 57], fluorescence increases from successive APs combine linearly [61, 70, 138] so APs can be inferred by deconvolution [51, 53, 71, 70, 105, 113, 132, 138]. Genetically encoded calcium indicators (GECIs, reviewed in *[64]), on the* other hand, can label neurons with a specific cell type or projection target and track neuronal populations over days with single cell resolution. In particular GCaMP sensors, derived from calmodulin and green fluorescent protein [90], have been repeatedly improved [1, 122, 123, 127] and GCaMP6s approaches the sensitivity of synthetic sensors [18]. However, inferring APs from GCaMP fluorescence remains challenging and unreliable due to AP responses that and are complex, nonlinear and variable over neurons [1, 18, 64]. This complexity, nonlinearity and variability can be qualitatively explained by cooperativity across GCaMP’s 4 calmodulin-derived binding sites, slow and multiphasic kinetics of the indicator, dependence of AP-evoked fluorescence amplitude and shape on total GCaMP concentration [64] and out-of-focus signals from background structures that can increase baseline fluorescence differently for each neuron. However, incorporating these effects into a quantitative description of GCaMP-expressing neurons requires many unknown parameters to be determined. Instead, AP inference methods have been proposed based on phenomenological modeling [29], deep learning with many free parameters but no interpretable model [126] and thresholding [32, 56]. While some of these approaches outperform linear deconvolution on GCaMP data, they produce inaccurate results for many neurons.

Here we describe a sequential binding model (SBM) linking APs to GCaMP6s fluorescence based on biophysical principles. This model quantitatively describes GCaMP6s fluorescence *in vivo* and *in vitro* and allows for more accurate AP inference than previous methods. Given an AP sequence, the SBM describes the binding kinetics that determine concentrations over time for free calcium, calcium bound to endogenous buffers and GCaMP6s binding states with 0 to 4 calcium ions. Solving the resulting differential equations provides GCaMP6s binding state concentrations at each measurement time, which we use to directly predict fluorescence. We fit the SBM to a library of combined optical/electrical recordings from neurons in mouse visual cortex, showing it can capture the AP-fluorescence relationship despite nonlinearity and variability over neurons. We also used the SBM to describe calcium-GCaMP6s interactions *in vitro* by applying the same biophysical framework to fluorescence spectroscopy, isothermal titration calorimetry and stopped-flow fluorescence experiments. Globally fitting the SBM to data from all three*in vitro* binding assays yielded parameters that accurately predicted AP-evoked fluorescence *in vivo*. To use the SBM for AP inference, we developed a GPU-based sequential Monte Carlo algorithm [49] and evaluated accuracy on held-out data. SBM-based AP inference outperformed previous algorithms [29, 32, 105, 126, 132], producing more accurate firing rates, AP counts and AP times, while modeling the effect of AP discharge on fluorescence in interpretable biophysical terms.

## Results

### AP-evoked GCaMP6s fluorescence *in vivo*

To characterize how AP discharge sequences give rise to GCaMP6s fluorescence changes, we imaged neuronal populations expressing virally transfected GCaMP6s in L2/3 mouse visual cortex while juxtasomally recording one neuron’s spontaneous APs (Figure 1a, n = 26 neurons, 10 animals, 15464 APs, 9.4 hours total, mean firing rates 0.008 to 12.5 Hz, median 0.16 Hz). Based on previous observations in L2/3 mouse visual cortex [93], we identified 4 recordings as putative interneurons from the electrophysiological data based on higher mean spontaneous firing rates, AP waveforms with larger afterhyperpolarizations (AHPs) and shorter peak-to-AHP latencies than in pyramidal neurons. We extracted fluorescence from each neuron’s somatic cytosol using a semi-automated algorithm (Figure 1 - figure supplement 1, methods).

**Figure 1:**
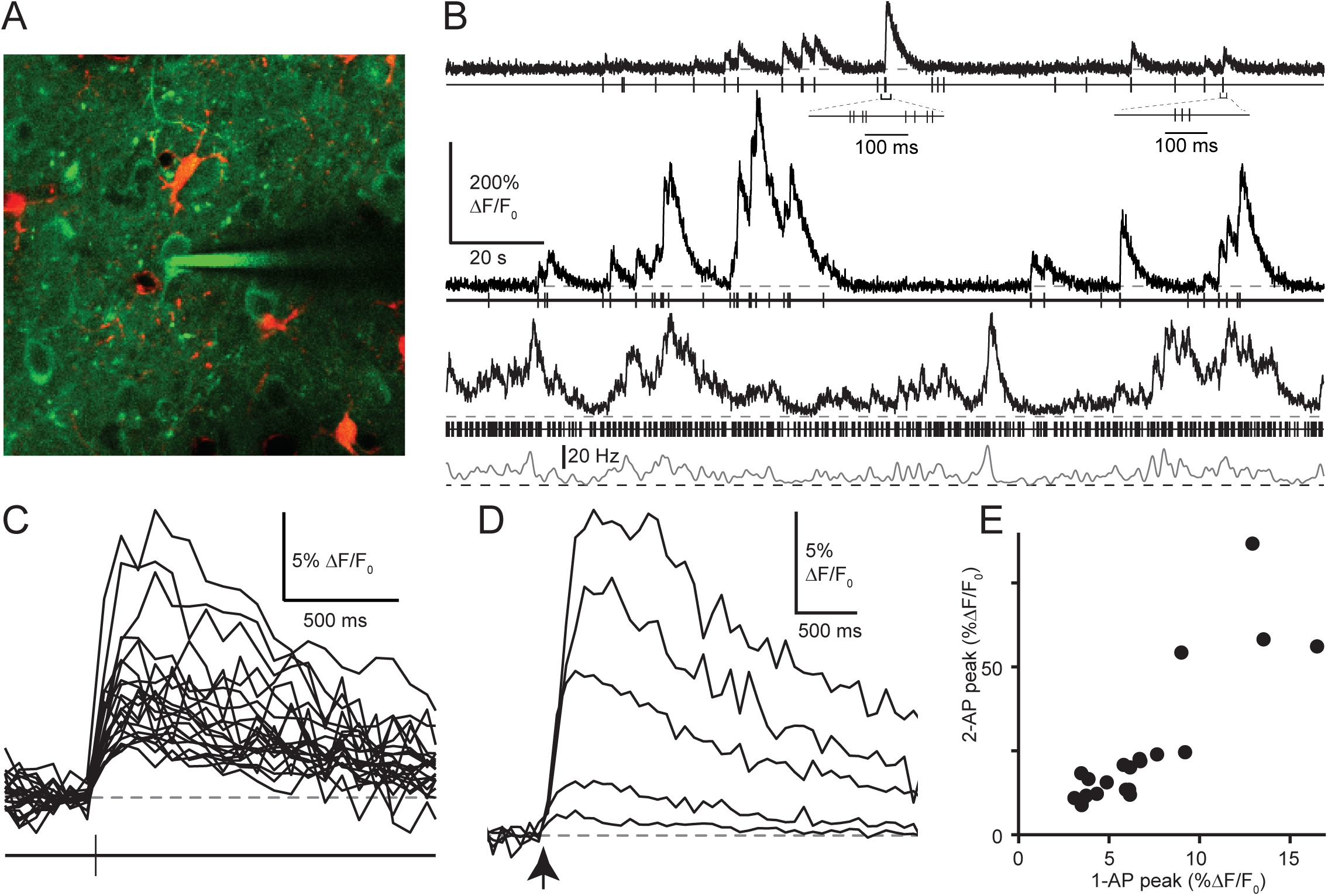
**(A)** Two-photon image of neuronal population expressing GCaMP6s (green) in L2/3 of mouse visual cortex, with astrocytes stained using sulforhodamine 101 (red) and juxtasomal electrical recording of APs in a single neuron. **(B)** Simultaneous electrical recording of AP times and imaging of neuronal fluorescence in 3 different neurons. For the third, an interneuron, firing rate was calculated by Gaussian filtering of the AP sequence with *σ* = 500 ms. Dashed lines indicate 0% Δ*F/F*_0_ and 0 Hz. **(C)** Average fluorescence evoked by isolated single APs (no APs in the preceding 5.5 s) in 22 pyramidal neurons. **(D)** Average fluorescence evoked by single APs and bursts of 2-5 APs in a pyramidal neuron. Bursts were defined as groups of APs spanning < 200 ms with no other APs 5.5 s before the first AP in the burst. **(E)** Comparison of peak amplitude evoked by 1 vs. 2 APs; same neurons as in (C). Peak amplitude was calculated as the maximum of the average fluorescence response to 1 or 2 APs for each neuron.

These recordings exhibited a wide variety of AP discharge patterns, evoking a diverse range of fluorescence signals (Figure 1b). In pyramidal neurons, isolated single APs and brief AP bursts evoked transient fluorescence increases (Figure 1b, upper, peak latency 173 *±*70 ms), but during periods of prolonged AP discharge fluorescence did not return to baseline between bursts (Figure 1b, middle and lower). Isolated single APs were sufficient to increase fluorescence in pyramidal neurons [18], but response shape exhibited considerable variability across neurons (average 1-AP peak ranged from 3.0%-18.7% Δ*F/F*_0_, mean 7.6%, s.d. 4.7%, Figure 1c) that could not be explained by variation across animals or differences in expression time (Figure 1 - figure supplement 2). The fluorescence evoked by AP burst of 2 or more APs grew supralinearly with the number of APs (Figure 1d, burst duration < 200 ms), with the peak fluorescence evoked by 2 APs ranging from 2-7 times the 1-AP peak (Figure 1e, mean 3.5, s.d 1.2, n = 22). In interneurons, fluorescence increased during periods of more frequent AP discharge (Figure 1b, lower) but single APs did not evoke discernible fluorescence changes (maximum peak amplitude 0.2% Δ*F/F*_0_, Figure 1- figure supplement 3). These results show that accurately inferring APs from GCaMP6s fluorescence requires methods capable of dealing with both nonlinearity and variability over neurons.

We used this dataset to test several existing AP inference algorithms: the phenomenological model-based method MLspike [29], a threshold-based approach (thr-*σ*) [32] and a deep-learning method [126] trained on the current data (c2s-t) or using published parameters (c2s-s). While these approaches successfully detected single APs and bursts in some cases, for a majority of neurons either less than half the APs were detected or more than half of inferred APs were false positives, for every method tested (Figure 1 - figure supplement 4). These results show that inferring AP sequences from GCaMP6s fluorescence signals remains a challenging open problem. We therefore developed a quantitative model linking AP discharge to fluorescence in GCaMP6s-expressing neurons, to better understand the relationship between the two and to provide a mathematical foundation for improved AP inference.

### A sequential binding model for GCaMP6s-expressing neurons

To quantitatively link APs to GCaMP6s fluorescence, we constructed a biophysical model based on the chain of causal effects through which APs cause fluorescence changes. First, the AP depolarizes the somatic membrane and calcium ions enter the neuron through ion channels [5, 17, 40, 60, 107, 124] (reviewed in [15, 62]). This temporary increase in cytosolic calcium concentration is further shaped by physiological processes, including buffering by endogenous proteins [60, 80, 111, 116] and extrusion [60
, 80, 110, 111, 115, 116]. At the same time, some calcium ions bind reversibly to GCaMP6s, which contains a calmodulin-derived protein domain with 4 binding sites [90]. Calcium binding to GCaMP6s increases its fluorescence [18], with proposed mechanisms based on conformational changes that alter fluorophore protonation [133, 2, 59, 122, 7]. We reasoned that since the individual parts of this system have been extensively studied and quantified, by modeling them together we could quantitatively predict fluorescence from APs and infer APs from fluorescence (Figure 2a). Furthermore, our combined optical/electrical recordings could be used to fit model parameters, assess the quality of model fits to data and evaluate the accuracy of model-based AP inference.

**Figure 2:**
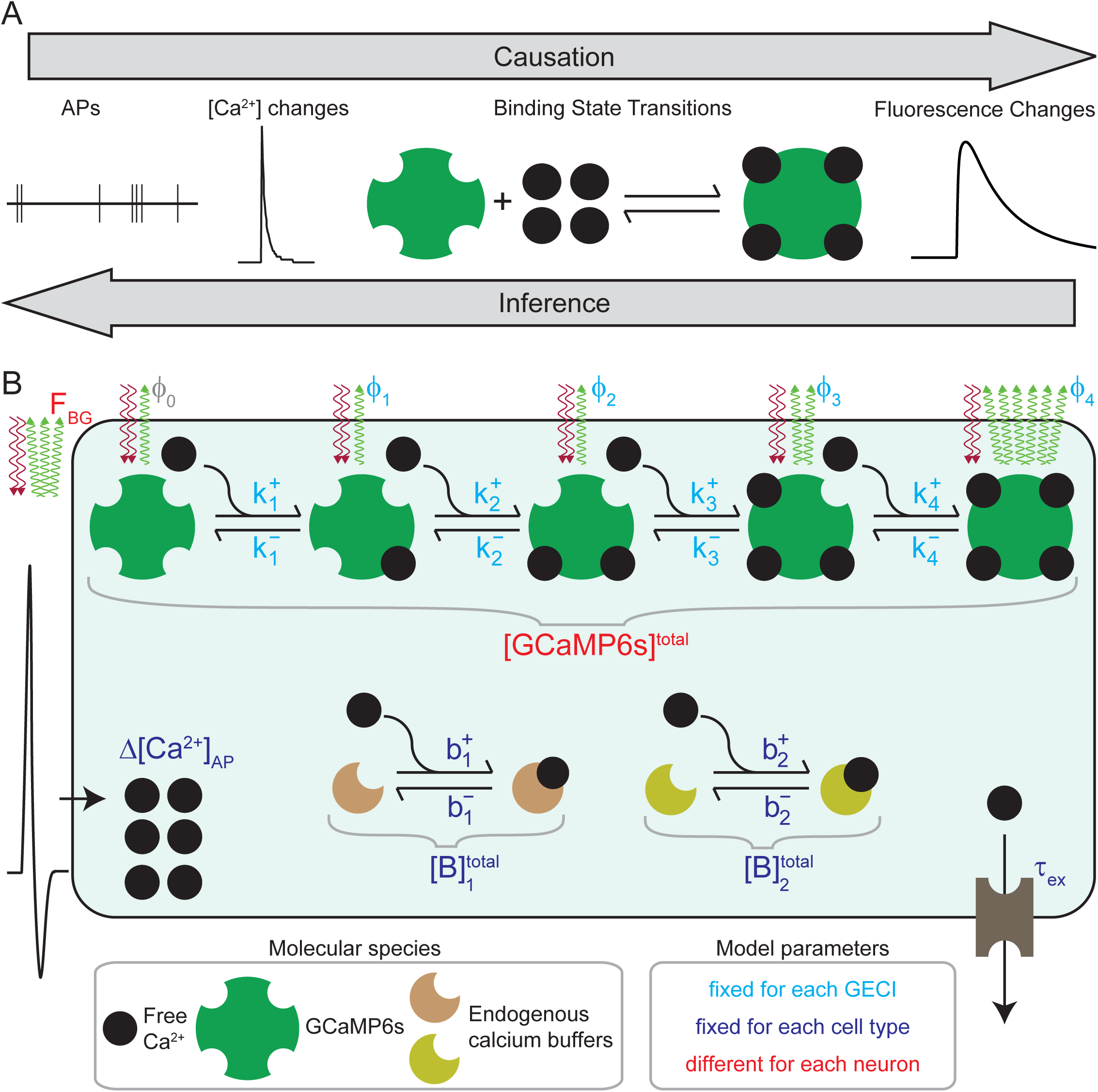
**(A)** Conceptual illustration showing biophysical modeling used to quantitatively link fluorescence and APs. AP discharge, calcium kinetics, calcium-dependent binding state transitions and the resulting fluorescence changes are linked by a causal biophysical model (left to right) and inference procedure (right to left). **(B)** Diagram showing the sequential binding model (SBM). Each AP causes a calcium influx that increases cystosolic free calcium concentration by Δ[Ca^2+^]_AP_. Binding of calcium to GCaMP6s then proceeds with on-rates 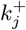 and off-rates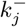, and binding to endogenous buffers with on-rates 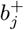 and off-rates 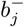 [GCaMP6s]^total^ denotes the total GECI concentration in the neuron and the total concentration of each endogenous buffer is 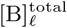 The calcium dependence of GCaMP6s fluorescence is described by brightness values *ϕ* _*j*_ for each binding state Ca_*j*_ GCaMP6s. The total fluorescence predicted by the SBM is calculated (eq. 6-7) based on the current GCaMP6s binding state concentrations, the brightness *ϕj* of each binding state and the background fluorescence *F*_BG_.

We therefore developed a sequential binding model (SBM) of GCaMP6s-expressing neurons (Figure 2b). The core idea of the SBM is to model each individual binding step as calcium binds the four sites of GCaMP6s:

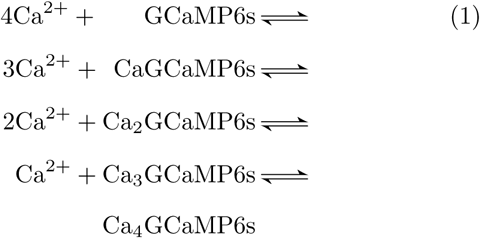

These four reversible binding reactions involve free calcium and five possible GCaMP6s binding states, with 0 to 4 ions bound. We modeled these reactions using standard mass action kinetics [87, 137]: each forward rate is proportional to the concentrations of free calcium and the previous binding state, while each backward rate is proportional to the concentration of the next binding state:

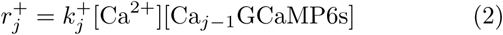

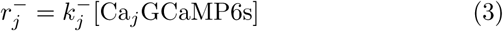

The constants of proportionality are given by the kinetic rate:constants 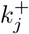 (on-rates) and 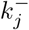 (off-rates) for each binding step, which together describe calcium’s interactions with GCaMP6s. The SBM includes a different total indicator concentration [GCaMP6s]^total^ for each neuron to incorporate varying levels of indicator protein expression [64].

In order to link APs and fluorescence, however, the SBM must describe not only the interaction between calcium and GCaMP6s but also the effects of AP discharge, endogenous buffering and extrusion on calcium concentration. We modeled AP discharge as an increase in free calcium concentration by Δ[Ca^2+^]_AP_, and binding to endogenous buffers with two additional reactions:

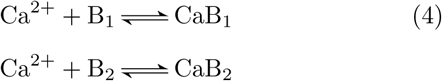

where B_1_ and B_2_ are endogenous buffers with on-rates 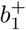 and 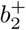, off-rates 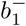 and 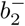 and total concentrations 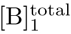 and 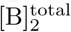. Extrusion was included as the reaction Ca^2+^ → ∅ involving only calcium ions, with the rate given by:

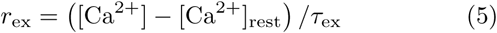

where *τ*_ex_ is the extrusion time constant and [Ca ^2^+]_rest_ is the neuron’s resting calcium concentration. Because GCaMP6s, the endogenous buffers and the extrusion process compete for calcium ions, the SBM’s calcium dynamics are shaped by all three together.

The SBM thus contains 7 reactions and 10 molecular species. Given the current concentrations of these 10 species, the rates of change of all concentrations are given by a global rate equation (Figure 2 - figure supplement 1). The rate equation can therefore be used to calculate the evolution of all molecular species’ concentrations over time, which the SBM then uses to predict the neuron’s fluorescence at the time of each measurement. We modeled the total fluorescence *F*_cyt_ arising from GCaMP6s located in the neuron’s somatic cytosol by adding up the fluorescence arising from each binding state of the indicator:

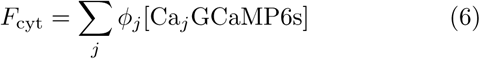

where *ϕ*_*j*_ is the brightness of Ca_*j*_GCaMP6s. For example, when each *ϕ*_*j*_ is greater than the previous value *ϕ*_*j−*1_, GCaMP6s fluorescence increases for every calcium ion bound.Alternatively, if *ϕ*_*j*_ is the same for all binding states with less than four calcium ions bound, only the final binding step increases fluorescence. In this way the SBM model class can describe spectroscopically silent steps, in which binding of a calcium ion does not change the protein’s fluorescence properties.

Since under *in vivo* imaging conditions neuronal fluorescence can exhibit time-varying baseline fluorescence [51, 70] as well as contamination from tissue autofluorescence [125], neuropil background signals [18, 70] and misfolded indicator protein [23, 90], the SBM also incorporated a driftingfluorescence baseline *F*_BL_ and a constant background fluorescence term *F*_BG_. Combining these effects (see methods), the observed fluorescence predicted by the SBM is

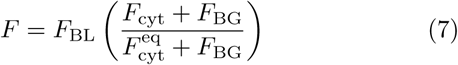

where 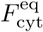 is the equilibrium value of *F*_cyt_ when no APs are discharged and [Ca^2+^] = [Ca^2+^]_rest_. Since rescaling the SBM’s concentration units along with *F*_BG_ does not change the predicted fluorescence, we fixed [Ca^2+^]_rest_ to 50 nM in accordance with previous measurements *in vitro* [60, 80, 116] *and in vivo* [142]. Consequently, all concentration values in the SBM can alternatively be interpreted as multiples of the resting calcium concentration [Ca^^2^+^]_rest_.

Overall, the SBM contains 22 different free parameters (Table 1): 12 GECI-specific parameters describing the calcium-binding and fluorescence properties of GCaMP6s, 8 cell type-specific parameters describing calcium influx, buffering and extrusion and two remaining parameters*, F*_BG_ and[GCaMP6s]^total^, that can vary for each neuron. Having designed the SBM to quantitatively link APs to fluorescence based on established biophysical principles, we next fit its parameters to our *in vivo* GCaMP6s dataset.

### Fitting the SBM to combined optical / electrical recordings *in vivo*

We fit the SBM to our *in vivo* dataset by adjusting all its parameters to give the best possible predictions of fluorescence from AP sequences (Figure 3a-d). For this purpose, we first estimated *F*_BL_ by interpolating fluorescence values from silent periods without APs (Figure 3 - figure supplement 1, see methods; interneurons lacked silent periods and were excluded from fitting, but included when testing AP inference below). We next initialized the SBM parameters to default starting values, using estimates from experimental studies where available (e.g. calcium influx per AP and extrusion rate, full details in methods). To predict fluorescence from a given AP sequence, we started at the beginning of the recording and moved forward in time, incrementing free calcium by Δ[Ca^2+^]_AP_ at each AP while solving the rate equation to determine the concentration over time for all molecular species (Figure 3c, time step 10 ms). We then added up the contributions of all 5 GCaMP6s binding states (eq. 6-7) to generate a prediction of the neuron’s fluorescence (Figure 3b, orange) and compared the result to observed fluorescence values (Figure 3b, black). While calculating fluorescence predictions AP sequences in this way, we then adjusted the SBM parameters to minimized the mismatch between predicted and observed fluorescence values across all pyramidal neurons’ recordings (n = 22), using iterative optimization of with multiple initializations (Figure 3 - figure supplement 2, methods). This fitting procedure resulted in parameters (Table 1) that closely predicted the measured fluorescence signals (Figure 3a-d). Simplified versions of the SBM (Figure 3 - figure supplement 3) without endogenous buffers or variation across neurons in *F*_BG_ or [GCaMP6s]^total^ predicted fluorescence signals less accurately for most neurons (Figure 3 - figure supplement 4, p < 0.005, sign tests). Using more than 2 buffers, more elaborate extrusion mechanisms or a shorter time step did not improve fit quality (p > 0.05).

**Figure 3:**
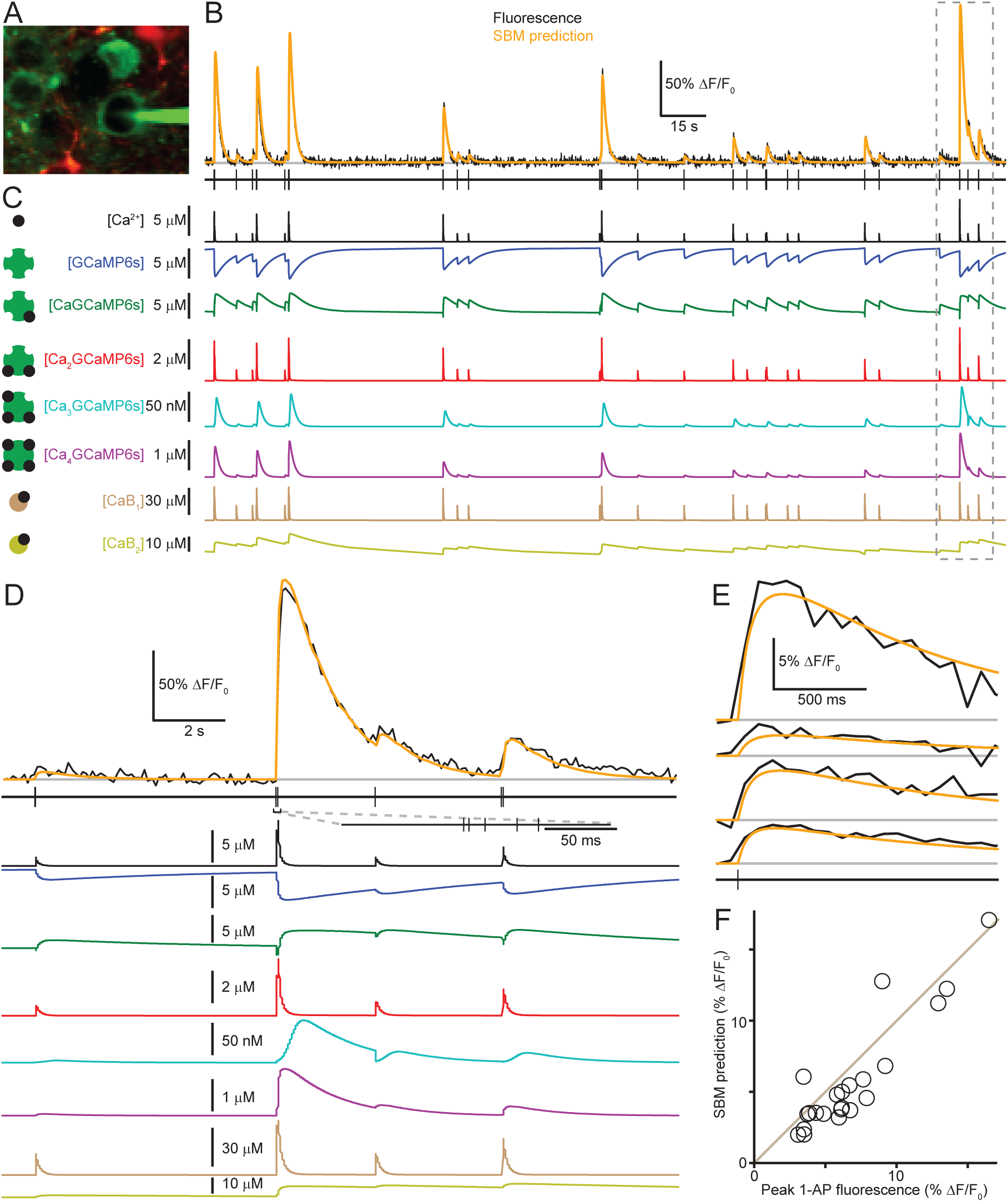
**A** Two-photon image of neuronal population expressing GCaMP6s (green) in L2/3 of mouse visual cortex, with astrocytes stained using sulforhodamine 101 (red) and juxtasomal electrical recording of APs in a pyramidal neuron. **(B)** GCaMP6s fluorescence (black, upper) and simultaneous electrical recording of APs (lower) from the neuron shown in (A). The baseline fluorescence values used for model fitting are shown in gray, and the SBM’s prediction after globally fitting all model parameters to the full dataset is shown in orange. **(C)** Time-varying concentrations of free calcium, GCaMP6s binding states and calcium-bound endogenous buffers calculated by the SBM to generate the prediction in (B). **(D)** Expansion of the period indicated by the dashed box in (B-C). **(E)** Average 1-AP fluorescence increase in four pyramidal neurons (black) and SBM fits (orange). **(F)** Peak fluorescence increase following single APs compared to SBM-fit values for all pyramidal neurons.

**Table 1.** SBM parameters fit *in vivo*

| Symbol | Description | Best-fit value | Range of fit values<sup>##</sup> | Units |
| --- | --- | --- | --- | --- |
| $\Delta[\text{Ca}^{2+}]_{\text{AP}}$ | 1-AP calcium influx | 20.2 | 2.3 - 32.6 | $\mu\text{M}$ |
| $\tau_{\text{ex}}$ | Extrusion time constant | 7.5 | 5.5 - 64.9 | ms |
| $[\text{Ca}^{2+}]_{\text{rest}}$ | Resting free calcium concentration | 50** | | nM |
| $k_1^+$ | First GCaMP6s on-rate | 2.5 | 1.6 - 52.1 | $\mu\text{M}^{-1} \text{s}^{-1}$ |
| $k_2^+$ | Second GCaMP6s on-rate | 16.9 | 1.2 - 20.2 | $\mu\text{M}^{-1} \text{s}^{-1}$ |
| $k_3^+$ | Third GCaMP6s on-rate | 1.1 | 0.5 - 636 | $\mu\text{M}^{-1} \text{s}^{-1}$ |
| $k_4^+$ | Fourth GCaMP6s on-rate | 1069 | 54 - 1766 | $\mu\text{M}^{-1} \text{s}^{-1}$ |
| $k_1^-$ | First GCaMP6s off-rate | 0.1 | 0.1 - 961 | $\text{s}^{-1}$ |
| $k_2^-$ | Second GCaMP6s off-rate | 205 | 2 - 984 | $\text{s}^{-1}$ |
| $k_3^-$ | Third GCaMP6s off-rate | 11.8 | 2.9 - 515 | $\text{s}^{-1}$ |
| $k_4^-$ | Fourth GCaMP6s off-rate | 5.8 | 1.4 - 35.1 | $\text{s}^{-1}$ |
| $\phi_1/\phi_0$ | CaGCaMP6s/GCaMP6s brightness ratio* | 1.0 | 1.0 - 2.6 | *** |
| $\phi_2/\phi_0$ | Ca<sub>2</sub>GCaMP6s/GCaMP6s brightness ratio* | 1.0 | 1.0 - 42.9 | *** |
| $\phi_3/\phi_0$ | Ca<sub>3</sub>GCaMP6s/GCaMP6s brightness ratio* | 1.0 | 1.0 - 81.0 | *** |
| $\phi_4/\phi_0$ | Ca<sub>4</sub>GCaMP6s/GCaMP6s brightness ratio* | 81.0 | 33.1 - 81.0 | *** |
| $[\text{GCaMP6s}]^{\text{total}}$ | Total GCaMP6s concentration | 7.4 **** | | $\mu\text{M}$ |
| $F_{\text{BG}}/F_{\text{cyt}}^{\text{eq}}$ | Background/cytosolic fluorescence at rest | 2.5 **** | | *** |
| $b_1^-/b_1^+$ | Fast buffer dissociation constant | 3.4 | 2.2 - 100 | $\mu\text{M}$ |
| $1/b_1^-$ | Fast buffer time constant | $< 1.0^\#$ | | ms |
| $[B_1]^{\text{total}}$ | Fast buffer concentration | 64 | 0.0 - 154 | $\mu\text{M}$ |
| $b_2^-/b_2^+$ | Slow buffer dissociation constant | 0.59 | 0.1 - 3.6 | $\mu\text{M}$ |
| $1/b_2^-$ | Slow buffer time constant | 17 | 12 - 30 | s |
| $[B_2]^{\text{total}}$ | Slow buffer concentration | 119 | 23 - 129 | $\mu\text{M}$ |

\*For two photon excitation at 920 nm *in vivo*
\*\*Fixed to this value based on previous reports, not fit to data
\*\*\*Unitless ratio
\*\*\*\*Maximum likelihood estimate for the mode of a log-normal fit to parameter's distribution over neurons
#Model fitting procedures results in the lowest allowed value of 1 ms for the fast buffer time constant.

We next examined how individual aspects of the SBM contributed to predicted fluorescence changes using simulations in which some model parameters were perturbed or set to zero. Changing [GCaMP6s]^total^ affected the simulated 1-AP fluorescence peak height, 2-AP peak height and peak timing (Figure 3 - figure supplement 5), as well as the simulated peak free calcium concentration after AP discharge (Figure 3 - figure supplement 6). This is consistent with previous observations that synthetic indicators at high concentrations contribute significantly to Ca^2+^ buffering [60, 80, 116], as well as a theoretical study predicting that GECI expression levels influence AP response amplitude and shape [64]. These simulations also reproduced the nonlinearity of GCaMP6s’s AP responses, with 2 APs evoking 4 to 7 times the fluorescence increases observed for 1 AP depending on the value of [GCaMP6s]^total^ (Figure 3 - figure supplement 7a-b), so that nonlinearity can vary over neurons as observed *in vivo* (Fig*ure 1e).* Simulations omitting one of the endogenous buffers *(Figure 3 -* figure supplement 8) showed that fluorescence responses to 1 or 2 APs were shaped by a fast endogenous buffer (time constant 1̃ms), but the return to baseline after long bursts was shaped by a slow buffer *(>* 10 s). Overall, by incorporating variation over neurons in [GCaMP6s]^total^ and *F*_BG_ together with endogenous buffering and extrusion properties that were fixed for all neurons, the SBM was able to capture variability in the rising and falling phases of observed AP responses (Figure 3e). This allowed the SBM to closely predict the peak fluorescence evoked by single APs over the entire range of peak amplitudes observed (Figure 3f, r = 0.91, n = 22 neurons).

While the SBM was able to predict fluorescence changes from AP sequences, this does not guarantee that a unique set of best-fitting parameter values can be unambiguously identified from the available data. On the contrary, given the SBM’s complexity its parameters may not be fully identifiable from our data, and multiple SBM parameter sets might predict the observed fluorescence signals equally well from the AP sequences. This uncertainty in parameter values is common in detailed biophysical [95] and biochemical models with many parameters [45, 54, 108]. Therefore, to test how tightly SBM parameters can be constrained given our *in vivo* dataset, we repeatedly fit the SBM while excluding one neuron at a time to generate multiple SBM parameter sets, reasoning that if the parameters are identifiable given our *in vivo* dataset then their optimal values should not be significantly altered when removing a single neuron’s data. Comparing the resulting parameter sets showed that most parameters ranged over about 1 order of magnitude, suggesting that they cannot be precisely determined given the available data (Table 1). Nonetheless, when tested on the same AP sequence these different SBM parameter sets produced nearly identical predictions of fluorescence (Figure 3 - figure supplement 9a). The concentrations over time of all molecular species were also highly correlated over SBM parameter sets (minimum correlation 0.98), although the absolute scale of the concentrations differed (Figure 3 - figure supplement 9b). These results show that fitting the SBM to *in vivo* data unambiguously identifies the quantitative mapping from APs to fluorescence, but multiple SBM parameter sets are consistent with the same mapping given our *in vivo* data.

### SBM analysis of *in vitro* binding assays

The SBM describes the interaction of calcium and GCaMP6s as a sequence of protein-ligand binding reactions, allowing quantitative data analysis through the biophysical framework of mass action kinetics. To test whether this framework could describe the same reaction sequence under controlled conditions, we carried out *in vitro* binding assays to measure GCaMP6s-calcium interactions outside of neurons. This allowed us to study these reactions beyond the physiological concentration ranges for GCaMP6s and calcium and using a more diverse set of measurement techniques. We used fluorescence spectroscopy to characterize GCaMP6s’s excitation spectrum at equilibrium (Figure 4a), and how the spectrum changes as calcium concentration is increased. We used isothermal titration calorimetry (ITC) to detect the heat absorbed or released as calcium was titrated into a GCaMP6s solution (Figure 4b), providing readout of calcium binding that does not depend on fluorescence changes. We used stopped-flow fluorimetry to measure fluorescence changes over time after GCaMP6s was rapidly mixed with calcium or a calcium-chelator (Figure 4c). We globally fit the SBM to all data simultaneously (n = 3 spectra, 8 ITC experiments, 19 stopped-flow traces, 4 pools of purified GCaMP6s) to determine kinetic and equilibrium binding properties for GCaMP6s, adjusting all model parameters to achieve the best possible prediction of spectra, heat and stopped-flow fluorescence data (Figure 4d-g, Figure 4 - figure supplement 1, details in methods). This global fit resulted in SBM parameters (Table 2) that closely predicted data from all three binding assays. Similarly to SBM fits on *in vivo* fluorescence data, we found that a range of values for each SBM parameter were consistent with *in vitro* measurements, possibly due to spectroscopically silent binding steps or reactions too fast to resolve using the methods employed. Nonetheless, these results show that the SBM provides a quantitative account of calcium-GCaMP6s interactions in all three binding assays.

We next examined whether the SBM rate constants obtained from *in vitro* binding assays could accurately predict AP-evoked fluorescence changes *in vivo*, as compared to parameters determined exclusively from *in vivo* recordings. To test this, we fixed the *in vitro*-derived rate constants and fit all remaining parameters to *in vivo* fluorescence signals. This resulted in fluorescence predictions similar to those arising from *in vivo*-derived rate constants (Figure 4h). Since this procedure fit the SBM to the same data with fewer free parameters, larger errors would be expected than when fitting all parameters on *in vivo* data alone. However, if the rate constants obtained *in vitro* can accurately describe calcium binding in neurons, then the fitting errors for *in vitro*- vs. *in vivo*-derived rate constants should be approximately equal, and should fall on a line with a slope of one. Comparing each neuron’s r.m.s. error value for the *in vitro*- vs. *in vivo*-derived rate constants in this way, we observed that *in vitro*-derived rate constants produced only slightly higher error values (Figure 4i, r.m.s. error 5.9 *±*3.0 vs. 6.3 *±*3.2% Δ*F/F*_0_), and that the errors fell on a line with a slope close to one (1.06, 95% confidence interval 1.02 - 1.09). Fits using *in vitro*-derived rate constants also predicted *in vivo* fluorescence more accurately than simplified versions of the SBM that omitted endogenous buffers or variation in *F*_BG_ or [GCaMP6s]^total^ over neurons but were fit to *in vivo* data alone (Figure 3 - figure supplement 4). These results show that *in vitro*-derived rate constants lie within the range of SBM parameter sets capable of accurately describing GCaMP6s fluorescence *in vivo*, and that a single set of rate constants can accurately describe both scenarios when the different *in vitro* and cytosolic environments are taken into account. In contrast, previous phenomenological fitting approaches based on rise times and Hill exponents have required different parameter values *in vivo* and *in vitro* [61].

**Figure 4:**
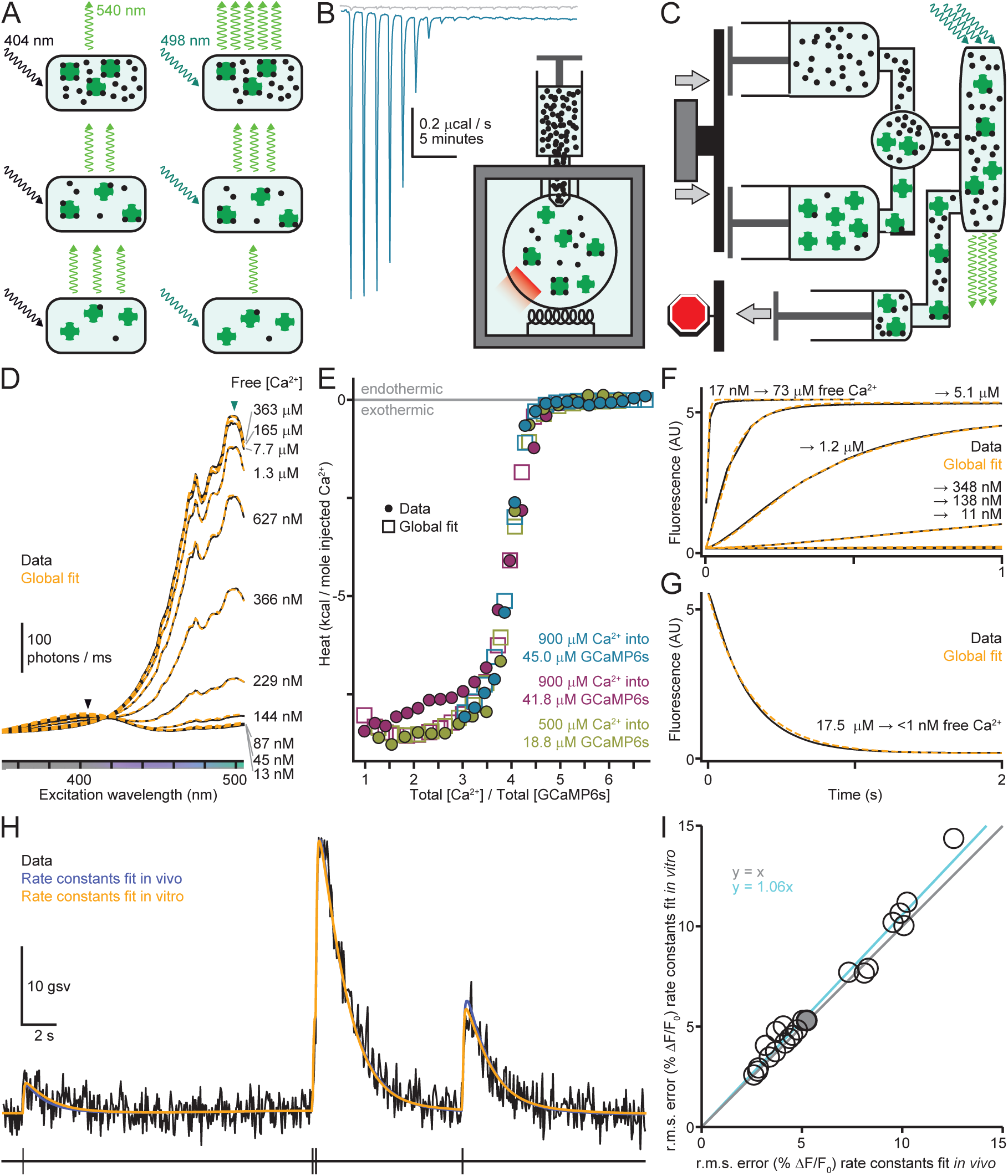
**(A)** Fluorescence spectroscopy of GCaMP6s; fluorescence showed the largest increase with [Ca^2+^] at 498 nm, and the largest decrease at 404 nm. **(B)** Isothermal titration calorimetry experiment, with repeated injection of concentrated Ca^2+^ into GCaMP6s and measurement of the heat absorbed or released upon binding. Raw calorimetric data (turquoise curve) show a net exothermic process for each injection (negative peaks) until Ca^2+^-saturation of GCaMP6s. Heats of dilution for Ca^2+^ (gray) were measured by injecting Ca^2+^ into low-calcium pH buffer (30 mM MOPS with 100 mM Kcl, pH 7.2). **(C)** Stopped-flow fluorescence measurements of GCaMP6s Ca^2+^-binding kinetics. BAPTA-buffered solutions of GCaMP6s and calcium are rapidly mixed and fluorescence is then measured over time. **(D)** Fluorescence excitation spectra of GCaMP6s (black, emission wavelength 540 nm) and global SBM fit (orange) for a range of calcium concentrations and 1.6 mM BAPTA. Arrows indicate 404 and 498 nm. **(E)** Integrated peak heats (circles) for 3 ITC experiments and global fit (squares) vs. the ratio of total Ca^2+^ and GCaMP6s after each injection. **(F)** Kinetics of Ca^2+^ binding to GCaMP6s measured using a stopped-flow device. 0.9 µM GCaMP6s in 37 µM BAPTA was mixed with 1.7 mM BAPTA and total Ca^2+^ranging from 0-1.9 mM. **(G)** Kinetics of calcium release from GCaMP6s. 0.9 µM GCaMP6s with 21.1 µM total Ca^2+^ was mixed with 8mM BAPTA. **(H)** Electrically detected APs (upper) and simultaneously recorded GCaMP6s fluorescence (black, lower) from a L2/3 mouse visual cortical pyramidal neuron, with model fit to *in vivo* data alone (blue) and model with rate constants fit to *in vitro* data and other parameters fit *in vivo* (orange). **(I)** Root-mean-square error for SBM with rate constants fit *in vitro* vs. *in vivo* (n = 22 pyramidal neurons). Unity line is shown in gray and linear regression in cyan.

### AP inference with the SBM

We next applied the SBM to the problem of inferring AP times and neuron-specific parameters ([GCaMP6s]^total^ and *F*_BG_) from fluorescence data alone. To identify the AP-sequence most consistent with the fluorescence data, we developed a sequential Monte Carlo algorithm [49] (SMC, or particle filtering, reviewed in [35]). SMC, a data-driven simulation technique that has also been used with synthetic indicators [131], generates many simulations or particles whose predictions are compared to observed data (Figure 5a). For the SBM each particle consisted of a simulated AP sequence along with time courses for free calcium, GCaMP6s binding states and baseline fluorescence (Figure 5b). The algorithm was advanced forward in time by randomly extending each particle’s AP sequence and baseline fluorescence while solving the rate equation to update all molecular species’ concentrations (Figure 2 - figure supplement 1), using *in vivo*-derived rate constants as these provided slightly tighter fits to the data (Figure 4i). At each fluorescence measurement the particles’ predictions were compared to the observed fluorescence value (Figure 5a) to calculate probability weights (Figure 5c, methods), and particles were then eliminated, retained or copied multiple times with probabilities determined by their weights. Particles whose predictions did not match the data were more likely to be eliminated, so those with incorrect AP sequences disappeared after a small number of measurements. After the SMC algorithm traversed all the fluorescence measurements, a fixed-lag smoother [73] combined the particles and weights at each time point to compute the probability of spiking over time given the fluorescence data. As the number of particles increases, SMC methods converge to unbiased Bayesian estimators of the hidden state variables (APs, binding states and baseline) and of the data’s likelihood given model parameters [27].

**Figure 5:**
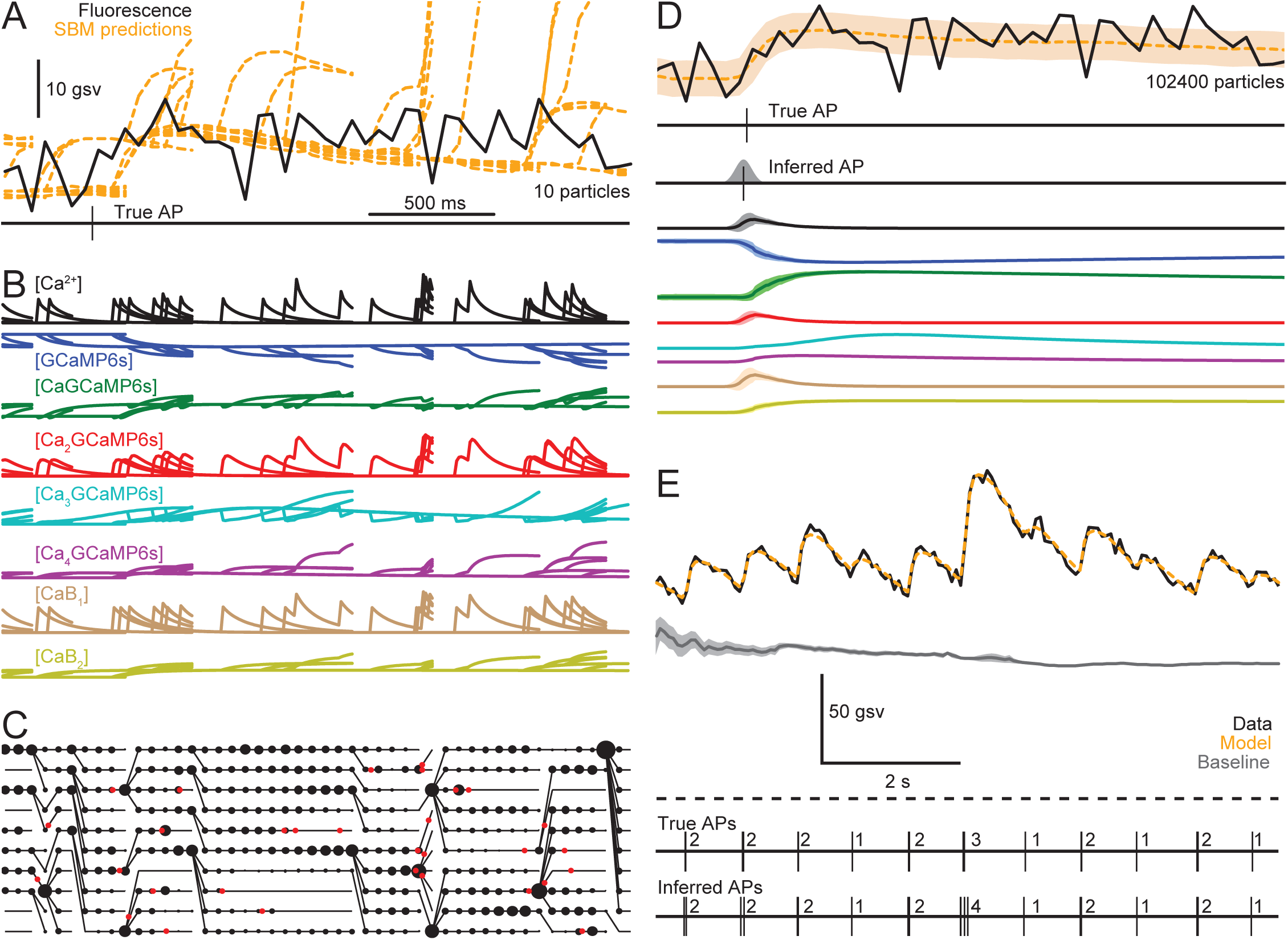
**(A)** GCaMP6s fluorescence (black, upper) from a L2/3 mouse visual cortical pyramidal neuron, with an electrically recorded AP (lower) and predicted fluorescence from ten SBM simulations used in a sequential Monte Carlo (SMC) algorithm (orange). **(B)** Time-varying concentrations of free calcium, GCaMP6s binding states and calcium-bound endogenous buffers for the SBM simulations in (A). Scalebar in (A): 4 µM Ca^2+^, 10 µM GCaMP6s, 10 µM CaGCaMP6s, 2.5 µM Ca_2_GCaMP6s, 20 nM Ca_3_GCaMP6s, 750 nM Ca_4_GCaMP6s, 40 µM CaB_1_ and 15 µM CaB_2_. **(C)** Probability weights (areas of black dots) for the SMC filtering distribution, calculated using the simulations and data in (A). Black lines indicate particle ancestry for the SMC algorithm; red dots indicate AP discharge times in the SBM simulations. **(D)** Inferred AP time with temporal uncertainty (upper) from an SMC algorithm with 102400 particles; same data as in (A-C). Means and standard deviations (lower) are shown for the SMC smoothing distributions of all molecular species’ concentrations given fluorescence data in (A) and 102400 particles, along with the posterior mean and standard deviation of de-noised fluorescence (orange). Data units are scaled as in (A-B). **(E)** SMC/SBM inference of AP times from 9 seconds of data during which fluorescence never decays to baseline (dashed line indicates 0 fluorescence).

Developing an SMC algorithm for the SBM required sev-eral new computational techniques to improve the speed and reliability of AP inference. To deal with the computational burden of solving the SBM’s global rate equation (Figure 2 - figure supplement 1) separately for each particle, we implemented a custom ordinary differential equation solver without loops or branch points to maximize the efficiency of GPU-based multithreaded computation (Figure 5 - figure supplement 1, methods). This solver uses a backward (implicit) Euler method, allowing time steps 1000 times longer than for a forward (explicit) solver while maintaining accuracy and numerical stability. We also designed new techniques for randomly extending particles’ AP sequences (the SMC sampler) and selecting particles at each fluorescence observation (resampler) specifically for SMC-based AP inference (Figure 5 - figure supplement 2, details in methods). Most importantly, our SMC algorithm analyzed the fluorescence data twice, with the results of the first round used to design sampling and resampling distributions for the second round. Finally, we adjusted the fixed-lag smoother delay, which determines how far the algorithm looks into future fluorescence data when inferring APs at each time point (see methods). Observing the standard tradeoff [73] between incorrect results at short delays and inconsistent results at long delays (Figure 5 - figure supplement 3), we set this value to 500 ms. These techniques greatly increased the reliability of the SMC algorithm’s output, but over 10^5^ particles were still required for consistently accurate inference (Figure 5d, Figure 5 - figure supplement 4).

**Table 2.** SBM parameters fit *in vivo*

| Symbol | Description | Best-fit value | Range of fit values* | Units |
| --- | --- | --- | --- | --- |
| $k_1^+$ | First GCaMP6s on-rate | 2.4 | 2.3 - 12752 | $\mu\text{M}^{-1} \text{s}^{-1}$ |
| $k_2^+$ | Second GCaMP6s on-rate | 263 | 2.9 - 413 | $\mu\text{M}^{-1} \text{s}^{-1}$ |
| $k_3^+$ | Third GCaMP6s on-rate | 31 | 2.5 - 44 | $\mu\text{M}^{-1} \text{s}^{-1}$ |
| $k_4^+$ | Fourth GCaMP6s on-rate | 30 | 30 - 43 | $\mu\text{M}^{-1} \text{s}^{-1}$ |
| $k_1^-$ | First GCaMP6s off-rate | 2.2 | 0.3 - 7093 | $\text{s}^{-1}$ |
| $k_2^-$ | Second GCaMP6s off-rate | 390 | 1.5 - 532 | $\text{s}^{-1}$ |
| $k_3^-$ | Third GCaMP6s off-rate | 2.2 | 2.0 - 19 | $\text{s}^{-1}$ |
| $k_4^-$ | Fourth GCaMP6s off-rate | 3.7 | 3.4 - 4.3 | $\text{s}^{-1}$ |

\*Range of parameter values fit from multiple initializations that converged with a normalized r.m.s. residual <10% higher than the lowest r.m.s. residual over all initializations (17 / 50 initializations satisfied this criterion).

In addition to inferring APs and baseline drift, the algorithm also estimated [GCaMP6s]^total^ and *F*_BG_ for each neuron by maximizing the likelihood of fluorescence data given these parameters (Figure 5 - figure supplement 5, methods). After the SMC algorithm returned the probability of an AP every 10 ms, a subsequent processing step produced a single AP sequence without time discretization along with temporal uncertainty for each AP (Figure 5d-e, Figure 5 - figure supplement 6, methods).

When a neuron’s true AP sequence is known, baseline fluorescence can be calculated directly using periods without APs (Figure 3 - figure supplement 1). However, when AP times are unknown or have been held out for testing, baseline and APs must be inferred together based on fluorescence data alone. To test whether an SMC algorithm could cope with data where true baseline fluorescence is never observed due to high firing rates, we analyzed a recording from a pyramidal neuron where rhythmic AP bursts prevented the fluorescence from returning to baseline (Figure 5e) without including periods of inactivity before or after the analyzed data. Despite the lack of fluorescence observations at baseline, the algorithm correctly identified the number of APs in nearly all bursts and inferred a drifting baseline less than the minimum fluorescence value. Having verified that our SBM/SMC algorithm can successfully infer APs in individual cases, we next tested it on our complete *in vivo* dataset.

### Accuracy of AP inference techniques

We evaluated the accuracy of SBM-based AP inference along with alternative methods (MLspike, c2s-s, c2s-t and thr-*σ*), comparing true electrically detected APs to the APs inferred by each method (Figure 6a-c). For correlation-based analyses that do not depend on data units, we also included two linear deconvolution-based methods that do not infer APs, but rather a unitless quantity proportional to the neuron’s firing rate: FOOPSI [132] and CFOOPSI [105]. We tested performance using a cross-validation procedure for all methods and accuracy measures to prevent overfitting: the SBM or other method was fit with each individual neuron excluded from the training data, and accuracy was evaluated using the held out neuron.

**Figure 6:**
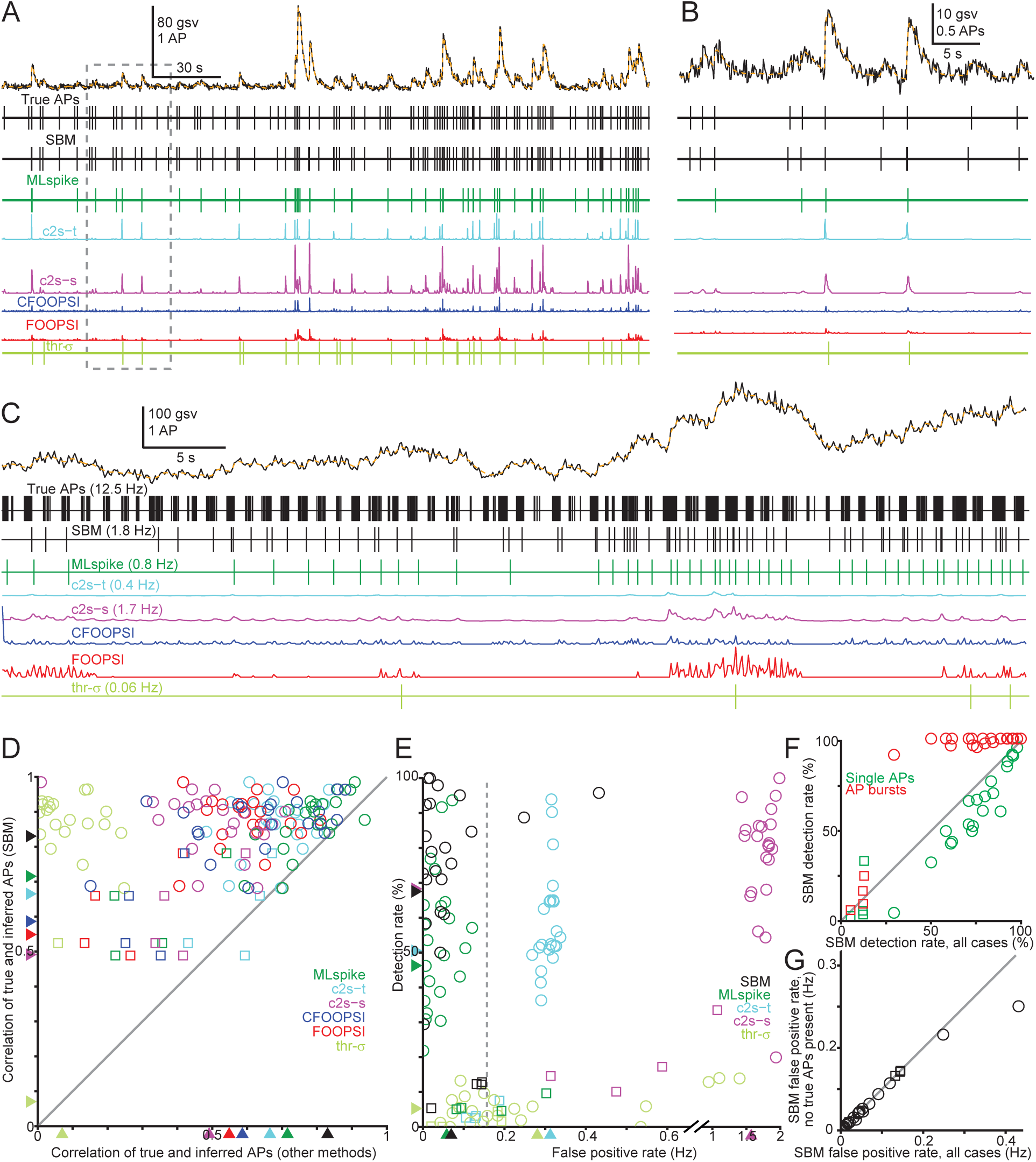
**(A)** Electrically detected APs and GCaMP6s fluorescence (black, upper) from a L2/3 mouse visual cortical pyramidal neuron. Orange curve shows the SBM’s posterior mean for denoised fluorescence. AP inference results (lower). **(B)** Detailed view from (A). **(C)** As in (A-B) for an interneuron neuron firing at 12.5 Hz. **D** Correlation between each neuron’s true and inferred AP sequences for the SBM (y-axis) compared to other methods, with 100 ms Gaussian smoothing (n = 26). Arrowheads indicate mean correlation over neurons for each method and squares indicate interneurons. **(E)** False positive and detection rates for each neuron and algorithm (n = 26). Arrowheads indicate mean over neurons, squares indicate interneurons and the dashed line shows the true mean spontaneous firing rate. **(F)** SBM detection rates for single APs (black) and bursts of 2 or more APs (red, burst duration <100 ms), compared to overall detection rates for each neuron (squares indicate interneurons). **(G)** Rate of false positives inferred by the SBM without any true APs within 200 ms (y-axis) compared to overall false positive rates including overestimation of AP counts (x-axis). Squares indicate interneurons.

We first evaluated the agreement of true and inferred AP sequences by calculating their correlation over time for each algorithm and neuron. We used Gaussian smoothing (*σ* = 100 ms, see methods) to emphasize accuracy of AP counts as opposed to precise AP discharge times within bursts (see below for analysis of timing accuracy). Correlation between the true and inferred AP sequences (Figure 6d) was highest for the SBM (0.83*±* 0.13, n = 26 neurons), with significantly higher mean (p < 5e-6, t-tests) and median correlations (p< 0.005, rank sum tests) than other methods and higher correlation for nearly all neurons (24-26/26, p < 2e-5, sign tests). The SBM also produced the highest correlations for 5 neurons recorded in 3 animals after development of the SBM and all algorithms for model fitting and AP inference (0*±*.77 0.15, next highest c2s-t at 0.63 *±*0.12). The F1-score [29], which like correlation reaches 1 for perfect accuracy but is also sensitive to the units of algorithms’ outputs (replacing every inferred AP with 2 APs does not change the correlation), was also highest for the SBM (0.65 *±* 0.23, Figure 6 - figure supplement 1).

To characterize the agreement of true and inferred AP sequences beyond the single values provided by the correlation and F1 measures, we next computed the rates of missed APs and false positive detections when no APs were present (Figure 6e). For these comparisons we used a timing tolerance of 100 ms when matching the true and inferred AP sequences. The SBM detected 67.5*±* 29.2% of APs (n = 26 neurons), with a false positive rate of 0.07 *±*0.09 Hz. MLspike detected APs at a lower rate of 46.3 *±*24.3% (p < 1e-6, t-test) but its false positive rate was not significantly different from the SBM (0.06*±* 0.06 Hz, p = 0.54). Similarly, c2s-s detected APs at the same rate as the SBM (68.3 *±*26.4%, p = 0.88) but with a false positive rate ten times the true median firing rate (1.57 *±*0.45 Hz), possibly because it was trained on data with firing rates of 2-5 Hz [126]. Compared to the SBM, lower detection and higher false positive rates were observed for c2s-t (50.5 *±*23.8%; 0.31 *±*0.10 Hz) and thr-*σ* (5.3±4.4%; 0.28*±* 0.34 Hz). In 19/26 neurons tested, the SBM detected over half of all APs and less than half its inferred APs were false positives. Of the 7 neurons for which SBM did not satisfy this criterion, 4 were interneurons. In contrast, MLspike satisfied this criterion for 11/26 neurons, and c2s-t, c2s-s and thr-*σ* for no neurons.

We next examined which features of the AP sequence were associated with various types of inference errors for the SBM. About half the APs missed by the SBM (52 *±*32%, n = 26 neurons) were found to be isolated single APs (as opposed to bursts or longer discharge sequences). Overall, *±*58 *±*31% of isolated single APs were detected by the SBM, though this ranged over neurons from 3% to 100% (Figure 6f, green). For isolated bursts of 2 or more APs within 100 ms, the chance of detecting at least 1 AP was *±*99 0% for pyramidal neurons (Figure 6f, red circles, n = 22), and 14 *±*7% for interneurons (Figure 6f, red squares, n = 4). When the SBM inferred false positive APs not present in the electrical recording, in 93 *±*12% of cases no true APs were present within 100 ms, while in the remaining cases APs were present but their number was overestimated (Figure 6g). In summary, false negative errors (missed APs) arose from a combination of failure to detect isolated single APs, underestimation of activity in interneurons and underestimation of AP counts in bursts. False positive errors arose predominantly from spurious detections when no APs were present, with overestimation of AP counts accounting for only a small fraction of these errors.

The SBM’s accuracy depended on the characteristics of signal and noise in each neuron’s fluorescence recordings. Neurons whose single APs evoked larger fluorescence increases exhibited higher correlations between true and SBM-inferred APs (Figure 6 - figure supplement 2, r = 0.42, p = 0.05, t-test, n = 22) and higher detection rates (r = 0.58, p = 0.005) but not lower false positive rates (r = 0.07, p = 0.74). Similarly, signal-to-noise ratio (SNR) was positively associated with correlation (r = 0.36, p = 0.05, Figure 6 - figure supplement3) and detection rate (r = 0.51, p = 0.003) but unrelated to false positive rate (r = −0.04, p = 0.8). Imaging frame rate showed no significant association with correlation, detection rate or false positive rate over the range of 10-60 Hz (p > 0.2).

Finally, we examined how AP inference accuracy depends on the type and quantity of training data used to fit SBM parameters. We first examined the effect of using GCaMP6s rate constants obtained from *in vitro* binding assays, while fitting physiological parameters from *in vivo* data with cross validation. This resulted in only a slight reduction in accuracy (correlation 0.81 *±*0.13, detection rate 65.2*±* 29.4%,false positive rate 0.07 *±*0.06 Hz, n = 26, Figure 6 - figure supplement 4) which did not significantly change correlation (p = 0.39). We also examined how our approach of model fitting followed by SMC-based AP inference would fare in an extremely data-poor scenario, when model parameters are fit from only a single neuron’s data. To compute accuracy across our entire dataset, we carried out AP inference for each neuron based on parameters fit from the previous neuron (Figure 6 - figure supplement 5, methods). Correlation between true and inferred AP sequences was nearly unchanged when fitting SBM parameters to a single neuron’s data, whether using GCaMP6s rate constants fit to *in vivo* data (r = 0.78*±* 0.13 vs. 0.83 *±*0.13, p = 0.20) or those fit to *in vitro* binding assays (0.80 *±*0.13 vs. 0.81*±* 0.13, p = 0.67). A much larger decrease in correlation values was observed when training c2s on a single neuron (r = 0.51 *±*0.18 vs. 0.67 *±*0.10, p = 0.0003); this difference might arise from c2s’ greater numberof parameters (1̃000) which could make it more susceptible to overfitting.

Overall, these results show that the SBM provides a clear advance in accuracy and robustness compared to currently available inference methods, for multiple error metrics and in both data-rich and data-poor conditions. Nonetheless, it is worth emphasizing that for some neurons many APs will be missed by all algorithms, as some single-AP responses are not distinguishable from noise (Figure 6 - figure supplement 6).

### Linear readout of neural activity

We next examined whether APs were accurately inferred at all levels of neural activity, or only for certain firing rates. We first divided our entire *in vivo* dataset, ranging from isolated single APs to high frequency bursts (Figure 7a), into 500 ms windows and calculated the true firing rate in each one (0-24 Hz for pyramidal and 0-48 Hz for interneurons) along with each algorithm’s inferred rate. This analysis revealed a tight linear dependence of the SBM’s average inferred rate on the true rate (Figure 7b) that spanned the full range of firing rates present in our dataset and did not depend on the window size (Figure 7 - figure supplement 1). For the SBM, this linearity was observed in all pyramidal neurons (Figure 7b, black, *r*^2^ = 0.98*±* 0.02, n = 22) and interneurons (Figure 7b, red, *r*^2^ = 0.96*±* 0.03, n = 4), while other methods provided a less linear readout of neural activity (p < 0.01, rank sum tests, Figure 7 - figure supplement 2a-b). The average SBM-inferred rate grew with the true rate at a slope near 1 for pyramidal neurons (0.93 *±*0.20, n = 22), while lesser slopes were observed for other methods (Figure 7 - figure supplement 2c). For all true firing rates, the ratio of the SBM-inferred and true rates was close to 1, but this was not the case for other methods (Figure 7c). All methods tested underestimated firing rates in interneurons (Figure 7d, Figure 7 - figure supplement 2d). The SBM also provided the best agreement between the true and inferred values of neurons’ overall mean spontaneous firing rates, and for pyramidal neurons this relationship was linear with a slope near 1 and y-intercept near 0 (Figure 7 - figure supplement 3). These results show that the SBM linearly reads out pyramidal neurons’ firing rates from <0.1 Hz (mean spontaneous rates) to the maximum rate recorded in our dataset (> 20 Hz over 500 ms windows) and can do so with correct units, while for interneurons firing rate readout is linear but underestimates the true AP count.

**Figure 7:**
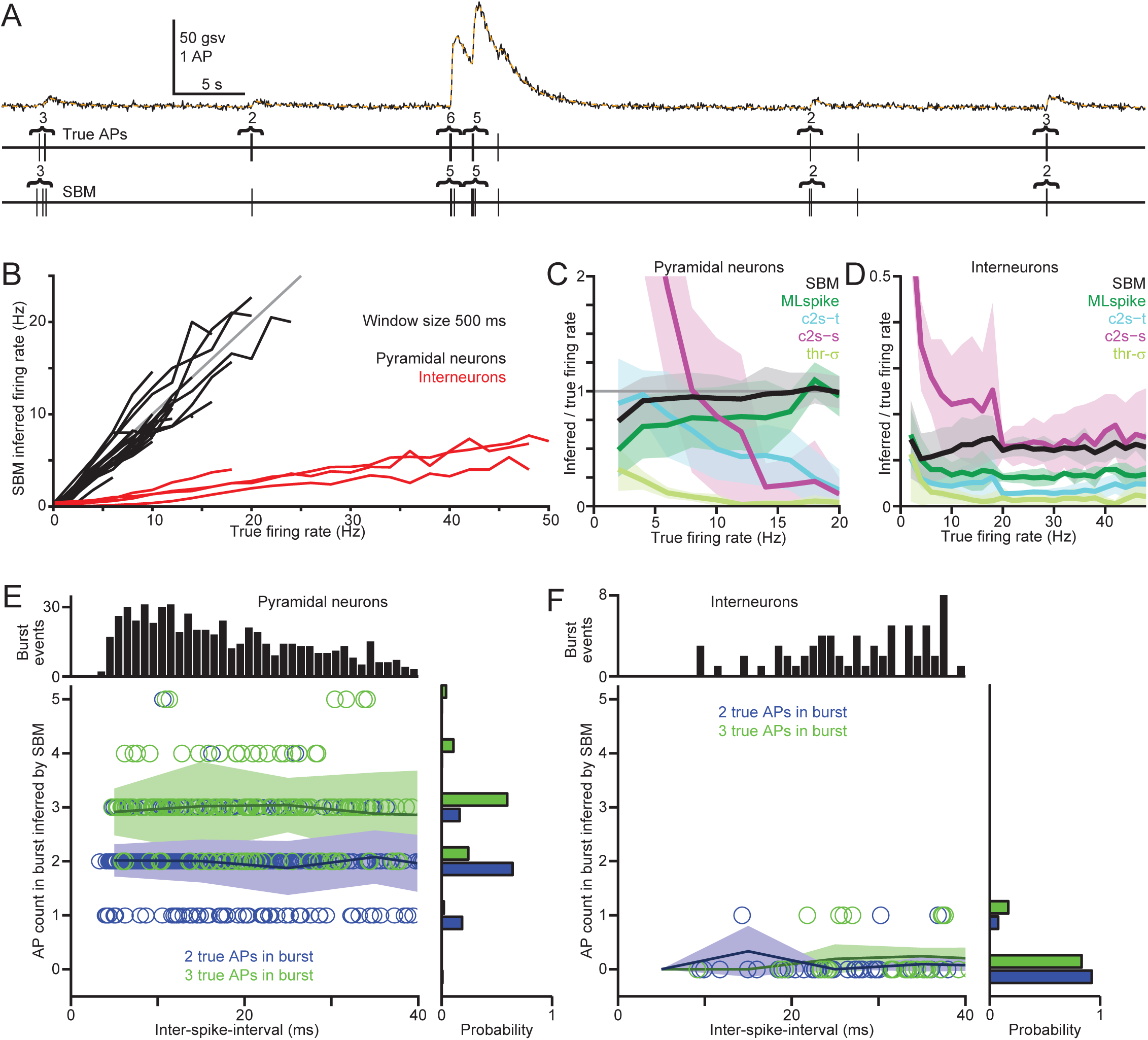
**(A)** Electrically detected APs and GCaMP6s fluorescence (black, upper) from a L2/3 mouse visual cortical pyramidal neuron, and SBM-inferred APs. Orange curve shows the SBM’s posterior mean for denoised fluorescence. Numbers indicate the true and inferred AP counts for each burst; single APs are unlabeled. **(B)** Firing rate inferred by the SBM as a function of the true firing rate, calculated across the complete *in vivo* dataset using 500 ms windows with 90% overlap. Each curve corresponds to a single pyramidal neuron (black) or interneuron (red). **(C)** Ratio of inferred to true firing rates in pyramidal neurons, as a function of the true firing rate, for each inference method. Each curve shows an average over neurons (n = 22) and shaded regions show standard deviations. **(D)** As in (C), but for interneurons. **(E)** *Center*: AP counts inferred by the SBM in pyramidal neurons as a function of inter-spike interval (ISI) for bursts of 2 (blue) and 3 APs (green). Solid lines show average inferred AP counts in 10 ms ISI bins. For 3-AP bursts, the mean of the two ISIs was used. Inferred APs were counted from 200 ms before the first AP in the burst to 200 ms after the final AP, and only bursts flanked by 400 ms without other true APs before and after were included. *Top*: Histogram of ISIs for 2- and 3-AP bursts. *Right*: Probability distributions for the number of SBM-inferred APs for bursts of 2 (blue) and 3 true APs (green). **(F)** As in (E), but for interneurons.

We also examined whether the inter-spike intervals (ISIs) between nearby APs affect the number of APs inferred by the SBM. In recordings from pyramidal neurons, we first identified 2-AP bursts with ISIs up to 40 ms (n = 528, shortest 3.3 ms), and compared the ISI to the number of inferred APs (Figure 7e, blue). Calculating the average inferred AP count as a function of ISI, we found these quantities to be uncorrelated (p = 0.92, 10 ms time bins). Similar results were observed for 3-AP bursts (Figure 7e, green, p = 0.88, n = 211) and for interneurons (Figure 7f, p = 0.96, n = 75 for 2-AP and p = 0.07, n = 59 for 3-AP bursts). As expected from the above analyses of firing rate, the SBM underestimated AP counts in bursts for interneurons, but did so in a uniform way that did not depend on ISIs. These results show that SBM-inferred AP counts do not depend on ISIs, and that our approach can successfully enumerate APs separated by < 10 ms.

### Precision of inferred AP times

When inferring events such as APs from measurements with limited SNR and temporal resolution, event detection times will generally not be precisely correct. Instead, they will exhibit some time difference from the nearest true event time, resulting in a probability distribution of timing errors that depends on the data, model and inference method. Investigating temporal precision is particularly important for AP times inferred from GECI fluorescence signals, as SNR can be <1 and the frame rate as low as 10 Hz (Figure 6 - figure supplement 3). To measure the accuracy of inferred AP times, we first examined how the correlation of true and inferred AP-sequences (Figure 6d) depends on the Gaussian filter size *σ* used to smooth both AP sequences before comparing them (Figure 8a). When the timing error between true and inferred APs is less than *σ*, the Gaussian filtering causes the smoothed AP sequences to overlap and increases correlation, so smaller *σ* values impose a more stringent requirement for accurate AP times. The SBM produced the highest correlations for all values of *σ*. Similarly, we calculated AP detection and false positive rates (Figure 8b-c) as functions of the timing tolerance within which true and inferred APs were considered as matching. For all timing tolerances, detection rates were similar for the SBM and c2s-s and lower for other methods, while false positive rates were similar for the SBM and MLspike and higher for other methods. We also examined cross-correlation as a function of the time lag from true to inferred APs (Figure 8d, Figure 8 - figure supplement 1d, *σ* = 25 ms); the SBM exhibited the highest cross-correlation peak (0.64*±* 0.18, Figure 8e) at the shortest time lag (−2.0 *±*29.7 ms, Figure 8f).

**Figure 8:**
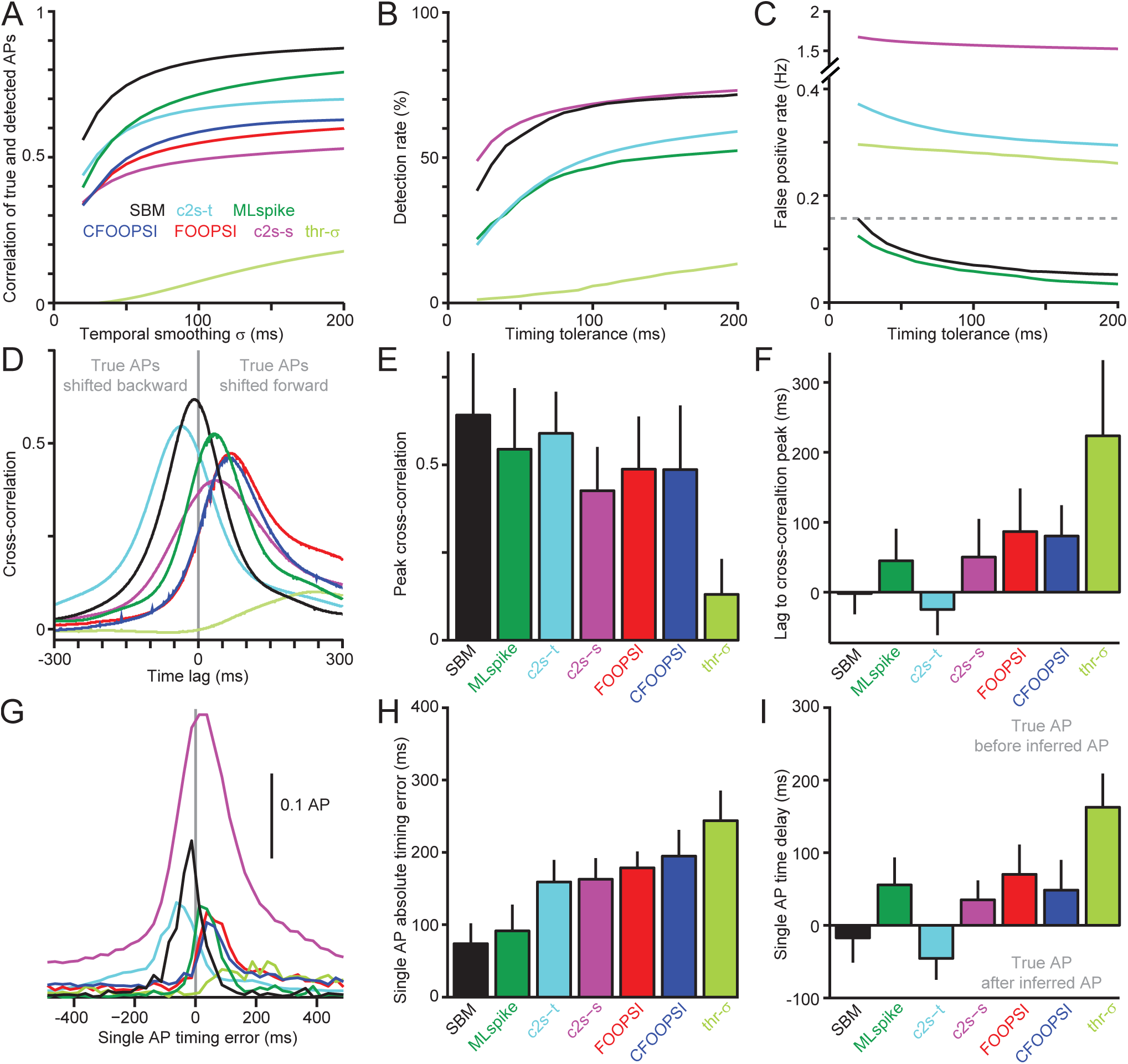
**(A)** Correlation between true and inferred AP sequences as a function of Gaussian smoothing *σ* and averaged over neurons (n = 26). **(B)** Detection rate as a function of timing tolerance. **(C)** False positive rate as a function of timing tolerance; dashed line indicates median spontaneous firing rate. **(D)** Cross-correlation between true and inferred APs for each inference method, as a function of time lag. Positive lags correspond to shifting true APs forward in time before comparing them to inferred APs. Cross-correlations were computed with a smoothing *σ* of 25 ms. **(E)** Mean and standard deviation over neurons of the peak cross-correlation. **(F)** Mean and standard deviation over neurons of the time lag for which cross-correlation was maximized. **(G)** Mean rates of AP inference relative to true AP times (gray vertical line) for single APs without other APs for 1 s before and 0.5 s after. The unitless outputs of CFOOPSI and FOOPSI have been rescaled for each neuron to give an average total output of 1 for each true single AP. **(H)** Mean and standard deviation over neurons of each algorithm’s absolute timing error for isolated single APs. **(I)** Mean and standard deviation over neurons of each algorithm’s average delay from true to inferred AP times for isolated single APs.

To further quantify timing precision, we analyzed the time differences between true and inferred APs for each algorithm. Using only isolated single APs (without other APs 1 s before and 0.5 s after), we calculated for each method the rate of inferred APs (or for FOOPSI and CFOOPSI, the unitless output) as a function of time difference from the true AP (Figure 8g, Figure 8 - figure supplement 1a-c). For each neuron with isolated single APs (n = 22), we then computed the average absolute time difference from the true AP for each algorithm’s output (Figure 8h), which was lowest for the SBM (64 *±*28 ms). The distribution of SBM-inferred AP times peaked 17.4*±* 51.4 ms before the true AP time, while other methods exhibited larger positive or negative delays (Figure 8i).

To examine whether the SBM’s timing errors can be attributed to some aspect of the image acquisition process itself, we also compared the absolute error and mean time delay for single APs to SNR and imaging frame rate (Figure 8 figure supplement 2). Absolute timing error decreased with SNR (r = −0.46, p = 0.015) but not frame rate (r = 0.18, p = 0.38) while the mean time delay was closer to zero at higher frame rates (r = 0.43, p = 0.024) but not at higher SNR (r = −0.20, p = 0.31). The absolute time differences between true and SBM-inferred APs could also be predicted by the timing uncertainty values output by the SBM for each AP (Figure 8 figure supplement 3), showing that the SBM can accurately quantify its own uncertainty regarding AP times.

Because inferred APs can precede true APs in time (Figure 8g), sensory-evoked APs detected optically may violate the assumption that causes precede effects [4, 50]. To illustrate this, we carried out simulations of L2/3 neurons’ sensory responses (Figure 8 - figure supplement 4) which showed that optical AP detection can cause stimulus-evoked APs to be assigned to times before the stimulus. Therefore, in order to facilitate the use of optically detected APs for causal time series analysis we calculated the fraction of inferred APs before the true AP, and how this fraction can be reduced by shifting inferred APs forward in time. For isolated single APs, 90% of SBM-inferred APs can be assigned after the true AP time when using a forward shift of 114 ms, but when the latency of stimulus-evoked APs is known a smaller shift can be used (Figure 8 - figure supplement 5). Together, these results show that the SBM results in smaller timing errors and average time differences from true to inferred APs than other methods, but care must be taken when identifying causal effects from optically measured AP activity.

### Quantifying sensory-evoked AP discharge

Having developed and validated the SBM using spontaneous activity in single neurons, we also applied it to infer sensory-evoked activity across neuronal populations. We recorded GCaMP6s fluorescence in visual cortex while presenting drifting grating stimuli (5 repetitions in 8 directions) and inferred AP times using the SBM, MLspike and thr-*σ* (Figure 9a-b). Sensory stimulation at each neuron’s preferred orientation resulted in SBM-inferred firing rates from 0-7 *Hz (mean*0.9 *±*1.3 Hz, n = 66), while lower firing rates were inferred by MLspike (0.3*±* 0.4 Hz, p = 2e-5, t-test) and thr-*σ* (0.5*±0.2*Hz, p = 0.04), consistent with these algorithms’ lower AP detection rates (Figure 6e, Figure 7c-d). Orientation tuning curves calculated using SBM-inferred APs (Figure 9c, black) were similar to previous observations [93], including a diverse range of preferred orientations as well as untuned and nonresponsive neurons. Some pairs of adjacent neurons had opposite orientation preferences (Figure 9c, neurons **i** and **iv**), as previously observed using synthetic indicators in rats [97] and mice [88]. These results show that the SBM can be used to measure neurons’ sensory tuning, and that quantitative estimation of this tuning can depend on the choice of AP inference method.

**Figure 9:**
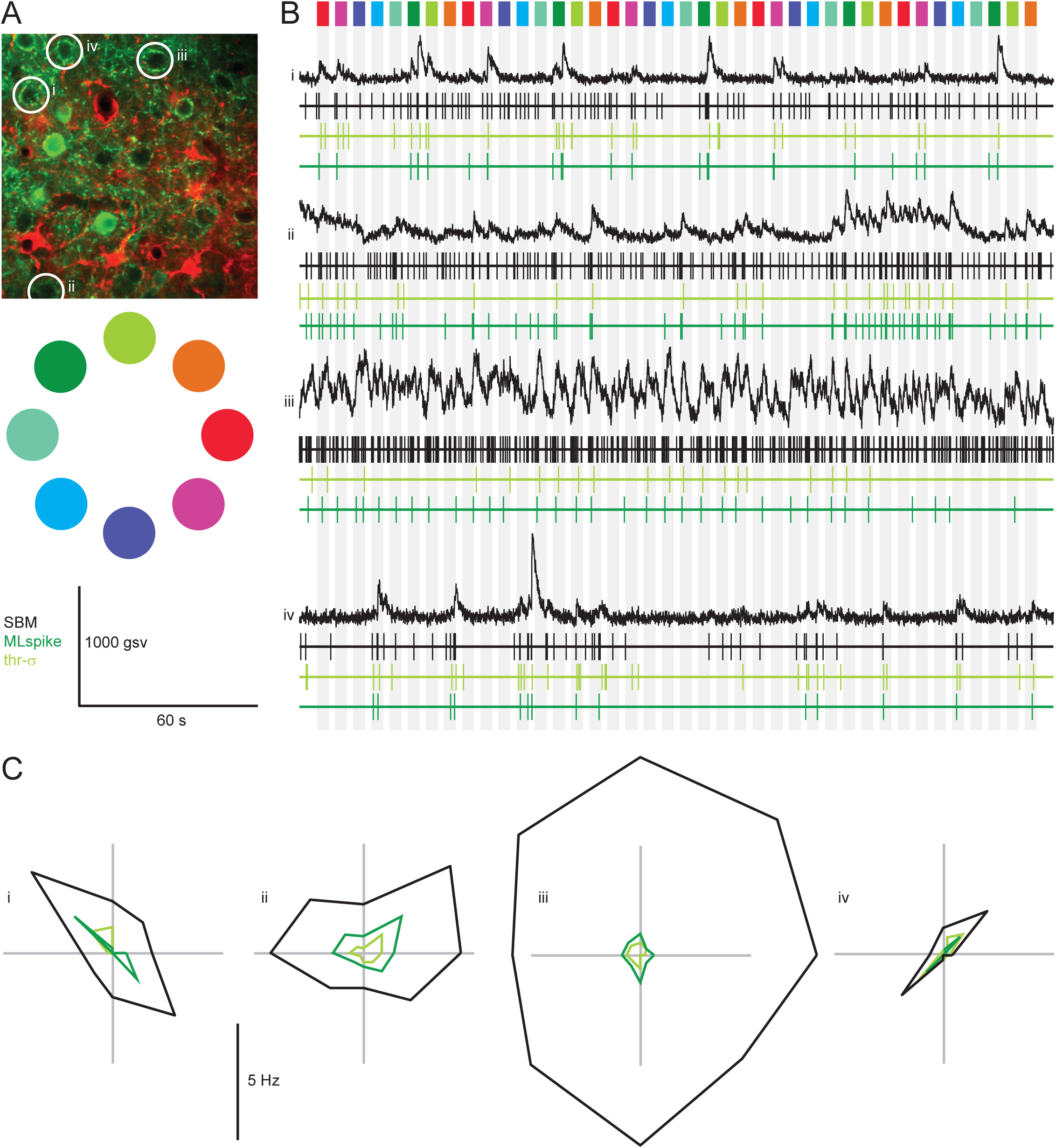
**(A)** Two-photon image of neuronal population (upper) expressing GCaMP6s (green) in L2/3 rat visual cortex, with astrocytes stained using sulforhodamine 101 (red) and imaged during visual stimulation with drifting gratings in 8 directions (colored circles, lower). **B** Fluorescence (black) recorded from 4 neurons from the population in (A) during presentation of drifting grating stimuli (gray bars; colors above indicate the direction of motion for each stimulus presentation). AP inference results are shown for the SBM (black), MLspike (dark green) and thr-*σ* (light green). **C** Orientation tuning curves showing the mean firing rate evoked by presentation of the drifting grating stimulus in 8 directions, for the 4 neurons in (A-B). Firing rates were calculated over the 2-second stimulus presentation.

## Discussion

We developed the SBM to address the complex, nonlinear and variable relationship between AP discharge and GCaMP6s fluorescence. By using quantitative biophysical modeling to link fluorescence and APs, it outperformed existing inference methods (figures 6-8) with more accurate firing rates, higher correlation and F1 values, higher detection rates, fewer false positives, greater linearity and more precise AP times. This improved accuracy can be explained by several differences from previous approaches. For methods based on linear deconvolution (FOOPSI [132] and CFOOPSI [105]) and thresholding (thr-*σ* [32]), accuracy is limited by the fact that GCaMP6s fluorescence neither increases linearly with AP counts nor crosses a fixed threshold upon AP discharge (Figure 1). For MLspike [29], the difference may arise because individual binding steps and endogenous buffers are not included in the phenomenological model, or because MLspike updates its internal state only once per image frame. c2s uses the largest number of parameters (1004) and is the most mathematically flexible method tested, but infers APs based only on fluorescence values within a one second window around each time point while other techniques look further into the past and future. In addition to the specific limitations of each of these methods, they all fail to address the variability of the AP-fluorescence relationship over neurons, which is a central challenge for AP inference from GECI fluorescence. While algorithms such as c2s which fix all parameters after fitting cannot account for this variability, the opposite extreme of estimating all parameters separately for each neuron would lead to overfitting and is likely to be computationally infeasible for models rich enough to describe nonlinear GECIs. A small number of parameters can be chosen to vary across neurons (for example, a single scaling factor as proposed in [29]), but for non-physical parameters such as polynomial coefficients or deep learning connection weights there is no reason for neuron-to-neuron variability to manifest in some parameters and not others. In contrast, the SBM captures this variability using the parameters [GCaMP6s]^total^ and *F*_BG_ (which are expected from their physical interpretations to vary over neurons), requires fewer total parameters than c2s and is more robust when trained on smaller datasets (Figure 6 - figure supplement 5).

The SBM’s false positive rate was 0.07 Hz, bringing GCaMP6s into the range achieved by synthetic sensors such as OGB1 [51, 53, 71, 70]. However, even with the SBM GCaMP6s simply cannot match OGB1’s sensitivity to single APs: excluding the 4 interneurons to match previous OGB1 measurements in pyramidal neurons brings the detection rate to only 78%, compared to 90-95% for OGB1[51, 53, 71, 70]. This may reflect an upper performance limit intrinsic to GCaMP6s that cannot be overcome by further improvements of the SBM or other methods, as visual inspection shows a total lack of fluorescence increase for some single APs (Figure 6 - figure supplement 6). Consequently, while bursts of 2 or more APs were nearly always detected, for many neurons isolated single APs were frequently missed (Figure 6f). These errors could complicate efforts to understand how neuronal populations carry out their essential functions, as even small numbers of APs in a single neuron can have strong effects on neuronal tuning through synaptic plasticity [102]. On the other hand, the fact that most of the SBM’s false positive errors occurred in the absence of true APs (Figure 6g) suggests that most subsequent analyses would be unaffected by these errors.

The SBM provides the most linear readout of neural activity (Figure 7a-d, Figure 7 - figure supplements 1-3), with the ratio of true and inferred firing rates independent of the true rate (Figure 7c-d) and of ISIs within bursts (Figure 7e-f). This is an important characteristic for an AP inference algorithm, since it facilitates subsequent stages of quantitative analysis such as computation of sensory tuning curves [92]. Furthermore, for pyramidal neurons the SBM allows comparison of firing rates across neurons with correct units (Figure 7b-c, Figure 7 - figure supplement 2c), despite the variation in AP-evoked fluorescence across neurons (Figure 1c, Figure 3e-f).

Compared to alternative methods, SBM-based inference introduced both lesser absolute timing errors and smaller mean delays between true and inferred APs. AP timing plays a central role in dendritic integration [13], sensory coding [26, 67, 85, 135] and both *in vitro* [11, 81] *and in vivo* synaptic plasticity [86, 102]. Nonetheless, for GCaMP6s SBM-based inference produced timing errors far larger than the timescale of synaptic input integration [134] and often larger than the image acquisition time. Since timing errors were reduced in recordings with higher SNR and mean delays were reduced at higher frame rates (Figure 8 - figure supplement 2), timing precision can be modestly improved by imaging faster or reducing noise, but major gains may require switching to a faster sensor. Because the timing uncertainty arising from optical measurement of APs (Figure 8d-i) can reverse the apparent order of an inferred AP and the sensory stimulus that evoked it (Figure 8 - figure supplement 4), causal analysis of optically measured neural activity should either take the uncertainty of inferred AP times into account or apply a corrective time shift (Figure 8 - figure supplement 5) before further analysis.

A current limitation of our SBM/SMC approach is that it underestimates activity for interneurons (figures 6-7). This can be attributed to interneurons’ negligible fluorescence increases for single APs (Figure 1 - figure supplement 3), as previously reported using OGB1 with post-hoc immunohistochemistry [69]. We emphasize that this shortcoming was specific to interneurons rather than high firing rates, as the algorithm accurately inferred APs in pyramidal neurons at firing rates >20 Hz and ISIs < 10 ms (Figure 7). Furthermore, the SBM’s inferred firing rate grew linearly with the true rate in interneurons from 0 to >40 Hz, albeit with a slope <1 (Figure 7b,d). These results could be explained by stronger calcium buffering in interneurons, in which case SBM performance could be improved by relaxing the assumption that the same endogenous buffers are present in all neurons. Including per-neuron buffering parameters would require solving a more difficult parameter estimation problem for each neuron, possibly using recent machine learning techniques for parametric model fitting [78, 101, 128, 14]. More generally, the SBM could benefit from more extensive quantitative knowledge of endogenous buffers, extrusion mechanisms and calcium flow between the cytosol, nucleus and endoplasmic reticulum [110, 112, 115]. The SBM remains extremely simplistic in comparison to the diverse set of calcium-binding molecules present in neurons; for example, the InterPro database lists 826 different mouse proteins containing the calcium-binding EF hand domain pair alone [42].

SBM-based AP inference could also benefit from new methods for acquiring and pre-processing imaging data. New scanning techniques can increase imaging speed and the number of recorded neurons [20, 53, 103] towards the levels achieved by electrical approaches. Reduction in neuron-to-neuron variability might be achieved by using transgenic animals instead of viral delivery [24, 79] or by localizing the indicator to subcellular compartments [36, 72]. Accuracy could also benefit from improved methods for motion correction [32, 52, 105] and removal of contaminating signals from nearby structures [31, 89, 105].

Current knowledge and experimental tools provide only a partial picture of the calcium dynamics evoked by AP discharge in neuronal somata, but the SBM is broadly corroborated by past experimental measurements. The value of Δ[Ca^2+^]_AP_ = 20.2 µM calcium influx per AP is consistent with measurements in midbrain neurons [107] using the action-potential clamp method [77] that reported 4.32 pC of charge carried by calcium ions for each AP. For a spherical soma 10-15µm in diameter, this would imply a concentration increase of 13-43 µM. While this far exceeds the sub-micromolar changes in free calcium typically observed upon AP discharge [116, 142], the SBM models most calcium ions inside the neuron as bound to endogenous buffers. For every free calcium ion, the SBM modelled 203 ions bound to endogenous buffers in a neuron at rest, consistent with previously reported values for pyramidal neurons *in vitro* [60, 80, 99].

Because the SBM describes the effect of AP discharge on fluorescence through the biophysical framework of mass action kinetics, we were able to apply it to *in vitro* binding assays as well as *in vivo* experiments. SBM parameters determined from global fits of multiple *in vitro* experiments can be used *in vivo*, both to predict fluorescence from APs (Figure 5) and to infer APs from fluorescence (Figure 6 - figure supplement 4). The fact the SBM can describe the same protein-ligand interaction under such divergent circumstances lends support to the model class as a quantitative account of calcium-GCaMP6s interactions. However, for both *in vitro* and *in vivo* data, individual parameters could not be precisely determined, though predictions of the system’s output were reliable and robust (Figure 3 - figure supplement 9). This parameter sloppiness could arise from spectroscopically silent binding steps (including the first 3 steps when using the best-fit SBM parameters for *in vivo* data, see Table 1), insufficient temporal resolution to resolve the most rapid binding reactions, insufficient number or diversity of experimental conditions *in vitro*, the limited number of recorded neurons or diversity of AP sequences *in vivo* or lack of structural identifiability in the model itself [8]. This sort of parameter sloppiness has been observed in many other complex models in biochemistry and systems biology [45, 54, 108].

The SBM and associated techniques for model fitting and AP inference have a number of applications beyond detecting APs with GCaMP6s. These procedures could be applied without modification to GCaMP6f/m [18], other GCaMPs, recently developed red fluorescent versions [23] and future calmodulin-based sensors. Because the SBM describes both the calcium sensor protein and some aspects of neuronal physiology, it could also prove useful in investigating the physiological side-effects of GCaMP expression in virally transfected neurons [2, 139] and genetically modified animal lines [119]. Given that SBM parameters obtained *in vitro* can be used to predict fluorescence and infer APs *in vivo*, SBM parameters fit from *in vitro* binding assays might be useful in large-scale screening of candidate GECIs [18]. The model could also be extended to include photoconversion of calmodulin-based activity integrators [44], multiple protein products of the same GECI gene [23] and calcium dynamics in dendrites, axons, spines and boutons [21, 39, 75, 115, 121]. As a framework for linking APs to experimental observations through reaction kinetics, more general versions of the SBM could describe biochemical reaction networks coupled to neural activity [10]. The models, fitting procedures and software tools we developed for *in vitro* binding assays could also be applied to other calcium-binding proteins or different protein-ligand interactions, as existing analysis software cannot easily fit model parameters for multiple binding sites or combine equilibrium and kinetic data from multiple binding assays in a common global fit [141].

The approach we have taken here, of fitting parametric generative models to experimental data while incorporating domain-specific knowledge into design of the model class, has already proved useful in several areas of neuroscience [37, 58, 66, 43, 109, 95, 100] and other fields of biology [45, 54, 94, 96, 108]. By improving the accuracy and interpretability of data analysis, these results help justify the use of quantitative causal models and domain-specific knowledge over model-free black box methods for the analysis of complex biological data.

## Methods and Materials

### Mathematical notation

*N*(*µ, σ*^2^) denotes a Gaussian distribution with mean *µ* and variance *σ*^2^, while *N*(*x*; *µ, σ*^2^) denotes the pdf of this distribution at the value *x*. We define a sum indexed by the empty set as zero, and a product indexed by the empty set as one. The operator denotes convolution. For a vector *v ϵ* ℝ^*n*^, diag(*v*) is an *n* x *n* matrix with *v* on the diagonal and zeros elsewhere. For a vector *xϵ*ℝ^*n*^, we denote by *x*_*j*:*k*_ the sub-vector starting at *x*_*j*_ and finishing with *x*_*k*_.

### Virus injection

Experiments were conducted according to German animal welfare regulations. Experimental subjects for our combined optical-electrical recordings were 6 male C57BL6 mice, 20-25g body weight at the time of the virus injection. Fluorescence imaging with visual stimulation was carried out 70g male Listar hooded rat. During all surgical and recording procedures, anesthetic depth was regularly monitored and body temperature was maintained at 37°C using a heating pad and thermal probe. Mice were anesthetized with an intraperitoneal bolus injection of ketamine and xylazine (120 mg/kg and 16 mg/kg respectively) while rats were anesthetized with fentanyl citrate, midazolam hydrochloride, and medetomidine hydrochloride (5 µg/kg, 2 mg/kg and 150 µg/kg respectively). Supplemental doses of anesthetic solution given as required. The target area for the virus injection (Lambda −1 mm, lateral 2.5 mm in mouse and Lambda +1.5 mm, lateral 4.5 mm in rat) was exposed and marked. A small craniotomy (∼0.5 x 0.5 mm) was opened approx. 1 mm anterior to the marked target area at the same lateral coordinate and a small opening made in the underlying dura. To induce expression of GCaMP6s in cortical neurons, a glass pipette (tip opening ∼20 µm) filled with virus solution (AAV1.Syn.GCaMP6s.WPRE.SV40, Penn State Vector Core, PA, USA) was advanced into the cortex to the target area at a 15-20° angle to the brain surface, and an injection was made over approximately minutes (250-400 nL in mouse and 1.5 µL in rat). The surgical site was then protected with silicone (KwikSil, World Precision Instruments, FL, USA) and skin incision closed. Animals were given a bolus dose (5 mg/kg flunixin meglumin for mice; 120 µg)/kg naloxone hydrochloride, 200 µg)/kg flumazenil, 750 µg)/kg atipamezole) for post-operative analgesia, and allowed to recover. Additional, buprenorphine hydrochloride and carprofen (30µg)/kg and 5 mg/kg respectively) were administered for the rat.

### Imaging with simultaneous electrophysiological recording

After an expression time of 10-15 days, animals were anesthetized with urethane (1.6 mg/kg), a custom-made headplate fixed to the skull using dental cement (Paladur, Kulzer GmbH, Hanau, Germany), an approx. 3 x 3 mm craniotomy centered over the injection target site opened and the dura removed. Astrocytes were counterstained using sulforhodamine 101 (0.5 mM, Sigma-Aldrich, MO, USA), which was applied topically to the cortical surface for 60-120 s, and the exposed cortex then covered with agar (1.2%, Sigma-Aldrich, MO, USA, dissolved in artificial cerebrospinal fluid (ACSF) of the following composition in mM: NaCl, 135; KCl, 5.4; CaCl2, 1.8; MgCl2, 1; Hepes, 5) and a coverslip to minimize brain movement during subsequent multiphoton and electrophysiological recordings.

Labeled neurons and astrocytes were visualized using custom-built multiphoton microscopes. Excitation light was provided by a Ti:Sapphire pulsed laser system (Mai Tai, Spectra Physics, CA, USA) tuned to 920 nm. Datasets were acquired using either an Olympus 20x (XLUMPlanFl, Olympus, Tokyo, Japan) or a Zeiss 20x (WPlan-APOCHROMAT, Zeiss, Oberkochen, Germany) objective lens. The scanning system consisted either of a conventional or resonance galvanometric system. With the conventional galvanometric system, imaging was carries out at 64 x 128 pixel resolution, acquired at frame rates of 10.4, 15.6, 18.6 or 37.2 Hz. With the resonance galvanometric system datasets comprised either 512×512, or 1024 x 256 pixel images, acquired frame rates of 30 and 60 Hz respectively. Motion within each frame of the imaging datasets was corrected using the method of [52] using red fluorescence from sulforhodamine 101.

Cell-attached electrophysiological recordings were made using glass pipettes (6-12 M? resistance) filled with ACSF containing either 2.5 µM Alexa 594 (ThermoFisher, MA, USA) or 2.5 µM Alexa 488 (ThermoFisher, MA, USA). Electrophysiological signals were amplified using a Multiclamp 700B (Molecular Devices, CA, USA), equipped with a CV 7B headstage. Signals were lowpass filtered at 10 kHz (Bessel filter) and digitized at 20 kHz (Power 1401, Cambridge Electronic Design, Cambridge, UK). Cells were visually targeted under multiphoton guidance for acquiring simultaneous imaging and cell-attached datasets.

### Imaging with simultaneous visual stimulation

After an expression time of 14 days, the rat was anesthetized with Ketamine-Medetomidine (100 mg/kg and 2 mg/kg respectively) with supplemental doses of anesthetic solution given as required. A custom-made headplate fixed to the skull, a craniotomy was opened and the dura was removed with the same procedure as for simultaenous optical/electrical recordings. Visual stimuli were generated using a custom-written Matlab script and the Psychophysics Toolbox, and presented with a monitor (Faytech, 16 x 12 cm, 60 Hz refresh rate) placed 15 cm in front of the rat, covering approximately 56° x 42° of the visual field. Full-length drifting bar gratings (0.05 cycle/°, 2 cycle/s, 8 orientations, 2 s duration, 3 s inter-stimulus-interval, 100% contrast) were binocularly presented with 5 repetitions. Two-photon imaging with another custom-built microscopy visualized labelled neurons and astrocytes similarly as described above, but apart from using an objective lens (Throl Optics).

### Expression and purification of GCaMP6s

GCaMP6s was expressed in E. coli BL21 (DE3) pLysS (Novagen) in LB medium supplemented with 180 µg/mL Ampicillin and 20 µg/mL Chloramphenicol for 24h at 25°C in the absence of IPTG. The pellet was resuspended in 30 mM Hepes, pH 7.2, 500 mM NaCl, 10 mM Imidazole, 1 mM PMSF, and a Roche cOmplete tablet (EDTA-free), lysed by sonication and cell debris was removed by ultracentrifugation. The cleared lysate was incubated with Ni-NTA Superflow beads (Qiagen). After washing with high-salt buffer (1000 mM NaCl), bound protein was eluted in a step gradient with elution buffer (30 mM Hepes, pH 7.8, 500 mM NaCl, and 500 mM imidazole). The pool was subjected to gel filtration (Superdex 200, GE Healthcare) in 30 mM MOPS, pH 7.2, 150 mM KCl.

### Preparation of low calcium buffer (LCB)

All plastic ware and stirring bars were incubated for 30 min in 20 mM Tris (pH 8) with 50 µM EGTA, then washed 15 times with clean water to remove all traces of EGTA. Standard buffer for the measurements was 30 mM MOPS and 100 mM KCl, prepared with trace select water (Fluka), the pH was adjusted to 7.2 (final) using KOH. The buffer solutions were incubated with BAPTA-based polystyrene beads (BAPTA-sponge S, ThermoFisher) on a rotating plate for 24-36 hours at RT to reduce background calcium levels. Calcium contamination was determined using Fura-2 Thermo Fisher, F-1200) by fluorescence spectroscopy.

### Calcium removal by refolding or dialysis

To obtain nearly calcium-free protein levels (calcium-free GCaMP6s) we unfolded GCaMP6s in denaturation buffer (6 M Guanidinium-HCl in 30 mM MOPS, pH 7.2, 100 mM KCl, 2 mM EGTA) followed by repeated concentration and dilution in denaturation buffer using AMICON Ultra centrifugal filters (Millipore). The protein was refolded by fast dilution into 30 mM MOPS, pH 7.2, 100 mM KCl, 2 mM EGTA, 0.5 M Arginine, and 1 M Guanidinium-HCl. The buffer was then exchanged to LCB using AMICON devices. Alternatively, calcium was removed by dialysis instead of refolding. The protein pool was dialysed in slide-a-lyzer buttons (2 K MWCO) for 48 hours at RT and protected from light, against 30 mM MOPS, 100 mM KCl, 10 mM EGTA, 10 mM EDTA, pH 7.5. The buffer was exchanged to LCB by multiple rounds of dilution and concentration at 4°C using 15 mL Amicon concentration devices (10 K MWCO). During calcium removal and in all subsequent steps, we exclusively employed chemicals and water (Fluka 95305) of the highest available grade and pre-cleaned all plastic ware.

### Fluorescence spectra

GCaMP6s was diluted to a nominal concentration of 2 µM in LCB with 2 mM BAPTA and 0 to 2 mM total calcium (final concentrations taking into account the reagent purity values determined during data fitting are shown in the main and supplementary figures). Spectra were recorded at pH 7.2 at 37 C on a PTI spectrometer in photon counting mode (at photon counts below 700000, to avoid nonlinearities in the photon counting module). Excitation wavelength was varied from 350 to 505 nm. The colored spectrum shown in Figure 4d was taken from Wikimedia commons^1^ and is available under the creative commons license.

### Isothermal titrational calorimetry

ITC measurements were carried out in an ITC200 micro-calorimeter (MicroCal, Malvern Instruments) at 37C. The cell and Hamilton syringe were incubated for 1 hour with 20 mM Tris/HCl (pH 8) and 50 µM EDTA, washed twice with the same solution, then washed twenty times with trace select water (Sigma Aldrich). Finally, the cell and syringe were washed four times with LCB.

All solutions were prepared in LCB (pH 7.2) using non-autoclaved Eppendorf tubes and pipette tips. Protein concentrations were calculated from absorption at 280 nm.

Nominal concentrations of 400 to 1200 µM CaCl 2 was titrated into 20 to 60 µM calcium-free GCaMP6s. The experiments consisted of 18 to 30 injections with volumes between 2 and 1.2 µL, 90 s spacing, and constant stirring at 1000 rpm. The initial active cell volume was 203.4 µL. The heat of a 0.2 µL dummy injection at the beginning of each titration was not analysed but its volume and calcium content was taken into account. Baseline correction and peak integration were carried out using [68]. The integrated heat signal was corrected for the heat of dilution (obtained from reference titrations of syringe component into buffer).

### Stopped-flow fluorimetry

Measurements were carried out in an SX20 stopped-flow device (Applied Photophysics) at 37°C in LCB (pH 7.2). The injection volume was set to 145 µL for each syringe. Calcium release from GCaMP6s was measured by mixing 1 µM GCaMP6s and 20 µM total CaCl_2_ in LCB (syringe 1) with 10 mM BAPTA in LCB (syringe 2).

In order to measure calcium binding to GCaMP6s, 1 µM protein in the presence of 50 µM BAPTA was mixed with a defined free calcium concentration (2 mM BAPTA and 0-1.8 mM total calcium) in LCB. These concentrations are nominal values, whereas corrected concentrations taking into account reagent purity values were calculated during data fitting and are reported in the main text and figures.

The instrument was incubated in 50 µM EGTA in LCB over-night before experiments. Syringes were washed using trace select water four times prior to the experiment, then an additional two times using low calcium buffer. Fluorescence was excited at 488 nm (bandwidth 0.5 nm) and a 515 nm cut-off filter was used for the emission. The digitization rate was adjusted to match the speed of the observed kinetics (80 kHz for the fastest trace).

### Extraction of fluorescence signals from *in vivo* imaging data

We used an feature extraction method based on nonnegative matrix factorization [76] to separate fluorescence signals from overlapping structures, similar to previously published techniques [105, 31, 89]. The aim of this method was not to detect fluorescent structures such as neurons and dendrites, but rather to isolate the signals from a single structure that was already identified and marked with a region of interest (ROI). In-depth exploration of the capabilities and limitations of this approach as compared to alternative methods are beyond the scope of the present work and will be described elsewhere (Voit et al., in final preparation).

### Biophysical model

The sequential binding model is a generative, parametric biophysical model designed to describe GCaMP6s and other GECIs both *in vitro* and *in vivo*. Here we first describe the model of reaction kinetics that make up the core of the SBM, while the specific ways in which various data types depend on these kinetics are described in later sections. For a GECI G with *m* calcium binding sites and total concentration [G]^total^, we model the sequential binding steps:

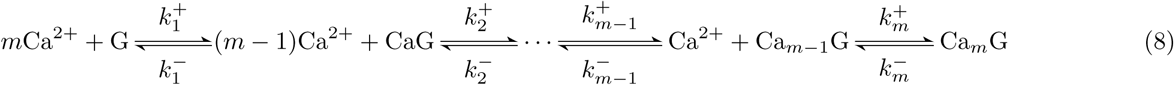

where 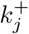 and 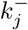 are the kinetic rate constants for binding and unbinding of the *m*-th calcium ion. The net rate of each reaction in the forward direction is then

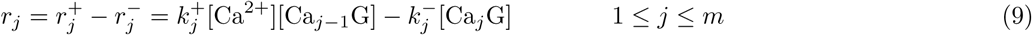

The SBM also includes *n*_buffers_ additional molecules B_1_, B_2_, …, B_*nbuffers*_ binding a single calcium ion, with rates constants 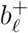 and 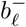 and total concentrations 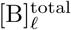. Finally, free calcium is extruded from the neuron at the rate *r*_ex_, which for the standard SBM is

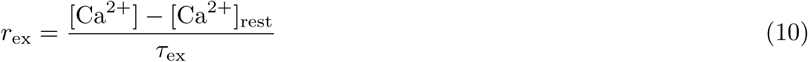

We also implemented a more complicated extrusion mechanism (Figure 3 - figure supplements 3-4), in which *n*_ex_ different extrusion reactions proceed with Michaelis-Menten kinetics. In this case, extrusion reaction *p* saturates at a maximum rate of 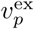 and the dependence on free calcium concentration is described by a Michaelis constant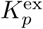, yielding the total extrusion rate

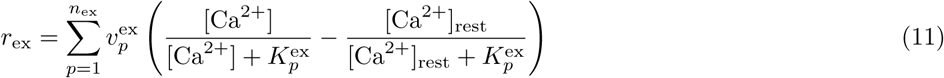

For both extrusion mechanisms, *r*_ex_ = 0 when [Ca^2+^] = [Ca^2+^]_rest_. When analyzing data from *in vitro* binding assays, we set *r*_ex_ = 0.

Together, these reactions define the rate equation for our model [87, 137], a system of coupled nonlinear first-order ordinary differential equations.

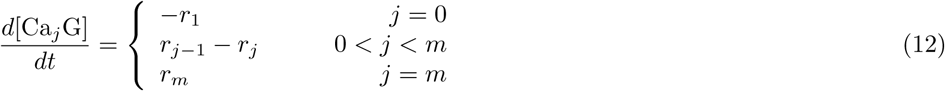

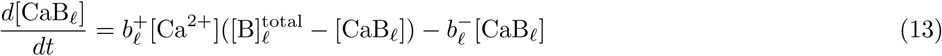

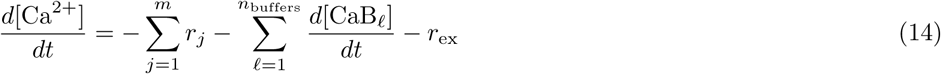

This system of *m* + *n*_buffers_ + 2 equations is slightly redundant, since 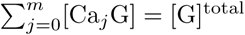 is fixed. However, we explicitly calculated all *m* + 1 indicator binding state concentrations due to the nature of our numerical ODE solver (see below).

#### Equilibrium state for known free calcium concentration

For a fixed calcium concentration [Ca^2+^], we calculate the equilibrium concentrations for all molecular species by setting derivatives to zero and solving the resulting system of linear equations. This gives for the GECI *G* the relations:

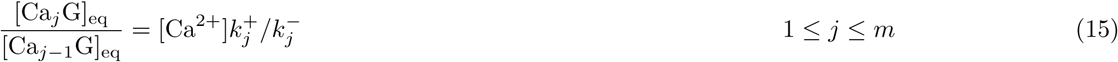

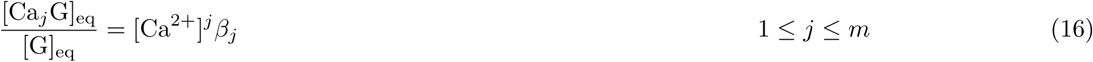

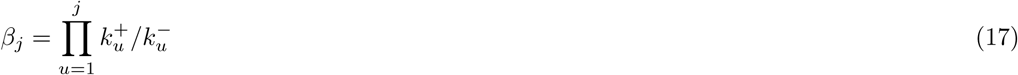

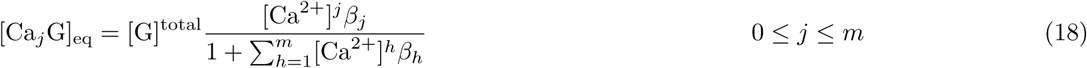

Eq. 18 is known as the Adair-Klotz equation [87, 137] and the *β*’s are macroscopic association constants. Similarly, for the buffers

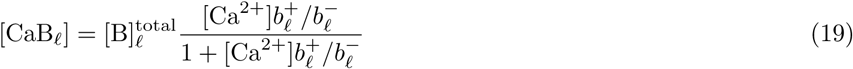

#### Equilibrium state for known total calcium concentration

For certain *in vitro* experiments, the total concentration of calcium [Ca^2+^]_total_ is known but the free concentration [Ca^2+^] at equilibrium is not. In this case we have the relation:

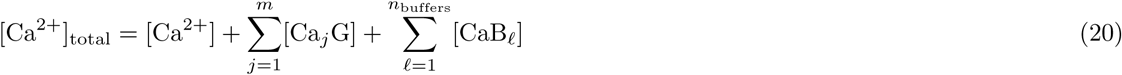

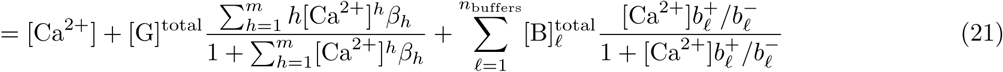

Multiplying by the denominators gives a polynomial equation for [Ca^2+^] of degree *m* + *n*_buffers_ + 1. Because free and total calcium concentrations are strictly increasing functions of each other, this polynomial must posses exactly one real root between zero and [Ca^2+^]_total_, which we obtain numerically using the NumPy function **roots**. Substituting the value of [Ca^2+^]_rest_ obtained from eq. 21 into eq. 18-19 gives the concentrations of all other molecular species.

##### Differentiability of the equilibrium state

Our procedures for fitting the SBM to *in vitro* data require calculating the derivatives of equilibrium binding state concentrations with respect to model parameters. For this purpose, we can obtain the derivative of [Ca^2+^] with respect to any total reagent concentrations or rate constants by implicit differentiation with respect to polynomial coefficients. Suppose that *x*_*R*_ is a root of some polynomial *P* (*x*) with degree *d*. Then considering *x*_*R*_ as a function of the polynomial coefficients, we have for the *j*^th^ order coefficient *p*_*j*_:

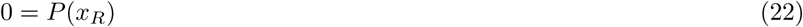

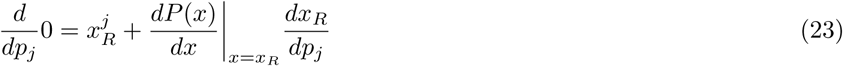

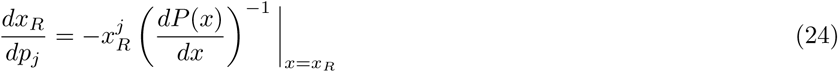

The second line follows from the product and chain rules. Note when the *x*_*R*_ represents the free calcium concentration being calculated from total reagent concentrations, the polynomial derivative 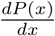 is zero only when *x*_*R*_ = 0, i.e. when both free and total calcium concentrations are zero.

#### *In vitro* data model

We analyzed data from three *in vitro* binding assays: fluorescence spectra, isothermal titration calorimetry (ITC) and stopped-flow fluorimetry.

For spectral and stopped-flow data, the calcium buffer BAPTA [129] was used to control calcium concentration. Using the Maxchelator program [9], we calculated an initial value for the dissociation constant *K*_*d*_ = *b*^*−*^*/b*^+^ = 222.8 nM at 37 °C, ionic strength 0.15 and pH 7.2. We constrained the *K*_*d*_ in subsequent fitting procedures to lie within 5% of this value. While BAPTA’s kinetics have not been as thoroughly investigated as its affinity, [91] reported an off-rate of *b*^*−*^ = 79.0 s^−1^ at 22 °C. We therefore initialized *b*^*−*^ = 90 s^−1^ for data at 37 °C and constrained 1*/b*^*−*^ to lie within 50% to 150% of its initial value of 11.1 ms.

##### Effective concentration and calcium contamination of GCaMP6s and BAPTA

All *in vitro* experiments involved solutions containing the calcium sensor GCaMP6s, calcium ions and in some cases the calcium buffer BAPTA. For each of these, nominal concentration values were calculated based on dilution factors, molecular weight (for BAPTA) and UV absorption at 280 nm (for GCaMP6s). However, true concentrations can differ substantially from nominal concentrations, due imperfect theoretical predictions of protein extinction coefficients [47], increase in the mass of salts through absorption of water and variations in purity across protein purifications or batches of chemical reagents [74, 82, 9, 98]. The effects of calcium contamination can also be significant, especially when considering calcium concentrations approaching the extremely low levels (< 100 nM) present in neurons at rest [116, 60, 80, 142], which we aimed to reproduce in the *in vitro* binding assays.

Therefore, for each pool of purified GCaMP6s or batch of BAPTA, we modeled the ratio of true to nominal concentration as a free parameter, which we refer to as the “purity” of the reagent. We also included for each GCaMP6s pool or BAPTA batch a “calcium contamination” parameter, defined as the ratio of excess total calcium concentration to the reagent’s nominal concentration. We also included a different GCaMP6s contamination value for each ITC experiment, since the absence of BAPTA makes these experiments extremely sensitive to the handling of the protein. We did not include a purity value for the calcium added intentionally from diluted 0.1 M stock, since its concentration errors are likely to be minuscule and since rescaling all concentrations and making a compensatory change to other parameters would not change the data values predicted by the model. Thus our standard calcium stock acts as a unit of concentration or “Urmeter” for our model fits.

##### Data model for fluorescence spectra

Spectral data were represented as a matrix *F* of fluorescence values, where *F*_*λ,v*_ indicates the fluorescence detected with excitation wavelength *λ* in experimental condition *v*. For each spectroscopy experiment, the experimental conditions differed only in the amount of total calcium present. Based on the nominal reagent concentrations, contaminations and purities, we calculated the true concentrations of GCaMP6s, BAPTA and total calcium for each condition. We next calculated free calcium concentration from eq. 21 and all binding state concentrations from eq.18. Denoting the calculated concentration of binding state *i* in condition *v* by [Ca_*i*_GCaMP6s]_*v*_, we modeled fluorescence according to

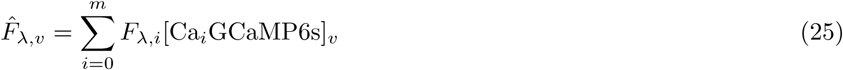

*F*_*λ,i*_ > 0 denotes the fluorescence of binding state *i* when excited by wavelength *λ*.

##### Data model for isothermal titration calorimetry

Data for a single ITC experiment were represented as a vector *Q ϵ ℝ* ^*nQ*^ of integrated peak heats. We first calculated the true concentrations of calcium and GCaMP6s for the syringe and the intial solution in the reaction chamber given the nominal concentrations, purities and contaminations. For each injection, we then updated the concentrations using the standard perfusion model [38]. Specifically, for a reagent *X* withtotal concentration [*X*]_0_ in the cell (after the dummy injection) and total concentration [*X*]_*S*_ in the syringe, the concentration in the cell after the *i*^th^ injecion will be:

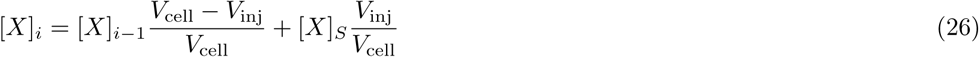

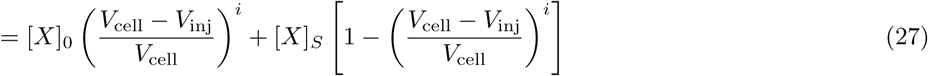

After calculating total concentrations of GCaMP6s and calcium at each stage of the ITC experiment, we calculated free calcium concentration from eq. 21 and all binding state concentrations from eq. 18. We next calculated for each mixture the total enthalpy difference from the calcium-free state arising from binding of calcium to the indicator:

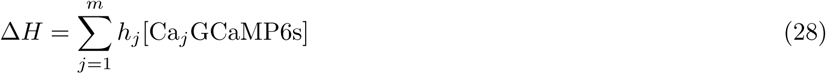

where *h*_*j*_ is the enthalpy difference between one mole of indicator with *j* calcium ions bound and one mole of calcium-free indicator. Finally, denoting the enthalpy difference after *i* injections by Δ*H*_*i*_, we model the integrated peak heat by

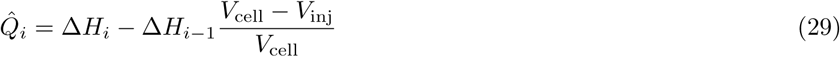

The modeled peak heat 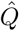 does not contain a term for the pre-reaction enthalpy of each injection, since the syringe did not contain GCaMP6s. Note that we do not explicitly consider heats of dilution, as the heat of dilution for the injection of concentrated calcium was already subtracted, and the dilution heat of GCaMP6s in the reaction cell is negligible compared to binding enthalpy.

##### Data model for stopped-flow fluorimetry

Data for a single stopped-flow experiment were represented as a vector *F* of fluorescence observations *F*_*i*_ at a sequence of times after mixing. For each syringe, we calculated free calcium concentration from eq. 21 and all binding state concentrations from eq. 18, giving the initial equilibrium states before mixing. We then calculated the non-equilibrium binding state concentrations immediately after mixing as the average of the two equilibrium states (since the syringe volumes are equal). We next integrated the rate equation (12-14) from time zero to *t*_dead_, representing the “dead time” delay between initial combination of the two solutions in the mixing chamber and observations made on solution in the fluorescence cell. We then calculated the model’s prediction of time-dependent fluorescence by

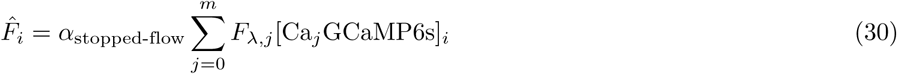

This formula differs from eq. 25 only in the presence of an additional scaling factor, which captures the differences between stopped-flow and spectral data in collection efficiency, excitation intensity, data units, etc. We then integrated the rate equation until the next fluorescence observation and repeated the procedure until the last data point.

#### *In vivo* data model

##### Time step

For imaging data with a time step of Δ between image frames, the SBM used a time step of *δ <* Δ to describe the AP sequence and all concentrations over time. Except where otherwise noted, we chose *δ* to be the largest possible value less than 10 ms for which Δ*/δ* is an integer.

##### AP discharge

Each neuron’s AP sequence was modeled as an independent Bernoulli variable for each time step. The probability of AP discharge was

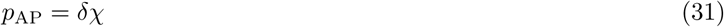

where *χ* is the neuron’s firing rate. The prior distribution of firing rate over neurons was modeled as a Gamma distribution *χ ∼* Γ(*k*_*χ*_*, θ*_*χ*_), where *k*_*χ*_ is a shape parameter and *θ*_*χ*_ is a scale parameter. We modeled each AP as an instantaneous increase in [Ca^2+^] by Δ[Ca^2+^]_AP_.

##### Fluorescence model

Fluorescent measurements are acquired as the neuron’s soma is scanned by the laser focus of the two-photon microscope. The infrared laser light excites the fluorophore leading to emission of visible photons, some of which are captured by the objective lens and detected by the photomultiplier tube (PMT). We model the expected value of the *i*^th^ fluorescence measurement by:

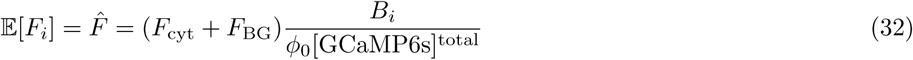

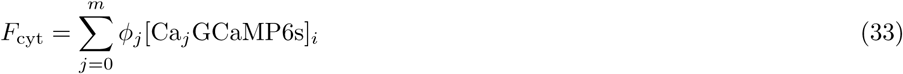

Neither the values of *ϕ*_*j*_ nor their ratios need be consistent with the values of *F*_*λ,j*_ for *in vitro* data, due to the difference between one- and two-photon excitation. *F*_*BG*_ represents fluorescence from out-of-focus structures that does not depend on binding state concentrations in the neuron. *B*_*i*_ represents many factors influencing the scale of the fluorescence measurement, some of which may vary over time. These include laser power, light scattering and absorption by tissue, changes in blood flow through vessels above the image plane, photobleaching of the fluorophore, collection efficiency, artifacts arising from uncorrected lateral or axial motion, PMT quantum efficiency, amplifier gain and the analog-to-digital (A2D) converter’s input range and bit depth. This formulation does not include an additive offset in the fluorescence values, and thus assumes that the recorded fluorescence values will on average be zero in the absence of incoming photons. This can be achieved by subtracting from the fluorescence data an offset recorded with the laser shutter closed before or after each imaging file, a standard feature in many imaging systems such as Scanimage [106].

To ensure that *B*_*i*_ is positive while allowing it to drift over time, we define *B*_*i*_ = *e*^*ρ*^ and modeled *ρ* as a Brownian motion with variance 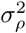 sec^−1^. That is, given the value of *ρ* at time *t*_1_ seconds, its value at a later time *t*_2_ has conditional distribution

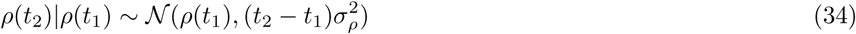

We set *σ*_*ρ*_ = 0.01 for the analysis of all *in vivo* data.

Finally, we define 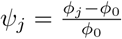 to give a parameterization that does not depend on the units of fluorescence or concentration:

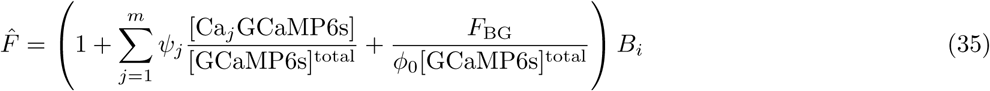

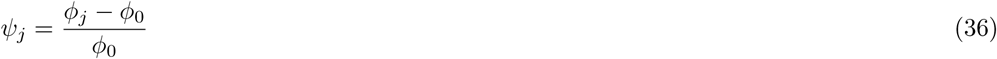

We assume that calcium binding does not decrease the indicator’s brightness, so each *ψ*_*j*_ > 0.

##### Joint prior distribution for [GCaMP6s]^total^ and *F*_BG_

Since resting calcium concentration is consistently 50-80 nM [116, 60, 80, 142], neurons with higher GECI concentrations would be expected to have higher brightness relative to the background fluorescence at rest. We therefore modeled the joint distribution over neurons on [GCaMP6s]^total^ and 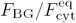 using a 2D log-normal prior:

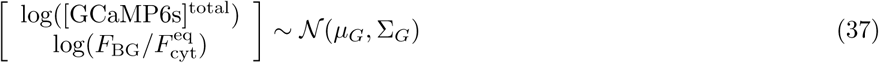

where 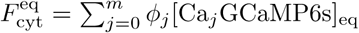, with equilibrium concentrations calculated for [Ca^2+^] = [Ca^2+^]_rest_. Note that [Ca^2+^]_rest_ is not an estimated parameter, but has been fixed to 50 nM (see results).

##### Fluorescence observation noise

For an expected fluorescence value 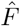 predicted by the SBM, we modeled the probability distribution on fluorescence observations as

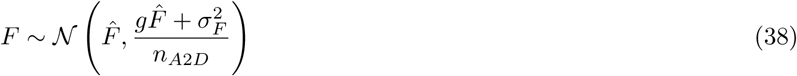

Where *g* is an unknown gain factor used to incorporate photon shot noise and other signal-dependent noise, *n*_A2D_ is the number of analog-to-digital values averaged when calculating the neuron’s fluorescence for each image frame and *σ*_*F*_ represents signal-independent noise. We implemented our model fitting and AP inference algorithms with the option to incorporate calibrated values of *g* and *σ*_*F*_, but when estimating noise properties from fluorescence data alone (see below), we set *g* = 0 and estimated only *σ*_*F*_. Thus for SBM-based AP inference in this study, *n*_A2D_ is known, *g* = 0 and a different value of *σ*_*F*_ is estimated for each neuron based on its fluorescence signals.

### Numerical solution of ODEs

Model fitting and AP inference with the SBM require integrating the rate equation (12-14) forward in time to calculate predicted fluorescence given an AP sequence. When fitting models to *in vitro* data, we used a standard ODE solver available as SciPy library function. We chose the LSODA solver [63] based on the backward differentiation formulas [6] with adaptive step size determination, since we did not know the time constants of the various binding steps in advance and since the stopped-flow data achieved measurement rates as high as 80 kHz. Another reason to use adaptive step sizes is that *in vitro* experiments can produce much larger and more rapid changes in free calcium than those observed in neurons.

However, fitting *in vivo* data required us to design and use a custom solver. This is because when we use our model to detect APs in a sequential Monte Carlo framework (see below), we will be carrying out many simulations in parallel using a GPU implementation. In order to run efficiently on a GPU, these simulations must maintain a low memory footprint per thread, must not allocate and deallocate memory and must avoid branch points such as “IF statements” (to avoid desynchronizing parallel simulations and causing the threads in non-active branches to wait). We therefore designed a custom ODE solver for use with AP inference on the GPU. We also used our custom ODE solver when fitting SBM parameters to *in vivo* fluorescence signals together with known AP sequences, both for speed and to ensure that any approximations or errors would be apparent in the model fits.

#### Backward Euler method

Let *y ϵ* ℝ ^8^ be the vector of binding state concentrations:

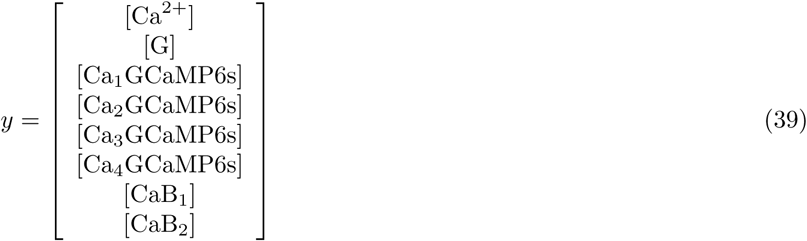

We designed our solver to use the backward Euler method, defined by:

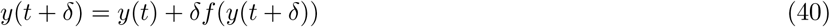

where *f* (*y*) is the vector of derivatives 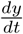, in our case defined by the rate equation (12-14). The backward Euler method possesses several important stability properties [16], allowing us to use a relatively large step size of 10 ms without strongly affecting the fluorescence predicted by the model (Figure 5 - figure supplement 1).

Applying eq. 40 requires solving a system of nonlinear equations, for which we use Newton’s method, an iterative technique based on linearizing *f* using its Jacobian *J*. Given a current estimate *y*^*i*^(*t* + *δ*) of *y*(*t* + *δ*), we approximate

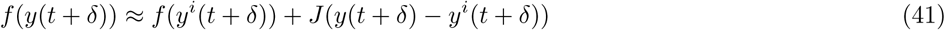

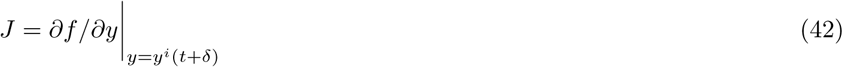

We then use this approximation to repeatedly refine the estimate of *y*(*t* + *δ*) by solving a system of linear equations. Initializing with *y*^0^(*t* + *δ*) = *y*(*t*), we have:

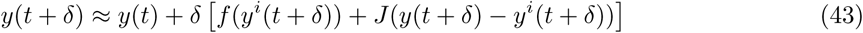

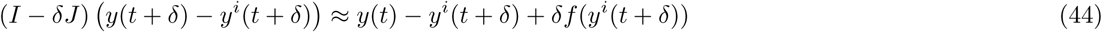

Setting Δ*y* = *y*(*t* + *δ*) − *y^i^*(*t* + *δ*) and *z* = *y*(*t*) − *y^i^*(*t* + *δ*) + *δf*(*y^i^*(*t* + *δ*)) gives:

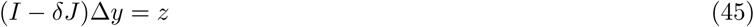

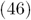

solving for Δ*y* then allows us to compute the next iterative estimate *y*^*i*+1^(*t* + *δ*) = *y*^*i*^(*t* + *δ*) + Δ*y*.

In order to solve this linear system with a low memory footprint and no branch points, we introduce a direct solver that generalizes the tridiagonal matrix algorithm (see below). To avoid additional branch points we also fixed the step size to 10 ms and the number of Newton iterations to 3. We show that this does not lead to inaccurate predictions of *in vivo* GCaMP6s fluorescence even at high firing rates (Figure 5 - figure supplement 1), though a shorter step size or a greater number of Newton iterations might be needed for larger calcium fluxes or faster sensors.

#### Jacobian

To apply the backward Euler method, we use the following Jacobian matrix:

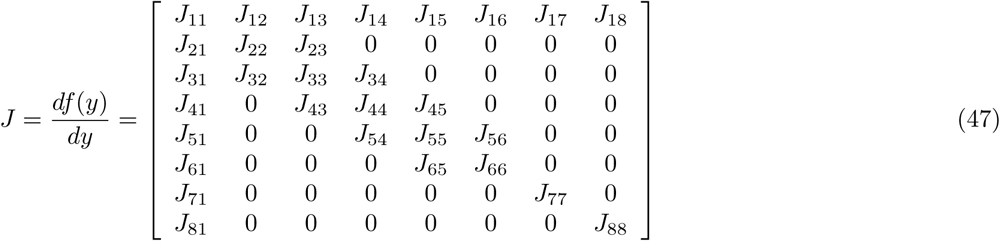

where the elements of *J* are defined as follows for 0 ≤ *j* ≤ 4 and ℓ ≤ *l* ≤ 2:

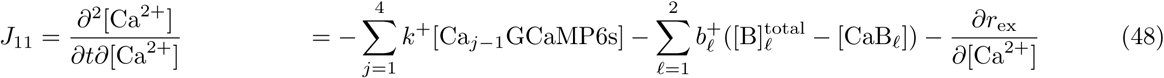

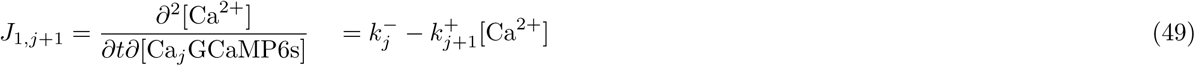

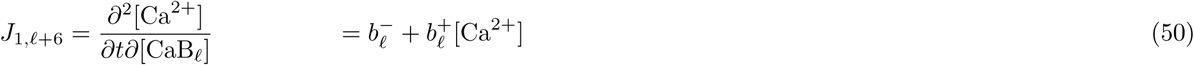

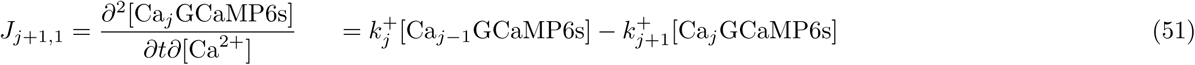

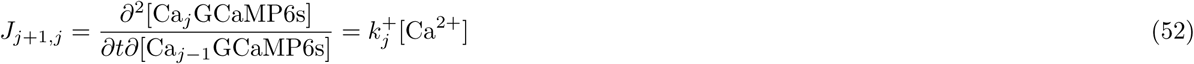

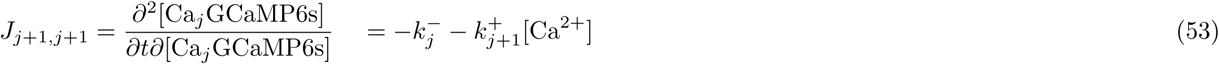

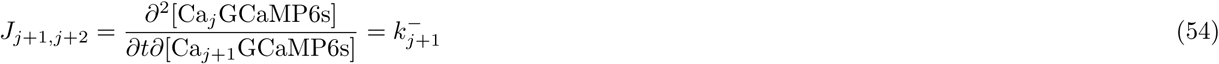

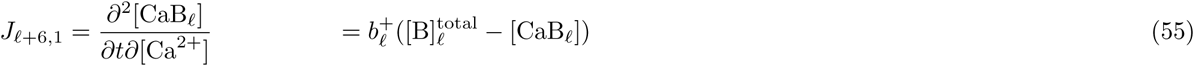

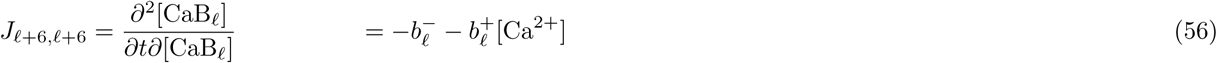

with the convention that with the convention that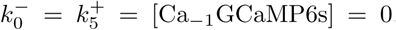. For the standard SBM 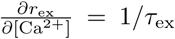, while with Michaelis-Menten extrusion 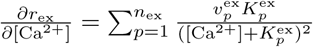.

#### Generalized tridiagonal solver

Except for its first row and column, *J* is a tridiagonal matrix. Here we describe an *O*(*n*) direct method (algorithm 1) for solving such a system of linear equations, generalizing the tridiagonal algorithm [25] to the present case. We consider the (*n* + 1)-dimensional linear system

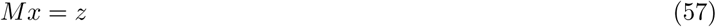

where

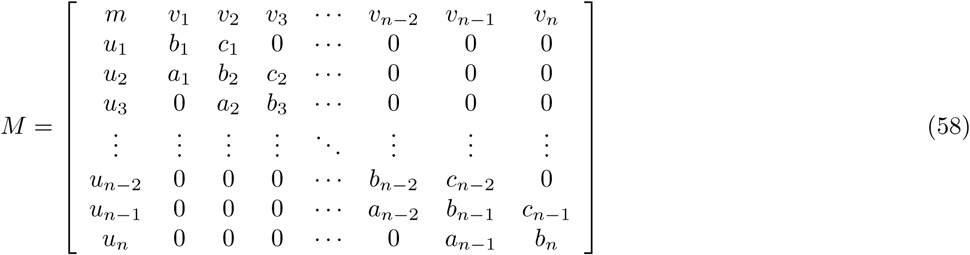

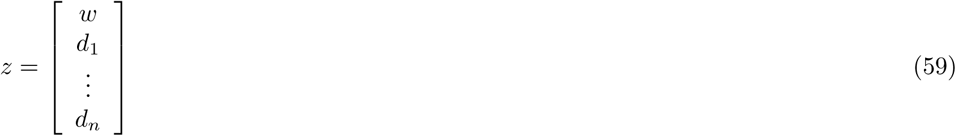

Following the original tridiagonal algorithm, we make a forward pass that sets *a* to zero and *b* to one, then a backward pass that sets *c* to zero. During the backward pass, we also set *v* to zero, and during both passes we keep track of the resulting changes to *u*, *v*, *w*, *m* and *d*. Finally, we make another forward pass that sets *u* to zero while changing only *d*, at which point we have reduced *M* to the identity matrix so that we can return the modified *z* as the desired solution *x*. Elements set to zero or one are not actually modified in memory since they will not be accessed again.

##### Algorithm 1 Generalized tridiagonal solver

~~~
*c* _*1*_ *← c 1 /b* _*1*_
*d* _*1*_ *← d 1 /b* _*1*_
*u* _*1*_ *← u 1 /b* _*1*_
**for** *i = 2 to n* **do**
    *p ← b* _*i*_ *← a* _*i*_ *c* _*i*_*←1*
    *d* _*i*_ *← (d* _*i*_ *← a* _*i*_ *d* _*i*_*←1)/p*
    *u* _*i*_ *← (u* _*i*_ *← a* _*i*_ *u* _*i*_*←1)/p*
**if** *i < n* **then**
    *c* _*i*_ *← c* _*i*_ */p*
  **end if**
**end for**
**for** *i = n ← 1 to 0* **do**
  **if** *i > 0* **then**
    *d* _*i*_ *← d* _*i*_ *← c* _*i*_ *d* _*i*_*+1*
    *u* _*i*_ *← u* _*i*_ *← c* _*i*_ *u* _*i*_*+1*
  **end if**
    *m ← m ← v* _*i*_*+1 u* _*i*_*+1*
    *w ← w ← v* _*i*_*+1 d* _*i*_*+1*
**end for**
*w ← w/m*
**for** *i = 1 to n* **do**
    *d* _*i*_ *← d* _*i*_ *← wu* _*i*_
**end for**
**return**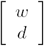
~~~

In our implementations we unroll the loops and use different variable names for each vector element, so that no array indexing or branch statements remain. We also skip the absent off-diagonal elements for endogenous buffers.

### Fitting SBM parameters to *in vitro* binding assay data

We developed procedures to perform global fitting of all *in vitro* parameters and implemented them in the Python programming language. The rate equation and its Jacobian with respect to all molecular species’ concentrations were implemented in Python and NumPy, then compiled using Numba for speed.

#### Objective function

We minimized the weighted sum of squares

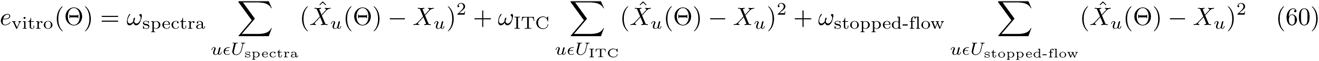

Θ is a vector of SBM parameters, u is an index over all observed data values *X* _*u*_ *, U* _*d*_ is the set of data values belonging to each data modality and *ω*_*d*_ is a weighting factor for data modality. The *ω*’s were chosen so that each data type would make approximately the same contribution to the objective function despite different data units, noise levels and numbers of observations. We first fit data of each type (spectra, ITC and stopped-flow) independently to determine the r.m.s. residual *σ*_*d*_ for each. We then normalized each data type by the corresponding squared residual, and further normalized by the number of observation for each data type:

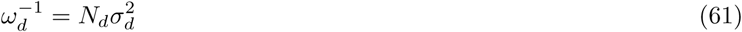

where *N*_*d*_ is the number of data values for data type *d*. Thus our objective function is not affected by rescaling the data units or by adding multiple copies of the data for one data type. It also weights noisier measurements less heavily when fitting the parameters.

#### Gradients

Gradients were calculated in closed form for ITC and spectral data, and using finite differences for stopped-flow data. Finite differences were calculated using **SciPy**’s **approx derivative** function, using the default step size and central (3-point) differences. For parameter values at the upper or lower bounds, **approx derivative** calculates derivatives using the tangent to a 3-point quadratic fit.

#### Profiling over coefficients

For all three types of data, the value predicted by the model can be written as a matrix product of regressors and coefficients:

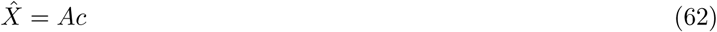

where rows of *X* are data points. For spectral and stopped-flow data, each row of *A* contains the sensor’s binding state concentrations for one data point, whereas for ITC data each row contains concentration changes after each injection for binding states with at least one calcium ion bound. For stopped-flow and ITC data, *c* is a vector of fluorescence values or enthalpy differences for each binding state, whereas for spectral data *c* is a matrix whose rows contain the excitation spectra of the binding states. Note that for clarity in eq. 62 we have ignored the weights *ω* but the results generalize in a trivial way to the case where weights are present.

Instead of directly optimizing over all model parameters, we can first calculate *A* as a nonlinear function of the rate constants, reagent purities, etc. and then optimize ĉ = *A*^*T*^ *A −*1 *A*^*T*^ *X* by linear regression. In the case where bounds on *c* prevent us from using a simple linear regression, we use **SciPy**’s **optimize.nnls** function (where only a positivity constraint exists) or **optimize.lsq linear** (for arbitrary upper and lower bounds). In either case, we obtain the global optimum on *c* since we are minimizing a convex function on a convex set. This allows us to greatly reduce the number of free parameters, especially for spectral data with many excitation wavelengths. It also speeds convergence by avoiding small alternating updates of *c* and *A*.

Let Θ = [Θ_*A*_ Θ_*c*_]^*T*^, where *A* depends only on Θ_*A*_ and Θ_*c*_ is simply *c* in vector form. As long as the columns of *A* are linearly independent, *ĉ* is well defined and we have

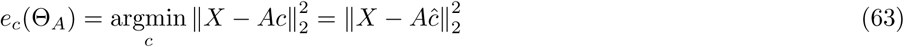

and

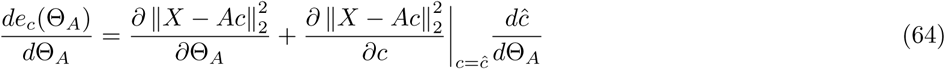

By definition, each element of 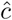 is either at a local optimum or at a bound. For an element 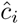 at an optimum, 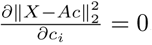 Elements at a bound and not at an optimum will not change upon an infinitesmal change in Θ_*A*_, so for these elements 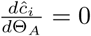. Consequently, the second term on the right hand side of eq. 64 is a dot product whose additive sub-terms are all zero. Thus, after optimizing with respect to *c* we can calculate the gradients of the objective function with respect to Θ_*A*_ while treating 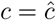 as a constant.

Using eq. 64 does not merely speed up the calculation of gradients. When using an iterative solver (see below), the current gradient and approximation to the Hessian matrix are used to compute a search direction, which is used as the basis for a line search. However, by optimizing out the coefficients we can make larger steps in Θ_*A*_ since *c* will be automatically adjusted during the line search to minimize the objective function.

#### Reparameterization

We used an alternative parameterization for the kinetic properties of GCaMP6s that possessed two advantages over on- and off-rates. First, it allowed us to separate equilibrium and kinetic responses into separate groups of parameters, so that fits to spectral and ITC data would depend on a smaller number of parameters. Second, since the transitions through unstable intermediate binding states might be extremely rapid, the values of some rate constants might be quite large. This is problematic since we do not want to set a limit that may reduce the expressiveness of our model, but very large or small rate constants may lead to numerical instability when integrating the rate equation or solving for the equilibrium state.

For the case of a single binding site, a natural alternative is to use the dissociation constant *K*_*d*_ = *k*^*−*^*/k*^+^ to describe affinity and the time constant *τ*_obs_ = *k*^*−*^ + *K*_*d*_*k*^+^ = 2*k*^*−*^ to describe kinetics. Both parameters have simple, intuitive interpretations: *K*_*d*_ is the ligand concentration at which the binding site is half-saturated, and the *τ*_obs_ is the time constant with which the difference in saturation from 0.5 decays to 0 when free ligand concentration is fixed to the *K*_*d*_. *K*_*d*_’s can also be calculated for molecules such as GCaMP that bind multiple ligands, in which case the j th 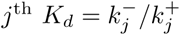 is the at ligand concentration where both the (*j −* 1)^th^ and *j*^th^ binding sites are almost totally filled or totally empty, so the *K*_*d*_’s do not tell us the concentrations at which the binding actually happens.

We therefore introduce a new quantity 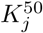, defined as the free ligand concentration at which precisely half the molecules have *j* or more ligands bound. Thus by eq. 18 we have the relations:

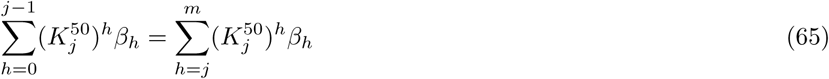

Therefore, given the *β*’s we can determine each *K*^50^ by solving a polynomial equation, and given the *K*^50^’s we can determine the *β*’s by solving a system of linear equations. Unlike for the *K*_*d*_’s, if we know that the molecule transitions from the apo to the saturated state occur over some ligand concentration range, then all the *K*^50^’s must lie in this range.

To describe the kinetics, we define 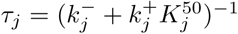, which is the time constant with which the (*j −* 1)^th^ and *j*^th^ binding states would reach equilibrium if the ligand concentration where set to 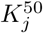 and all other binding state transitions were ignored.

Based on exploratory fits with multiple initializations, we set following loose constraints on the parameters: 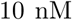 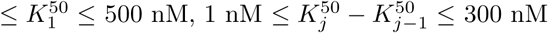 500 nM, 1 nM 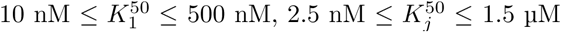 300 nM, 0.1 ms *≤ τ*_*j*_ *≤* 300 ms. Note that at any particular moment the actual kinetics of a binding step can be faster or slower than the corresponding *τ,* as [Ca^2^+] may be greater or lesser than the corresponding K ^50^

#### Initialization

Purities were initialized to 1, BAPTA contamination was initialized to 0 and GCaMP6s contamination was initialized to 1 (i.e. 25% saturation). Dead time was initialized to 1.5 ms. Values for the *K*^50^’s and *τ*’s were initialized randomly 50 times within their limits using latin hypercube sampling [83] as implemented in the **lhs** function of the **pyDOE** package. For each initialization, the *K*^50^’s and *τ*’s were fixed for the first 10 l-bfgs-b iterations.

#### Optimization

We minimized the objective function with the **minimize** function in the **optimize** module of SciPy, using the l-bfgs-b algorithm with options maxls = 100, mintol = 0.0 and maxcor = 20. To ensure that poor approximations of the Hessian matrix did not lead to false convergence, we repeatedly restarted the minimization function until the library reported convergence in a single iteration. We did not limit the number of iterations. Optimization with multiple initializations was parallelized using the **multiprocessing** python module.

Optimization proceeded with the following parameter bounds: 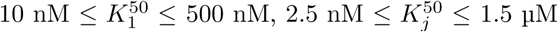 for *j >* 1, 1 ms ≤*τ*_*j*_ ≤ 5 s, *F*_*λ,i*_ > 0, 1 ms ≤*t*_dead_ ≤2 ms. Based on previous studies, purity values were limited to the range0.9 to 1.1 for GCaMP6s [47] and 0.75 to 1 for BAPTA [98]. Calcium contamination was limited to the range 0 to 0.01 for BAPTA and 0 to 10 for GCaMP6s.

### Fitting SBM parameters to *in vivo* fluorescence signals using known AP times

We developed procedures to perform global fitting of all SBM parameters to *in vivo* data and implemented them in the MATLAB programming language (Mathworks). We integrated the rate equation (12-14) using our custom ODE solver (see above), implemented through C MEX functions with openMP for parallelization over multiple fluorescence time series. We fit separate values of [GCaMP6s]^total^ and *F*_BG_ for each neuron. We fit our model only to fluorescence measurements with at least 1 AP in the preceding 20 seconds; other data were not included in the objective function (but were still used when calculating baseline fluorescence). Based on preliminary fits, we set *n*_buffers_ = 2. We explore other numbers of buffers in Figure 3 - figure supplements 3-4.

#### Normalization

Before further analysis, each sequence of fluorescence measurements was normalized by its median value. This allowed us to combine recordings from multiple microscope configurations, zoom factors and imaging rates for the same neuron even when absolute fluorescence values differed across these recordings. In the following material on model fitting procedures, *F*_*i*_ refers to these normalized measurements.

#### Simulation time steps

For neurons with data at multiple imaging rates, different values of *δ* were calculated for each fluorescence time series. The time of each fluorescence observation *t*_*j*_ was defined as the time when the laser focus passed through the center of the neuron. All electrically detected AP times were rounded to the nearest simulation time step. For each simulation time step, we first incremented [Ca^2+^] by Δ[Ca^2+^]_AP_ for each AP present, then integrated the rate equation (12-14) forward in time by *δ*. The first simulation time step begins Δ before the first fluorescence measurement (but much earlier for AP inference; see below). After every Δ*/δ* simulation time steps, we encounter a fluorescence observation for which we generate a predicted value based on the current binding state concentrations.

For example, if Δ = 99 ms, then *δ* = 9.9 ms, the first simulation time step begins 99 ms before the first fluorescence observation and the first prediction of a fluorescence value occurs after 10 ODE integrations of 9.9 ms each.

An exception to the above occurred when electrically recorded APs occurred before the first fluorescence measurement. In this case we included additional simulation time steps with the same value of *δ*, so that all APs up to 20 seconds before the first fluorescence measurement were included.

#### Objective function

We fit SBM parameters to *in vivo* data by minimizing the mismatch between predicted and observed values for normalized fluorescence. In order to weight all neurons in our datasets equally, we averaged square residuals over time for each neuron and then summed the result over neurons:

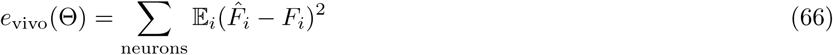

where the expectation with respect to *i* is an average over images frames for each neuron. We chose this objective function to ensure that all neurons contributed equally to the objective function, regardless of recording duration or imaging frame rate.

#### Model prediction of fluorescence using known baseline fluorescence values

The formula for model-predicted fluorescence (eq. 35) contains the unknown, time-varying scaling factor *B*_*i*_. To avoid having to explicitly determine the *B*_*i*_ for each time point, we use the concept of baseline fluorescence *F*_BL_, defined as the fluorescence that would be observed for a neuron at rest, with [Ca^2+^] = [Ca^2+^]_rest_ and all binding state concentrations at their equilibrium values. Thus we have by eq. 35

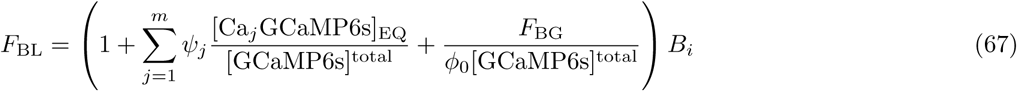

With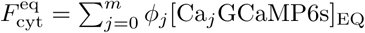 we can derive the relation

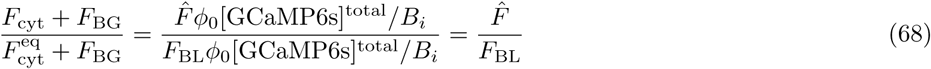

This quantity is simply the ratio of predicted fluorescence to baseline fluorescence, leading to eq. 7. Thus, if we known *F*_BL_ we can calculate 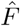 without explicitly calculating *B*.

#### Estimation of baseline fluorescence

For each contiguous sequence of *T* fluorescence measurements, we estimated baseline drift using fluorescence values and electrically detected APs. We first identified all fluorescence measurements > 1.5Δ before subsequent APs and > 6 seconds after previous APs, for which a binary mask *M* ^0^ was set to 1. To initialize baseline fluorescence, we interpolated and smoothed fluorescence measurements, using a kernel *K*_*b*_ consisting of a sum of two Gaussian functions with standard deviations 5 seconds and 500 ms. We calculated

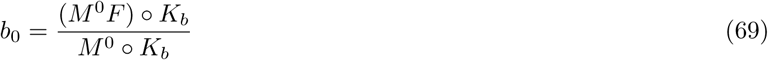

where the multiplication and division are carried out elementwise.

This initial estimate showed less perturbation due to AP discharges than a rolling average or median, but still increased during long AP bursts. To refine it further, we minimized an objective function *e*_BL_ that penalized both mismatch between data and predictions as well as excessive baseline fluctuations, while assuming exponential decay of fluorescence to baseline starting 5.5 seconds after AP activity.

We first computed a second binary mask *M* ^1^, defined in the same way as *M* ^0^ but starting 5.5 seconds after each AP instead of 6. Let the j-th contiguous sequence of time points for which *M* ^1^ = 1 be denoted *G*_*j*_, and let 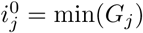 min(*G*_*j*_) be the first time point in *G*_*j*_. For each such sequence we introduce a free variable *A*_*j*_ representing the initial ratio of fluorescence to baseline for the first time point in *G*_*j*_. We then sought to minimize the objective function

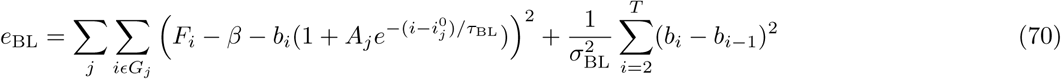

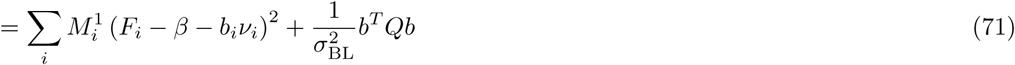

Where *Q*_00_ = *Q*_*T* *T*_ = 1, *Q*_*ii*_ = 2 for 1 *< i < T, Q*_*i,i*+1_ = *Q*_*i*+1*,i*_ = − 1 and *Q*_*ik*_ = 0 otherwise. We have also defined 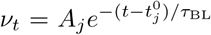 if *t* is in some *G*_*j*_ and *ν*_*t*_ = 1 otherwise. We used 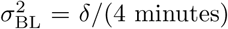, where *δ* is the time between fluorescence measurements. That is, we expect the baseline to drift with a standard deviation equal to its starting value every four minutes.

The free parameters of this objective function are the baseline values *b*_*i*_ > 0, the offset *β*, the initial amplitudes for each segment *A*_*j*_ > 0 and the time constant *τ*_BL_ > 0. The objective function consists of squared residual terms over time points for which *M* ^1^ = 1, as well as a regularization terms over all time points.

We alternated between minimizing *e*_BL_ over ({*A*_*j*_}, *τ*_BL_), *β* and *b*. We first optimized jointly over *τ*_BL_ and all the *A*_*j*_. We used golden section search with parabolic interpolation over *τ*_BL_ (Matlab’s **fminbnd** function), while calculating the optimal value for each *A*_*j*_ in closed form given *τ*_BL_ using a rectified linear regression. *τ*_BL_ was initialized at 500 ms.

We then updated *β* in closed form to

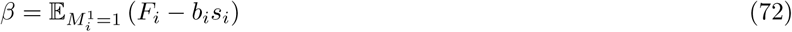

We then minimized *e*_BL_ over *b >* 0. If we ignore the positivity constraint, setting the derivative to zero yields the system of linear equations.

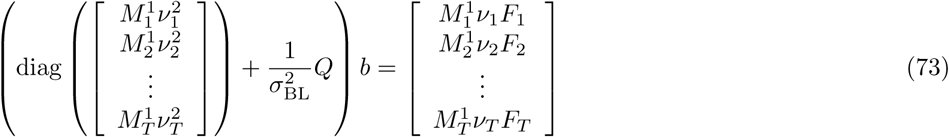

We accept this solution for *b* if it is nonnegative for all *t*. Otherwise, we must solve a quadratic programming problem (we implemented this contingency using MATLAB’s **quadprog** function, but this code was never called for baseline fitting of any data).

We iterated these alternating updates until the improvement in *e*_BL_ after all three updates together was less than machine precision (MATLAB’s **eps** function, about 2.2e-16), or when the improvement was less than 0.001% of the previous objective function value. Computation time for baseline fitting was negligible compared to the overall model-fitting procedure.

Finally, we calculate *F*_BL_ = *b* + *β*. The results of this model fitting procedure are shown in Figure 3 - figure supplement 1.

**Table 3.** Multiple initializations for 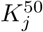

| Initialization | equilibrium binding state sequence | $K_1^{50}$ (nM) | $K_2^{50}$ (nM) | $K_3^{50}$ (nM) | $K_4^{50}$ (nM) |
| --- | --- | --- | --- | --- | --- |
| 1 | $0 \rightarrow 4$ | 350 | 352.5 | 355 | 357.5 |
| 2 | $0 \rightarrow 3 \rightarrow 4$ | 100 | 102.5 | 105 | 600 |
| 3 | $0 \rightarrow 2 \rightarrow 4$ | 100 | 102.5 | 597.5 | 600 |
| 4 | $0 \rightarrow 2 \rightarrow 3 \rightarrow 4$ | 100 | 102.5 | 351.3 | 600 |
| 5 | $0 \rightarrow 1 \rightarrow 4$ | 100 | 595 | 597.5 | 600 |
| 6 | $0 \rightarrow 1 \rightarrow 3 \rightarrow 4$ | 100 | 348.8 | 351.3 | 600 |
| 7 | $0 \rightarrow 1 \rightarrow 2 \rightarrow 4$ | 100 | 348.8 | 597.5 | 600 |
| 8 | $0 \rightarrow 1 \rightarrow 2 \rightarrow 3 \rightarrow 4$ | 100 | 266.7 | 433.3 | 600 |

#### Closed-form solution for *F*_BG_

When all other parameters of the model are fixed, we can minimize *e*_vivo_ (eq. 66) with respect to *F*_BG_ in closed form. For each neuron, we have by eq. 68

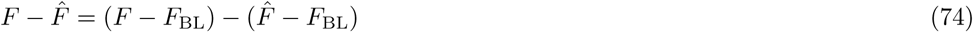

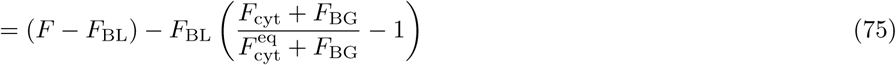

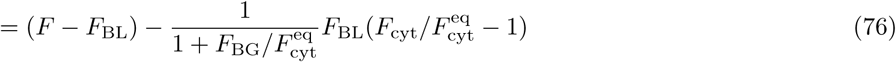

so minimizing *e*_vivo_ amounts to a simple linear regression with unknown slope 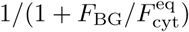. For model fitting, we imposed the restriction 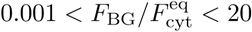.

### Initial parameter values

We initialized [GCaMP6s]^total^ = 20 µM and *ψ* = [0.45, 0.45, 22.5, 45]^*T*^. The [Ca^2+^]_rest_ was initialized to 50 nM [60, 116, 80, 142]. As previous studies suggest the extrusion time constant for calcium *τ*_ex_ is around 5 ms in dendrites [60, 115], we initialized this parameter to 20 ms to describe slower extrusion at the soma. The total calcium influx per AP Δ[Ca^2+^]_AP_ was initialized to 5 µM to be consistent with past measurements of total calcium influx [107] as well as smaller measured increases in free calcium [60, 142]. We initialized the endogenous buffers *K*_*d*_’s to 500 nM and their time constants to 15 ms and 1 s. We initialized all total buffer concentrations at the same value, chosen so that the calcium binding ratio of a neuron without GCaMP6s at rest was 125 [60, 99, 80]. The calcium binding ratio of buffer *ℓ* was defined as the derivative of the concentration of calcium bound to the buffer with respect to free calcium concentration:

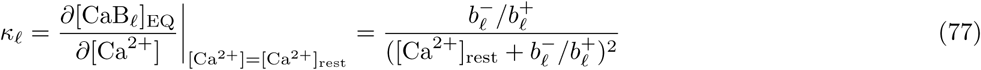

To describe the affinity and kinetics of calcium binding to GCaMP6s, we used the same reparameterization as for *in vitro* data (see above), consisting of time constants *τ*_*j*_ and affinity parameters 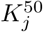. We initialized each *τ*_*j*_ to 100 ms. We used 8 different initializations for the 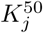 (Table 3), corresponding to the 8 possible ascending sequences from zero to four ions bound. These 8 initializations corresponded to the presence or absence of the 3 intermediate binding states at equilibrium as free calcium concentration was increased (absent binding states could still occur transiently in binding state kinetics evoked by AP discharge). When a a binding state [Ca_*j*_GCaMP6s] was absent from a the equilibrium binding state sequence of an initialization (Table 3, second column), we also imposed the restriction 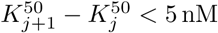 5 nM during optimization. After the optimization converged, we then released these restrictions and optimized further.

#### Optimization

We computed the gradients of the error function using finite differences. For each parameter, we set the step size for the finite gradient as 5e-7 or 5e-7 times the absolute value of the parameter value, whichever was larger. We used a positive step only (asymmetric finite differences). We optimized over *F*_BG_ for each neuron in closed form during finite difference calculation and line search.

Using this procedure for gradient calculation, we minimized *e*_vivo_ using the sequential quadratic programming technique available in MATLAB R2014a’s **fmincon** function. We used the following settings: algorithm = ‘sqp’, tolfun = 1e-7, tolx = 1e-8, tolcon = 1e-10, gradobj = ‘on’. We did not limit the number of iterations.

Optimization proceeded with the same parameter bounds on 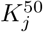 and *τ*_*j*_ as for *in vitro* data, and the additional bounds 2.5 ms *≤ τ*_ex_ *≤* 300 ms, 10 nM *≤* Δ[Ca^2+^]_AP_ *≤* 100 µM, 10 *≤ ϕ*_4_ *≤* 80, *ϕ*_*j*_ *≥ ϕ*_*j−*1_, 250 µs 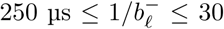 30 second, 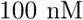 100 µM and 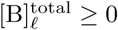. We also enforced the bound 0.5 µM *≤* [GCaMP6s]^total^ *≤* 300 µM for each neuron.

#### Hyperparameter fitting

After model-fitting, we obtained maximum likelihood estimators of the firing rate hyperparameters *k*_*λ*_*, θ*_*λ*_ by applying MATLAB’s **gamfit** function to the true firing rates. We also fit *µ*_*G*_, σ_*G*_ by maximum likelihood, yielding:

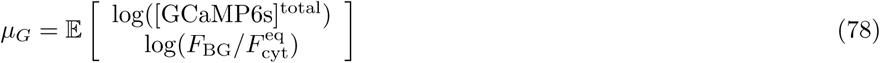

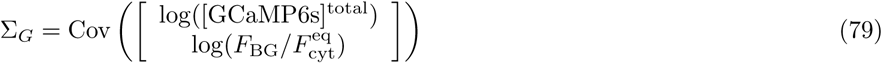

where the expectation and covariance are over neurons. We refer to *µ*_*G*_, σ_*G*_ as hyperparameters since they define a distribution on the parameters *F*_BG_ and [GCaMP6s]^total^, but for fixed values of *F*_BG_ and [GCaMP6s]^total^ in a specific neuron they do not affect the likelihood of that neuron’s fluorescence data.

#### Leave-one-out cross validation

In order to validate our procedures for AP inference and for estimation of *F*_B_ and [GCaMP6s]^total^ from fluorescence measurements alone (see below), we first performed model fitting on all neurons but one in our dataset. We then discarded the neuron-specific parameters [GCaMP6s]^total^ and *F*_BG_ and transferred the remaining parameters and hyperparameters to the last neuron. This allowed us to infer the remaining neuron’s AP sequence along with *χ*, [GCaMP6s]^total^ and *F*_BG_ from fluorescence data alone (see below), without using any parameters trained on its data.

When fitting the SBM using only a single neuron as training data, the algorithm was trained on neuron 1 and tested on neuron 2, trained on neuron 2 and tested on neuron 3, etc. If the neuron previous to the testing neuron was an interneuron (and hence could not be fit using our optimization procedures due to the lack of silent periods for fluorescence baseline estimation), the previous pyramidal neuron was used instead.

### Sequential Monte Carlo-based AP inference

We aim to infer the presence or absence of AP discharge at each SBM simulation time step *j*, denoted *s*_*j*_*ϵ{*0, 1*}*. We first describe procedures for inferring the posterior probability that each *s*_*j*_ = 1 given fluorescence data when all parameters are known, followed by procedures for inferring neuron-specific data from fluorescence data alone.

We used Sequential Monte Carlo to approximate posterior distributions over a vector *y* of hidden states using finite weighted sums of delta functions. Each combination of state vector and weight is termed a particle. For the SBM *y* consists of *s*, *ρ* and all molecular species concentrations. We update the particles once for every fluorescence measurement, yielding one new value for *ρ* and Δ*/δ* new values for *s* and the binding state concentrations. Thus at the *i*^th^ fluorescence measurement the state vector includes spiking and binding state concentrations for the simulation time step range *U*_*i*_ = (*i −* 1)Δ*/δ* + 1: (*i* − 1)Δ*/δ*. For *m* = 4 binding steps and *n*_buffers_ = 2, we have the state vector

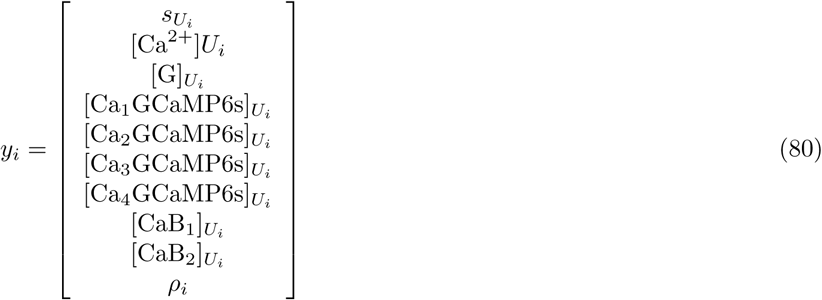

#### Filtering distribution

Given the SBM model parameters, the filtering distribution *P* (*y*_*i*_ *|F*_1:*i*_) describes the posterior probability distribution on the hidden states *y* at the time of the *i*^th^ fluorescent measurement *F*_*i*_, given all fluorescence data up to and including *F*_*i*_. We approximate the filtering with a finite number of samples, using the standard SMC estimator:

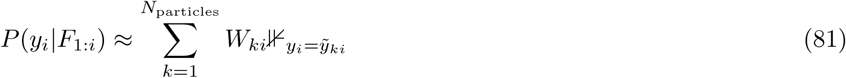

where 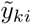 is the state vector value for the k^th^ particle at the ith fluorescence measurement, 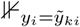 is an indicator function and Σ _*k*_W _*ki*_ = 1 for all i. In eq. 81, *y* _*i*_ areunknown hidden variables whose probability distribution we approximating, and the particle states 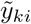 are known values we are using for this approximation. The particles are propagated from *t*_*i−*1_ to *t*_*i*_ by sampling a new *y* for each *k* from the proposal *q*(*y*_*i*_*|y*_*i−*1_). The weights are then updated according to the standard “sequential importance sampling” SMC update:

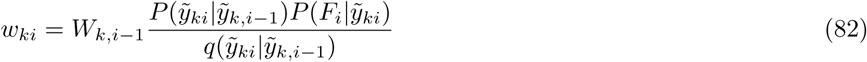

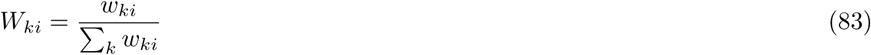

The data likelihood 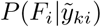 is defined by eq. 38, while the transition probability 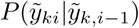 is defined by eq. 31 and

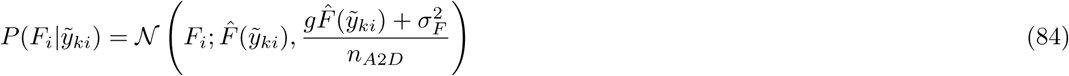

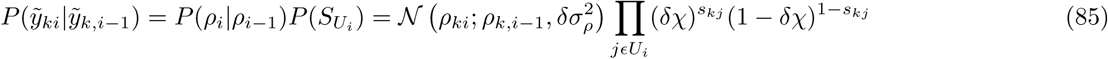

Our proposal *q* first samples spiking from a Bernoulli distribution *q*_*s*_(*j*) that is different for each *j* but does not depend on the state vector (see below). Once *s*_*j*_ has been sampled for each *jcU*_*i*_, binding state concentrations are calculated deterministically by integrating the rate equation with our custom ODE solver. We sample *ρ* from the transition prior, a normal distribution with variance 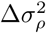, so that the terms involving *ρ* cancel from the numerator and denominator of eq. 82. Together these updates produce an SMC estimate of *P* (*y*_*i*_*|F*_1:*i*_) given *F*_*i*_ and the SMC estimate of *P* (*y*_*i−*1_*|F*_1:*i−*1_).

#### Resampling

After the SMC update has been carried out many times, if the proposal distribution does not precisely match the posterior most of the particle weights will approach zero and a small number of particles will dominate the filtering distribution. This situation, known as particle degeneracy [35] can lead to increasing variance in the estimates of the filtering distribution for increasing *t*. To prevent this, a resampling operation is carried out in which particles are sampled with replacement from a discrete distribution *q*_RS_(*k*) on *{*1*, · · ·, N*_particles_*}* using the systematic resampling technique [73]. For each particle index 1 *≤ k ≤ N*_particles_, a new index *k′* is sampled from *q*_RS_. The weights and states are then updated according to:

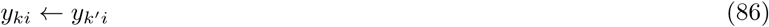

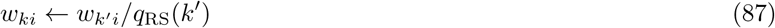

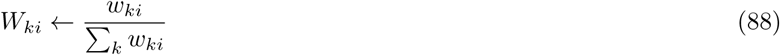

Most SMC algorithms choose *q*_RS_ to be equal to the current filtering distribution so that *W*_*ki*_ are all 1*/N*_particles_ after resampling, but other distributions have been used as well [41]. We used specialized distributions tailored to our model to reduce the variance of SMC algorithm results, as described below.

After each update of the particle weights, we calculated the effective sample size

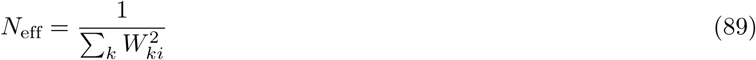

and resampled if *N*_eff_ *< N*_particles_/2. When illustrating the particle filter technique in Figure 5a-c, we resampled at every fluorescence measurement.

#### Initial state distribution

To assign weights and states at the first time step we sample random AP sequences before the first fluorescence measurement and calculate their probabilities based on *χ*. We also sample initial values of *ρ* from a a broad prior *π*_*ρ*_(*ρ*_1_). We first defined *F*_init_ as the 10^th^ percentile of the first 120 seconds of fluorescence values. If this gave a negative value, then *σ*_*F*_ was assigned to *F*_init_ instead. We next sampled *ρ*_1_ from the prior

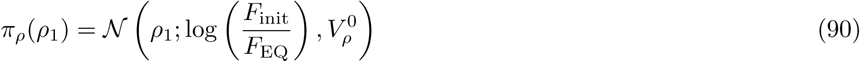

With 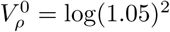.

For each particle *k* we set [Ca^2+^] = [Ca^2+^]_rest_ and set the concentrations of all other molecular species to their equilibrium values using eq. 18. We next a random AP sequence *s*_*k,U*_1, with *U*_1_ in this case indexing 2.5 seconds before the first fluorescence measurement, and update the concentrations of indicator and buffer binding states accordingly by integrating the rate equation. As for the standard SMC state update, the probabilities of randomly sampling an AP for each simulation time step are taken from *q*_*s*_, the form of which is specified below. We then calculated for each particle *k* the maximum likelihood estimate of *ρ* given the indicator binding state concentrations and a fluorescence observation of *F*_init_:

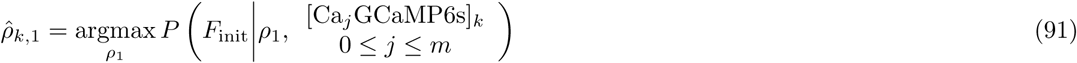

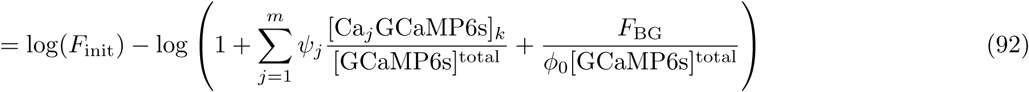

and calculated the Laplace approximation [19] to the likelihood of F init given ρ, centered at 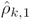

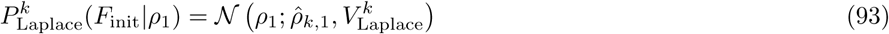

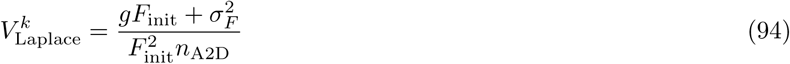

Using this approximation and the prior *π*_*ρ*_, we calculated the sampling distribution for each *k* by multiplying the prior and the Laplace approximation:

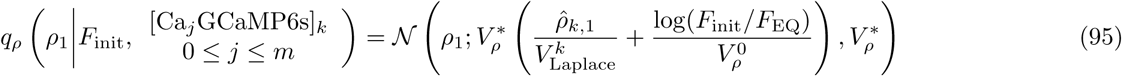

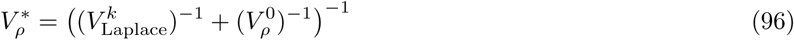

We then sampled each *ρk*,1 and calculated weights as the ratio of prior and sampling probabilities:

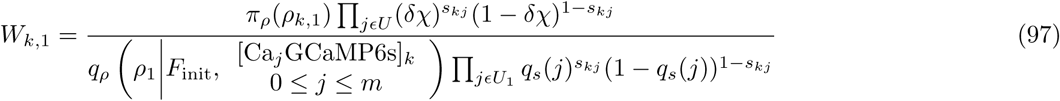

#### Smoothing distribution

To estimate the smoothing distribution *P* (*y*_*i*_ *F*_1:*T*_), we used a standard filter-smoother [73] with lag of 500 ms. We kept track of each particle’s history of previous state values as we advanced the filtering distribution forward in time. Then, at each time point the current filtering weights together with the state values of each particle’s ancestor 500 ms back in time define the smoothing distribution. For particle *k* at fluorescence measurement *i*, the smoothing weight 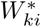 is defined as the summed weights of its descendants 500 ms after *t*_*i*_ (or for *t*_*i*_ *> t*_*T*_ − 0.5, descendant weights at *t*_*T*_). 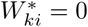 = 0 for particles with no descendants 500 ms in the future. Smoothing weights are then used to calculate posterior means and variances of *y*:

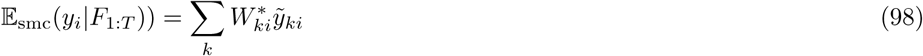

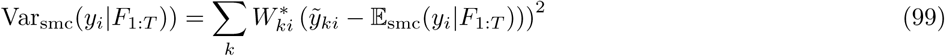

where the squaring in the definition of Var_smc_ is carried out elementwise.

#### Sampling and resampling techniques for SMC-based AP inference with the SBM

The quality of an SMC algorithm, measured by the number of particles required to obtain accurate filtering and smoothing distributions, depends strongly on the choice of proposal distribution *q* [34, 19, 35]. When the generated samples match the posterior poorly, many will be rejected during resampling events and most computation will be wasted. For the SBM, the key question is whether the algorithm extends AP sequences forward in time in a way consistent with fluorescence data. Successful inference requires at least some particles to generate an AP at the time of each true AP, and for these particles to survive all resampling steps until the filter-smoother lag has been reached. However, if the needed APs are not sampled or are lost to resampling, the algorithm will fail to detect the true AP. In this section, we describe three techniques for designing proposal and resampling distributions to generate the needed spikes and ensure their survival through resampling steps.

##### Increased AP discharge probability in the sampling distribution

*For δ* = 10 ms and our median firing rate of *χ* = 0.16 Hz, the prior probability *P* (*s*_*j*_ = 1) is only *p*_AP_ = *χδ* = 0.0016. Therefore, even with many particles the chance of simply failing to generate spike trains consistent with the data is not insignificant when sampling *s*_*j*_ from its prior distribution. We therefore sample APs with a higher probability than the prior to generate a greater diversity of AP sequences. While this mismatch between the transition prior and proposal will cause an overall increase in the frequency of resampling when APs do not occur, the fit to the data will be better when APs are present.

We sampled APs with probability max(*p*_AP_, 0.01). We demonstrate how sampling APs at a higher rate than the prior can reduce variability in the SMC algorithm’s output in Figure 5 - figure supplement 2.

##### Multiround filtering with time-varying proposals

To further improve the quality of sampled AP sequences, we first run the SMC algorithm with increased probability of AP sampling as above to obtain the smoothing distribution and 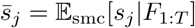 We then designed a time-varying proposal on AP discharge

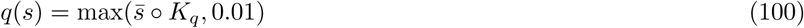

where ○ denotes convolution and *K*_*q*_ is a Gaussian kernel with *σ* = 100 ms. We extend the calculation of both 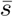 and the time-varying proposal *q*(*s*) to the full 2.5 seconds before the first fluorescence measurement for which SBM simulations are carried out (see section “Initial state distribution” above). We show how this technique further reduces Monte Carlo variance in Figure 5 - figure supplement 2.

##### Resampling to the proposal

For GECIs in general and for GCaMP6s in particular, an additional difficulty for SMC algorithms arises from the delay between AP discharge and peak fluorescence. The first fluorescence measurement after AP discharge may not show a strong increase in fluorescence, and for higher imaging rates this can increase to 2 or 3 measurements. Consequently, for particles generating an AP at the correct time, the resulting increase in their weights relative to other particles without the correct AP may not appear until the algorithm has passed over at least one second of fluorescence measurements. So even even if APs are sampled by the SMC algorithm at the correct time, they may be lost during resampling steps before the predicted fluorescence increase has been fully observed. Since the *p*_spike_ is low, sampling an AP will decrease the probability weight of a particle at the next fluorescence measurement compared to other particles (eq. 82). This will both reduce *N*_eff_ to make resampling more likely and increase the chance that the particle with the AP will be lost.

This problem is not alleviated by using an increased or time-varying *q*(*s*), since while this causes more APs to be sampled when needed it also decreases the weights of particles with APs even further due to the presence of *q* in the denominator of eq. 82. We might sometimes be fortunate enough not to encounter a resampling event between the sampling of an AP and the subsequent correct prediction of a fluorescence increase, but this will not always be the case. The possibility of these two different outcomes, where the spikes needed to match the data are retained or are lost to resampling, sharply increases the variability of the SMC algorithm over multiple runs.

We therefore introduce a novel modification of the SMC algorithm which we term “resampling to the proposal,” designed specifically to deal with state transitions that have strong but delayed effects on the predict measurements. In addition to the standard raw weights 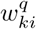 and normalized weights 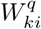, we also maintain a second set of raw weights *w*^*q*^and normalized weights *W*_*ki*_. The second set of weights, which we refer to as “proposal weights” are obtained by calculating the particle weight updates as if the time-varying proposal on APs were also the prior on APs. Both sets of weights share the same statevariables 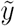 This leads to the update rules:

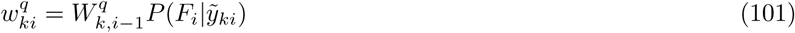

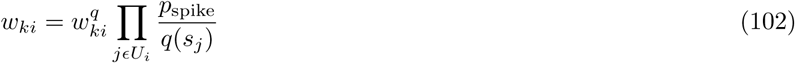

with normalized weights calculated as above. Note that *s* does not appear in the update of 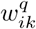 since the proposal and*ik*prior on *s* are the same, so the numerator and denominator in eq. 82 cancel. Thus particles with more APs do not have lower resampling weights 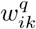, although they do have lower standard weights *w*_*ik*_.

To reduce the chance of losing needed APs to resampling events, we simply set the resampling distribution to be equal to the normalized proposal weights: 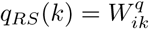. Resampling is used to select a single set of indices, but the standard and resampling weights are updated differently. The updates now take the form

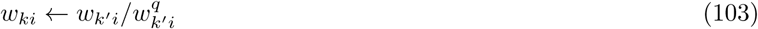

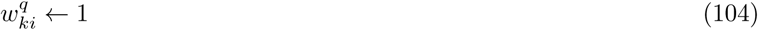

with normalized weights calculated as above. We calculate the effective sample size using the proposal weights 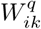 Finally, when calculating the smoothing distribution we use the original weights *W*_*ki*_ to calculate 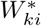 as before. Unlike the proposal weights, the standard weights used for smoothing take into account *χ* through a weight decrease for every discharged AP, but they also reflect fluorescence measurements acquired after those APs. This technique allowed us to sample APs with high probability from our time-varying prior, keep those spikes through resampling events and still obtain the smoothing distribution for the original problem. We show how this further reduces Monte Carlo variance of the SMC algorithm in Figure 5 - figure supplement 2. In order to illustrate the SMC technique in Figure 5c, the area of the black dots was chosen to be proportional to proposal weights, using a *q* based on increased AP discharge probability but not multiround filtering.

To our knowledge, resampling to the proposal is the first approach to use two particle filters with the same states but different weights to deal with lags between state changes and changes in the observed data. Previous approaches to dealing with such lags have generally focused on modifying SMC states in the past using MCMC steps [46, 33], but these approaches are often unreliable for time series more than a few hundred measurements [3] and could impose impractical computational and memory requirements for GPU-based SMC with many particles. However, the idea of coupling two particle filters to use the same indices during resampling is not unprecedented, and has been applied to compute finite difference gradients of the data likelihood with respect to static model parameters [22]. However, the use of proposal weights for resampling distinguishes resampling to the proposal from previous approaches, and when combined with proposal distributions based on multi-round filtering may prove useful for a wide range of nonlinear filtering problems.

#### Fitting a spike train to posterior moments

The SMC algorithm described above calculates each 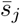. These values are discretized in time with a step size of *δ*, and represent the marginals of the joint posterior distribution provided by the SMC smoother. While for certain applications, such as calculation of receptive fields, 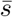 itself can be used directly, at other times it is preferable to infer a single AP sequence from a fluorescence time series.

We therefore developed a procedure to infer a single, non-time-discretized AP sequence from 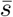 Inspection of the time-varying posterior spiking probability from the SMC smoother revealed that true, electrically detected APs gave rise to Gaussian-like peaks in 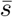, which can be interpreted as uncertainty in the times of APs. We therefore fit to 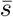 an approximation consisting of a sum of *n*_APs_ Gaussian functions with means *µ*_*k*_ and standard deviations *σ*_*k*_. We modeled the individual values of 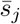 as integrals of the Gaussian functions over each time step of length *δ*:

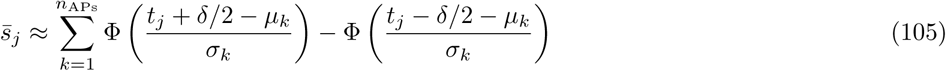

which we optimize in least squares with the restriction that δ/2 ≤ σ k ≤ 50 ms.

We determine *n*_APs_ and the values of *µ*_*k*_ and *σ*_*k*_ using a greedy algorithm. First, we attempt to add a new spike while varying the new *σ* value in 100 steps, ranging from 1% to 99% of the interval between the minimum and maximum allowed values. For each value of *σ* for the new AP, we shift the mean to each *t*_*j*_, and choose the location which decreases the sum of square errors most. We then adjust all the *µ*’s and *σ*’s to minimize the sum of square errors using Gauss-Newton optimization, while allowing both to vary continuously without discretization. The algorithm terminates when no new spikes can be added to reduce the sum of square errors, yielding the *µ*’s as spike times and the *σ*’s as temporal uncertainties. For the step of adding new spikes, we precompute a lookup table of all necessary ϕ values for speed. This method is illustrated in Figure 5 - figure supplement 6.

### Parameter estimation from fluorescence data alone

Most of the SBM’s parameters are determined by fitting *in vitro* binding assay data or fitting *in vivo* fluorescence data with true AP times known. However, when detecting APs from fluorescence recordings without simultaneous cell-attached recordings or where the true APs have been held out for testing, the remaining parameters (*χ, σ*_*F*_, [GCaMP6s]^total^*, F*_BG_) must be determined from fluorescence alone. For this purpose, we use a heuristic method to determine *σ*_*F*_ and an expectation-maximization method for *χ*, while [GCaMP6s]^total^ and *F*_BG_ are determined using an SMC-based estimate of the marginal data likelihood.

**Estimation of** *σ*_*F*_

We estimated *σ*_*F*_ as

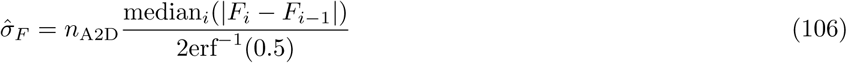

where erf^−1^ is the inverse error function. The numerator is the median absolute fluorescence difference across neighboring image frames, while the denominator is median absolute difference for two values drawn from a standard normal distribution.

#### EM estimation of *χ*

With *σ*_*F*_ known and for fixed values of [GCaMP6s]^total^ and *F*_BG_, we perform maximimum a posteriori (MAP) estimation of *χ* using the SMC version of the Expectation-Maximization (EM) algorithm [28]. In the E-step, use the SMC algorithm to estimate the smoothed posterior on *s* given *F*. In the M-step, we use this posterior to maximize the expected complete log-likelihood times the parameter prior:

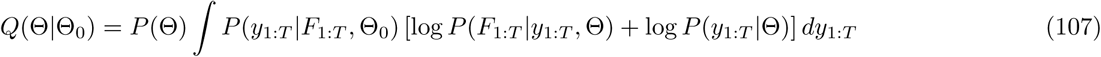

where Θ_0_ is the current estimate of the parameters and Θ is the new estimate. The SMC version of the EM algorithm uses the SMC smoothing distribution, which is a sum of delta functions, to approximate *P* (*y*_1:*T*_ *F*_1:*T*_, Θ_0_). Thus maximization of eq. 107 over *χ* reduces to maximization of a gamma prior times a product of Bernoulli likelihoods, which we consider in the log domain:

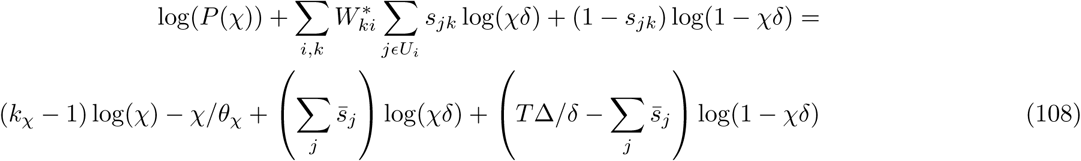

Having computed 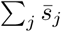 using the SMC smoother, we maximized the expression in eq. 108 numerically with the Matlab function **fminbnd**.

We found that a single EM-iteration was sufficient to estimate *χ*, in the sense that multiple iterations did not improve inference of the AP sequence or the marginal data likelihood (see below). Since we already use two SMC passes over the data to reduce Monte Carlo variance (see section “Multiround filtering with time-varying proposals” above), we simply estimated *χ* after the first round of SMC and used this value for the second round.

#### Marginal likelihood

In addition to providing filtering and smoothing distributions, SMC algorithms can also be used to estimate the marginal data likelihood *P* (*F*_1:*T*_ *|*Θ), where Θ is a vector of model parameters that determine the transition prior, the initial state distribution and/or the distribution on observed data values given the model state. We seek to compute this function in order to perform model fitting: that is, to find values of Θ consistent with our data. The marginal likelihood can be a difficult quantity to calculate since it involves integration over the joint distribution on hidden states over all time points:

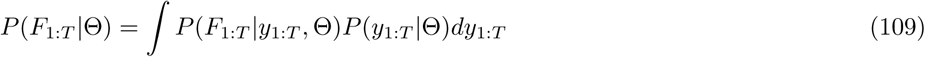

We follow the standard approach of decomposing

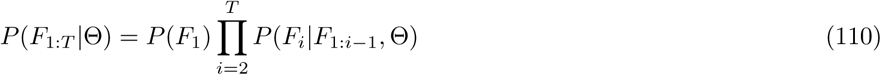

The terms of this product, which are the likelihoods of each fluorescence measurement condition on previous measurements, can be calculated using the SMC weights [104]:

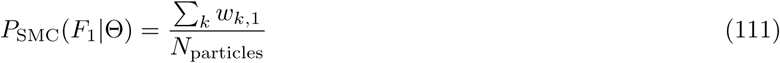

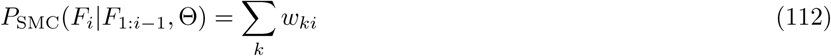

### Maximization of tempered posterior

For a given choice of [GCaMP6s]^total^ and *F*_BG_ we can obtain an estimate 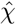 of *χ* via EM, and then calculate the SMC estimate of the marginal likelihood. Multiplying the result by the prior on *χ*, [GCaMP6s]^total^ and *F*_BG_ yields

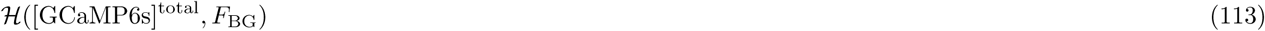

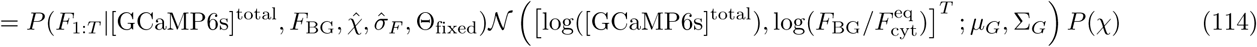

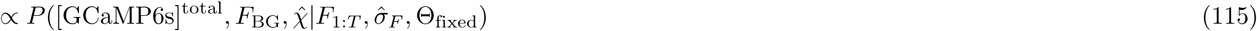

Where Θ_fixed_ contains SBM parameters that do not vary over neurons. Since 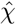 is a partial optimizer of the posterior given the other parameters, maximizing ℋ over [GCaMP6s]^total^ and *F*_BG_ yields a maximum a posteriori estimate over all three parameters.

Direct maximization of ℋ is not possible, since even for 10^5^ particles the variance of log ℋ for fixed input can reach 10 for as little as 30 seconds of imaging data (Figure 5 - figure supplement 2). Instead, we seek to characterize the shape of the function ℋ and find its maximum in a way that is robust to the variability in the calculated value of ℋ ([GCaMP6s]^total^*, F*_BG_) over multiple runs arising from random number generation within the SMC algorithm.

To do this, we use the adaptive independent Metropolis Hastings (AIMH) technique of [48]. This method draws samples from a density proportional to a function while simultaneously fitting a Gaussian mixture to the sample set by harmonic K-means clustering. The algorithm was implemented in Matlab as described in [48], and run for 400 iterations. For harmonic K-means clustering, we used the **kmeanhar** implementation of [140]. Our implementation accepts multiple time series for the same neuron, but by default we limit the length of the input data to 12500 fluorescence measurements for speed. Because we are interested only in the maximum of *ℋ* and not in characterizing the entire distribution, we increased the acceptance rate by “tempering” ℋ by a temperature 𝒯. That is, instead of using ℋas the target distribution for the AIMH technique, we used ℋ1*/𝒯*. To choose the temperature, we first calculated ℋ15 times at the initial values of [GCaMP6s]^total^ and *F*_BG_, which we set to equal *µ*_*G*_. We then choose 𝒯 so that log(ℋ1*/𝒯)* would have a variance of 0.1 or less:

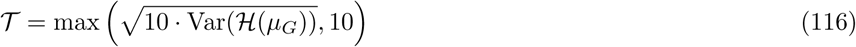

After 400 AIMH iterations, we returned the mode of the Gaussian mixture as a point estimate for ([GCaMP6s]^total^*, F*_BG_), along with the corresponding values of 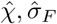. These procedures are illustrated in Figure 5 - figure supplement 5.

### Other inference methods

We implemented Matlab wrappers for all other AP inference methods, a Matlab GUI capable of running all algorithms on individual data segments, neurons and datasets and calculating accuracy measures based on the results. All individual and group data comparisons of various algorithms were carried out on the same data with the same pre-processing, including feature extraction and motion correction. For inference methods

### FOOPSI

Fluorescence data were first detrended using Matlab’s built-in **detrend** function. Next, the minimum value was subtracted from the detrended data, the result was divided by its maximum and a value of machine epsilon (2.22e-16) was added. Finally, the result was passed into the **fast ̲oopsi** function, with the parameter tau c = 0.5.

### CFOOPSI

Unmodified fluorescence values were passed to the **constrained ̲ foopsi** function, with no additional inputs. Commit 08468bc1b25c9b8617b861a6404bbf4576cf156c was retrieved from github.com/epnev/constrained-foopsi, and was used with CVX version 2.1, Build 1116 (d4cc5c5).

#### c2s

For c2s-s, we used the **c2s preprocess** command to upsample fluorescence data to 100 Hz as descriped in [Theis et al.], while providing metadata indicating which fluorescence time series were from the same neuron. We then used the **c2s predict** command to infer spike probabilities. For c2s-t, we used the **c2s leave-one-out** command to carry out training and to infer spike probabilities with cross validation. To calculate the number of parameters used by c2s, we trained it on our complete dataset (n = 26 neurons) with standard settings, for which c2s selected 10 PCA components consisting of 1000 elements but only 945 degrees of freedom after accounting for orthonormality constraints. The spike triggered mixture model used 20 feature parameters, 3 bias parameters, 6 weights and 30 predictor coefficients, for a total of 1004 parameters. Commit 9f12398a33a17e557b4a689f1dce902123c6f2eb was retrieved from github.com/lucastheis/c2s.

#### MLspike

We used MLspike with the published parameters for GCaMP6s [29], as the autocalibration feature failed whenever tested on our GCaMP6s data. These parameters were: a = 0.113, tau = 1.87, pnonlin = [0.81, −0.056], drift.parameter = 0.01. One call was made to the **tps ̲mlspikes** function for each different imaging rate in a neuron’s data, with the parameters and the frame interval passed along with fluorescence data. For an image frame *i* for which a single AP was inferred, we assigned it to the midway point between the times when the neuron was scanned: (*t*_*i−*1_ + *t*_*i*_)/2. For an image frame for which *k* APs were inferred, we distributed the inferred APs evenly across the interval between the two scan times, assigning the *j*^th^ AP time as *t*_*i−*1_ + Δ(*j −*0.5)*/k*. Commit 048122135c7d77457ee8c8c026a572ac40739c3f was retrieved from github.com/MLspike/spikes, along with commit 048122135c7d77457ee8c8c026a572ac40739c3f from github.com/MLspike/brick.

#### thr-*σ*

We first estimated fluorescence baseline *F*_BL_ as the 8^th^ percentile of fluorescence values over a 15 s window around each data point. For data points within 7.5 s of the start or end of a fluorescence recording, the first or last fluorescence value was repeated to substitute for the missing values. We then calculated a normalized Δ*F/F*_0_ measure as *y* = (*F* − *F*_BL_)/median(*F*_BL_). We then divided the fluorescence recording into 1 s windows and calculated the standard deviation of *y* in each window, and estimated the standard deviation of fluorescence noise *σ*_noise_ as the median over 1 s windows of the standard deviation of We then identified time points where *y* cross a threshold of 4*σ*_noise_. After each threshold crossing, additional threshold crossing events were ignored until after *y* decreased below 2*σ*_noise_. For each threshold crossing, an AP was assigned at the maximum value of *y* among the values after *y* increased over 4*σ*_noise_ and before it decreased below 2*σ*_noise_.

### Accuracy measures

To calculate the correlation of two AP sequences with AP times {*a*_*i*_} and {*b*_*j*_}, we convolved each AP sequence with a Gaussian kernel of width *σ* leading to a sum of Gaussian functions:

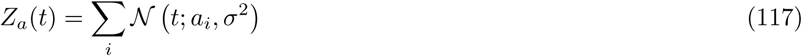

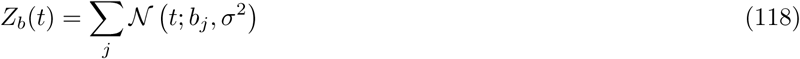

We then calculated the means of *Z*_*a*_, *Z*_*b*_, their product and their squares over the time window [*t*_0_*, t*_1_] over which the AP sequences are defined:

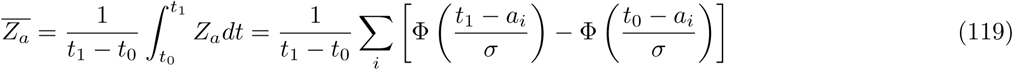

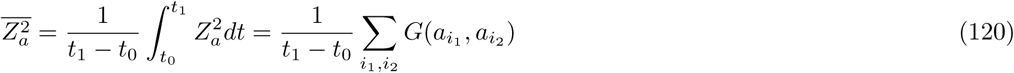

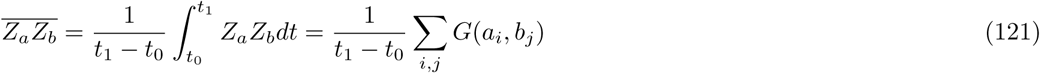

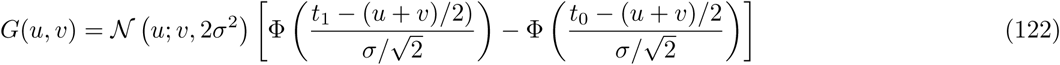

where ϕ is the Gaussian cdf and 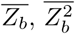, are defined correspondingly. Finally we calculated the correlation as

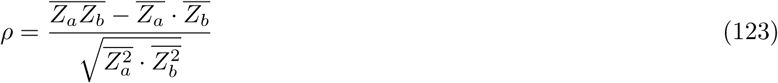

To calculate the average absolute timing error, we first collected all the outputs of each algorithm up to 500 ms before and after each isolated single AP. For algorithm that produced AP times (the SBM, MLspike and thr-*σ*), we then simply calculated the mean absolute time difference from the true AP. For other algorithms which produce expected AP counts (c2s) or unitless outputs (FOOPSI and CFOOPSI), we calculated (Σ _*i*_ *|t*_*i*_ *− t*_AP_*|y*_*i*_) / (Σ _*i*_ *y*_*i*_), where *y*_*i*_ is the algorithm’s output at time *t*_*i*_.

### Statistical analysis

To calculated the average fluorescence evoked by single APs in pyramidal neurons, we included only those without other APs in the preceding 5.5 s. We first calculated fluorescence baseline *F*_BL_ and Δ*F/F*_0_ values using fluorescence and true AP times as described above in the section “Estimation of baseline fluorescence” above. The baseline-fitting procedure also determined the time constant *τ*_BL_ with which fluorescence decays back to baseline after starting 5.5 s after AP discharge, as well as an exponential fit to all Δ*F/F*_0_ values at least 5.5 s after any APs. Therefore, to remove any effects of earlier AP discharge >5.5 s before single APs, we extrapolated the exponential fit forward through the fluorescence values around the single AP, and subtracted this from the Δ*F/F*_0_ values. We then subtracted the mean of Δ*F/F*_0_ values for image frames 0 to 200 ms before the AP. Finally, we calculated the Δ*F/F*_0_ values around single APs, using 75 ms bins with a bin edge centered on the time of AP discharge. For interneurons, the requirement that no other APs were present in the preceding 300 ms was used instead. To calculate fluorescence responses to bursts of multiple APs, we included bursts with up to 200 ms between the first and final AP, and no other APs afterwards, with a bin edge centered on the first AP time.

The amplitude or “height” of the response was defined as the maximum of the average from 0 to 300 ms for single APs, and 0 to 500 ms for bursts. Signal-to-noise ratio (SNR) was defined as the ratio of the single AP response amplitude to the standard deviation of the fluorescence noise. To calculate the noise standard deviation, we used all Δ*F/F*_0_ values 300 ms to 50 ms before isolated single APs as defined above.

All values are reported as mean +/- standard deviation unless otherwise stated.

### Data and code availability

All AP sequences, fluorescence signals and *in vitro* binding assay data are provided along with metadata (supplementary data). Raw fluorescence movies will be available in the future at caesar.de/sbm. Pre-alpha releases of code for analyzing *in vivo* and *in vitro* data are available at http://github.com/dgreenberg/sbmvivo and http://github.com/dgreenberg/sbmvitro.

## Supporting information

## Acknowledgements

We thank Stephan Irsen and Daniel Rudolph for assistance with cluster computing, Alexandr Klioutchnikov for assistance with *in vivo* optical imaging, Jeanine Klotz for technical assistance and Norbert Brenner for initial assistance with protein purification. We thank Jakob Macke for fruitful discussions and comments on the manuscript.

## Author contributions

*In vivo* GCaMP6s imaging experiments with simultaneous electrical recordings were designed by JNDK, DJW and DSG, and performed by DJW and UC. Imaging experiments with visual stimulation were performed by TH. *In vitro* binding assay experiments were designed by SW, YG, RS and DG and performed by SW. The SBM, model-fitting procedures and AP inference method were developed by DSG. *In vivo* data were analyzed by DSG and KMV. *in vitro* data were analyzed by DSG, AM, SW and YG. KMV and DSG developed the feature extraction technique. JTV helped design an earlier SMC method for small-molecule indicators. DSG and JNDK ran the project and wrote the paper.

---

**Figure 1–Figure supplement 1.**
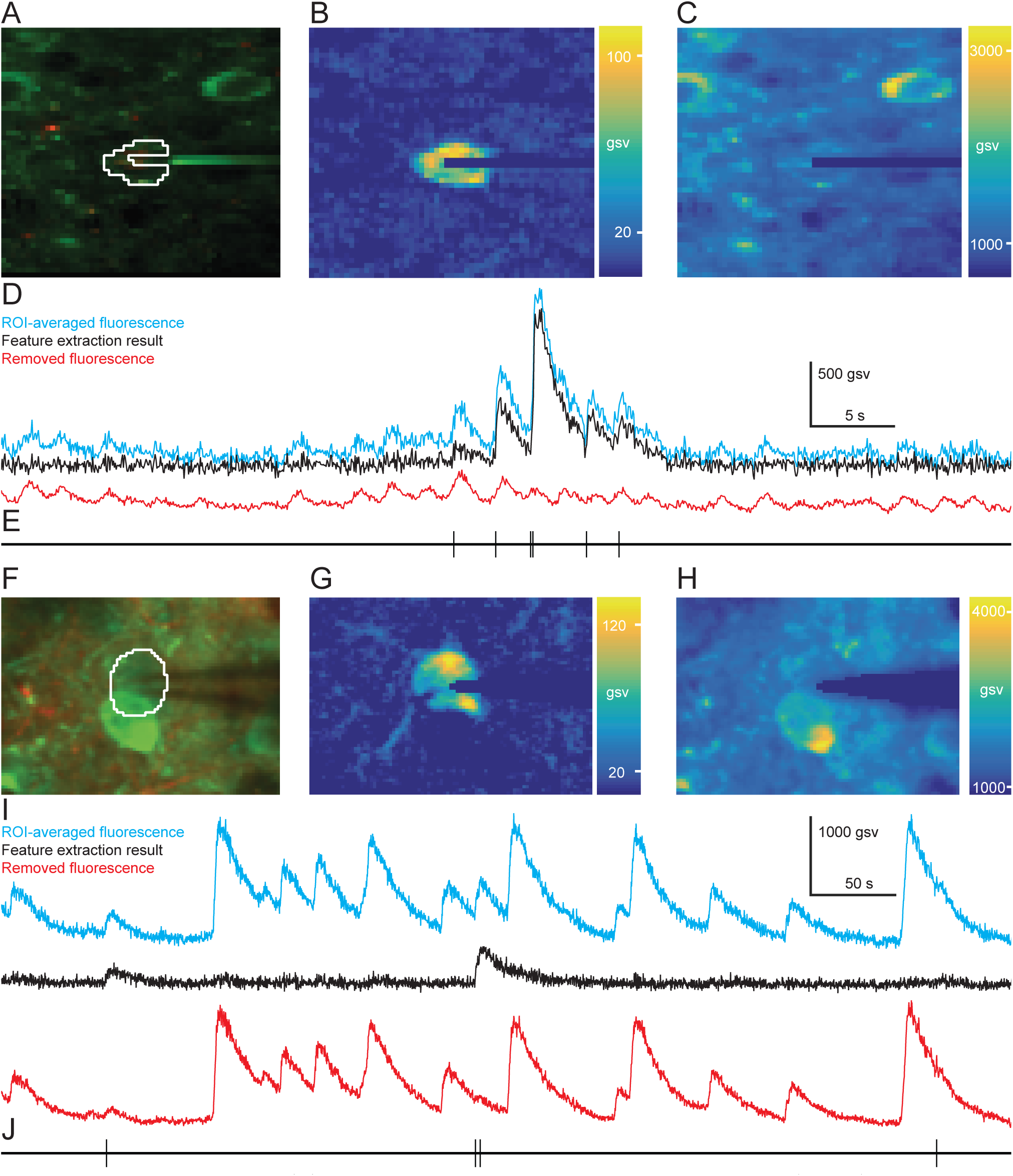
**(A)** Two-photon image acquired using galvanometric scanning (18.6 Hz), showing a neuronal population expressing GCaMP6s (green) in L2/3 of mouse visual cortex, with astrocytes stained using SR101 (red) and electrical recording of APs in a single pyramidal neuron. **(B)** Image showing the mean GCaMP6s fluorescence for each pixel after removal of extraneous signals using the feature extraction algorithm (see methods). **(C)** Image showing the mean GCaMP6s fluorescence removed from each pixel by the feature extraction algorithm. **(D)** Fluorescence time series calculated by a simple average (cyan) over pixels within a region of interest (ROI, white countour in (A)), along with the fluorescence removed by the feature extraction algorithm (red) and the resulting estimate of cytosolic fluorescence (black). Note the ongoing fluctuations in ROI fluorescence due to contamination from the neuropil background. **(E)** Electrically recorded AP times for the data shown in (D). **(F-J)**. As in (A-D), but for a second neuron recorded using resonance scanning (60 Hz). Note that in this case, most of the removed fluorescence arises from a second neuron, which due to the microscope’s point spread function overlaps the neuron recorded electrically.

**Figure 1–Figure supplement 2.**
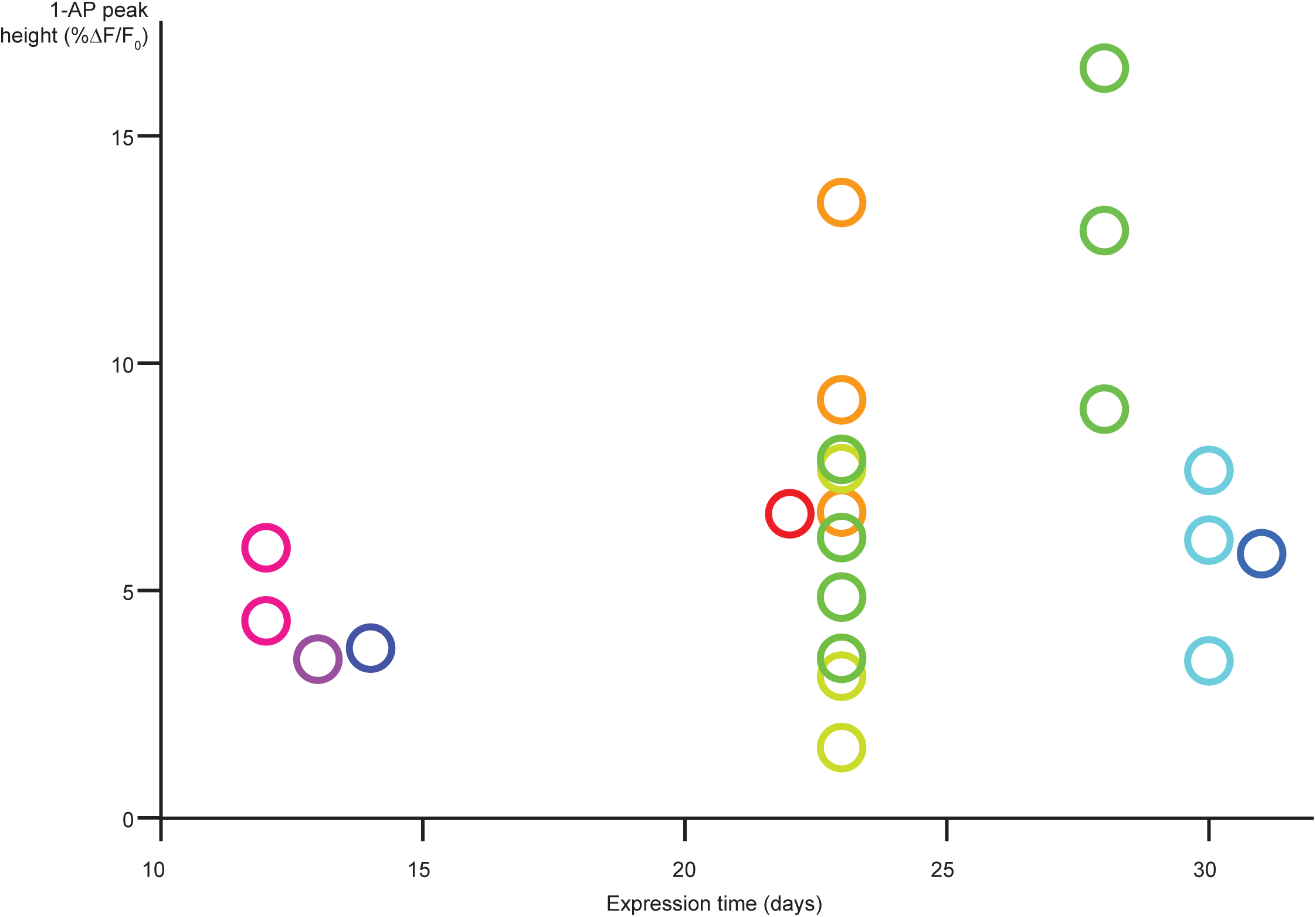
Peak GCaMP6s fluorescence increase evoked by isolated single APs as a function of time since viral injection. Each circle represents one pyramidal neuron, while each color represents one animal.

**Figure 1–Figure supplement 3.**
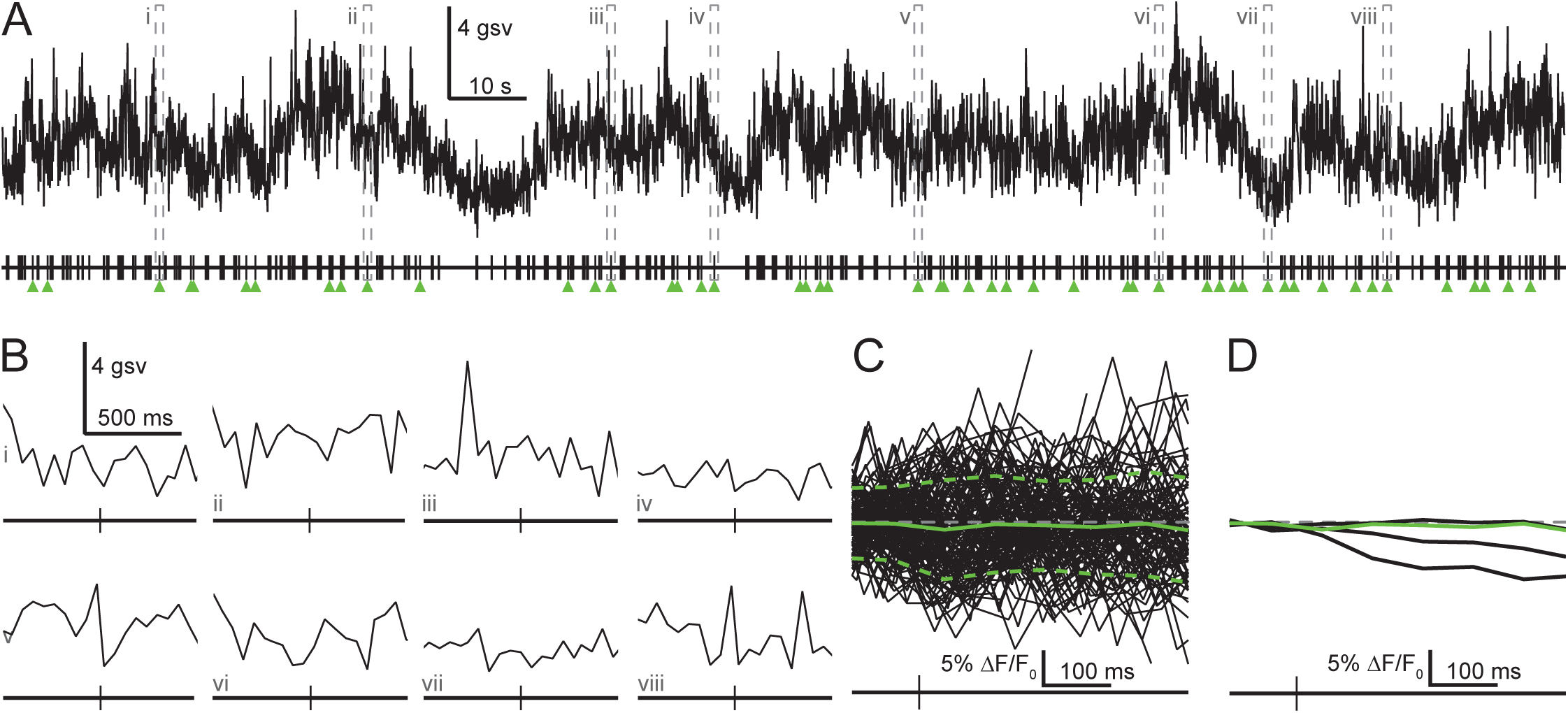
Single APs do not evoke fluorescence increases in interneurons. **(A)** GCaMP6s fluorescence (upper) and simultaneous electrical recording of APs (lower) from an interneuron in L2/3 mouse visual cortex that discharged APs at 2.6 Hz for the data shown and 2.4 Hz overall. Based on this neuron’s high firing rate, we defined its isolated single APs (green arrows) as those not preceded by other APs for 300 ms, instead of the 5.5 s required for pyramidal neurons. **(B)** Fluorescence signals recorded during 8 isolated APs within the data shown in (A). **(C)** Fluorescence change relative to baseline during all isolated APs (n = 131) for the neuron shown in (A-B). Solid and dashed green lines indicate mean and standard deviation over events; gray line indicates 0% Δ*F/F*_0_. To calculate Δ*F/F*_0_ = (*F −F*_BL_)*/F*_BL_ we estimated baseline fluorescence using the median of fluorescence values over a 60 second window around each time point, since our usual procedure of interpolating fluorescence values from periods at least 5.5 seconds after AP discharge (see methods) could not be applied due to the lack of silent periods. We also subtracted the mean Δ*F/F*_0_ value 0-200 ms before the isolated AP. While it is possible that this procedure could overestimate baseline for frequently firing neurons, resulting in incorrectly scaled Δ*F/F*_0_ values, it nonetheless shows that fluorescence does not show an average increase after single APs. **(D)** Average fluorescence change as in (C), but for all 4 interneurons (additional firing rates 12.5, 7.6 and 8.8 Hz). The green curve indicates the neuron from (A-C). Note that for the neurons with the highest spontaneous firing rates, isolated single APs occurred only after a pause in AP discharge after long periods of frequent AP discharge. As a result, fluorescence decreased after the AP due to decay from the previous period of activity.

**Figure 1–Figure supplement 4.**
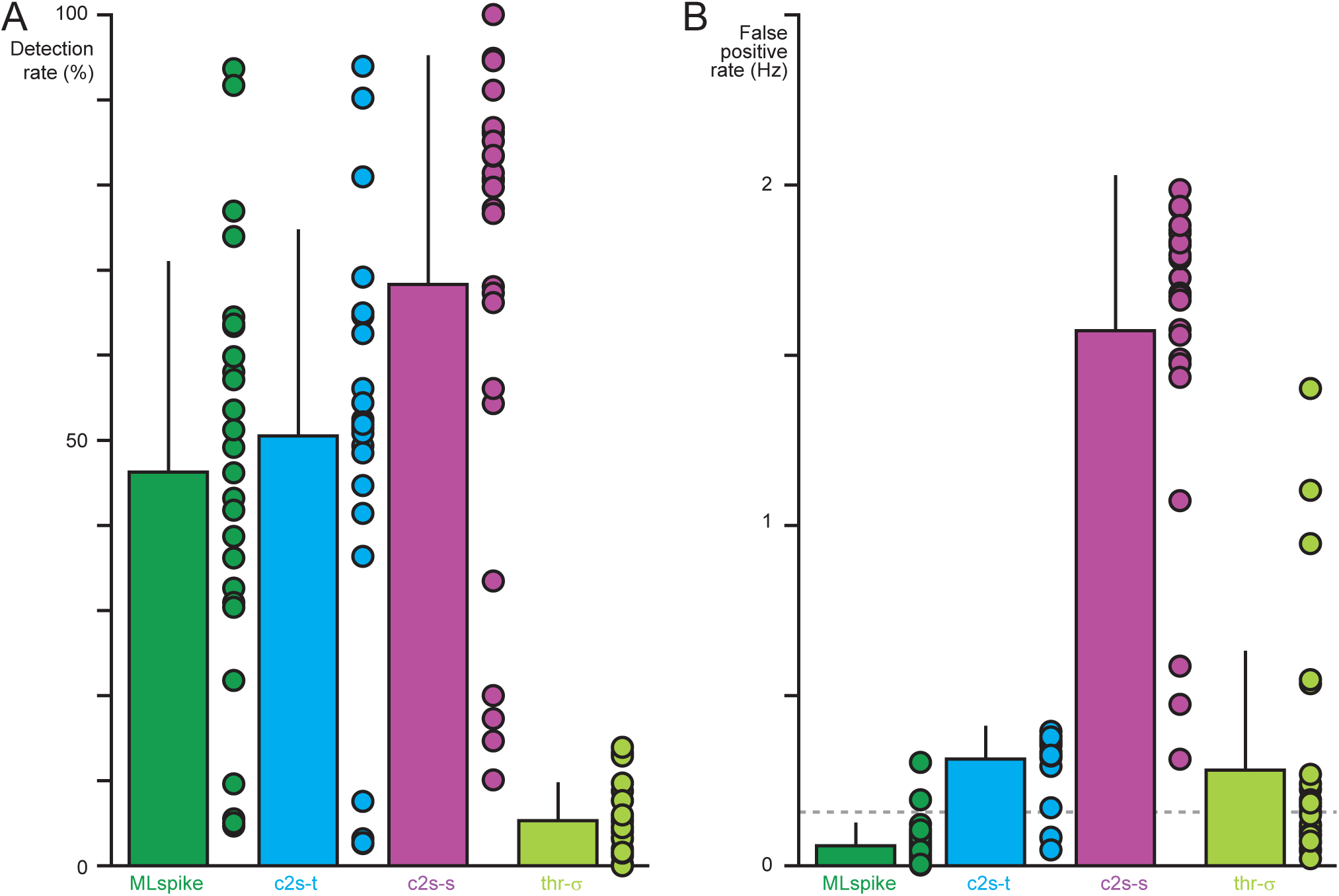
**(A)** AP detection rates for MLspike (dark green), c2s standard model (magenta), c2s re-trained (cyan) and thr-*σ* (light green). The detection rate is defined as the fraction of true APs that were successfully identified by each inference method. True and inferred APs within 100 ms were considered as matching. Bar graph shows mean and stadard deviation, while circles denote the detection rate for individual neurons. **(B)** False positive rates for the same inference algorithms. Dashed line indicates the median true firing rate (n = 26).

**Figure 2–Figure supplement 1.**
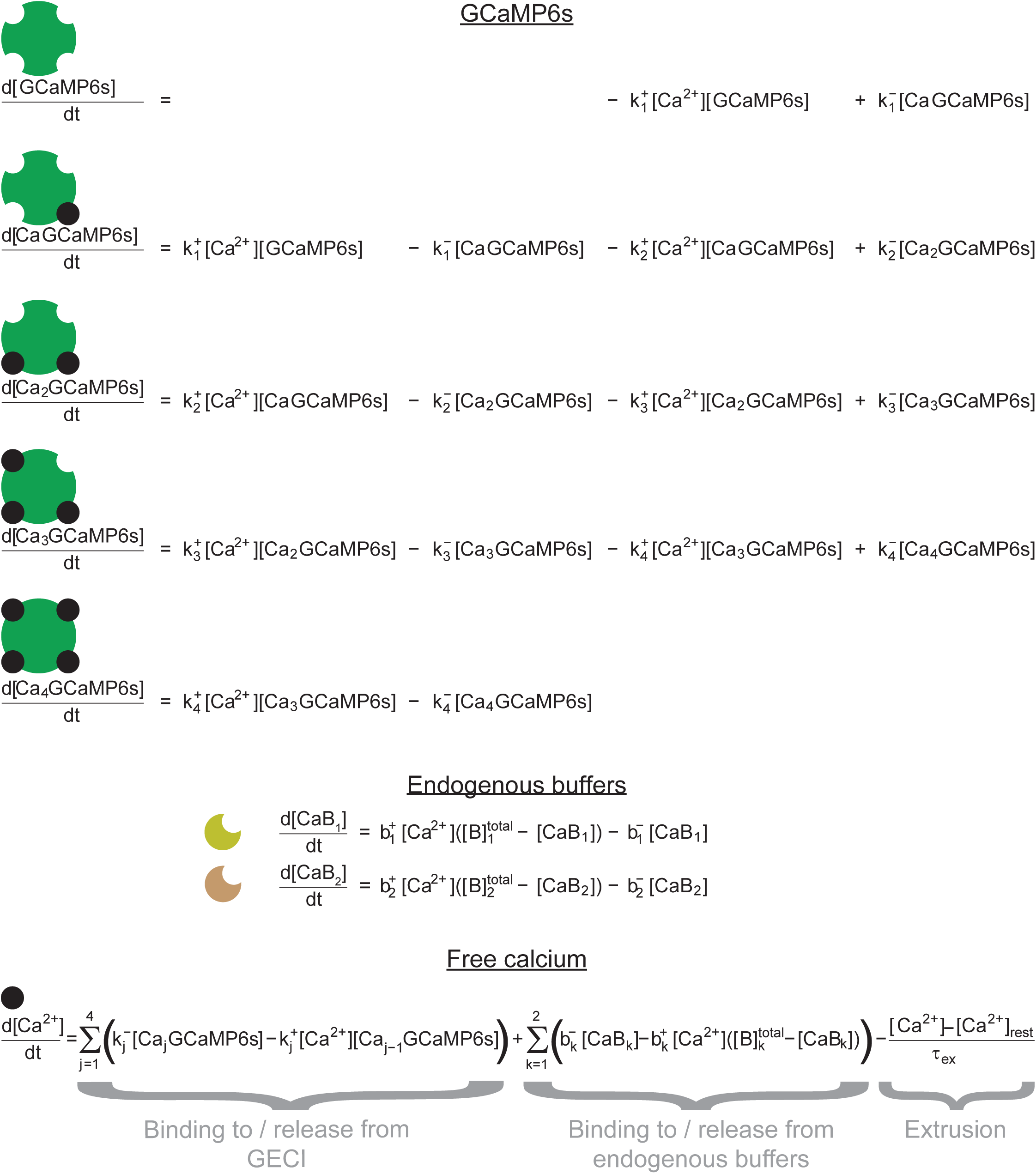
Global rate equation describing mass action kinetics and extrusion. Time derivatives of the concentrations of GCaMP6s binding states, endogenous buffer binding states and free calcium are given as instantaneous functions of the current concentrations.

**Figure 3–Figure supplement 1.**
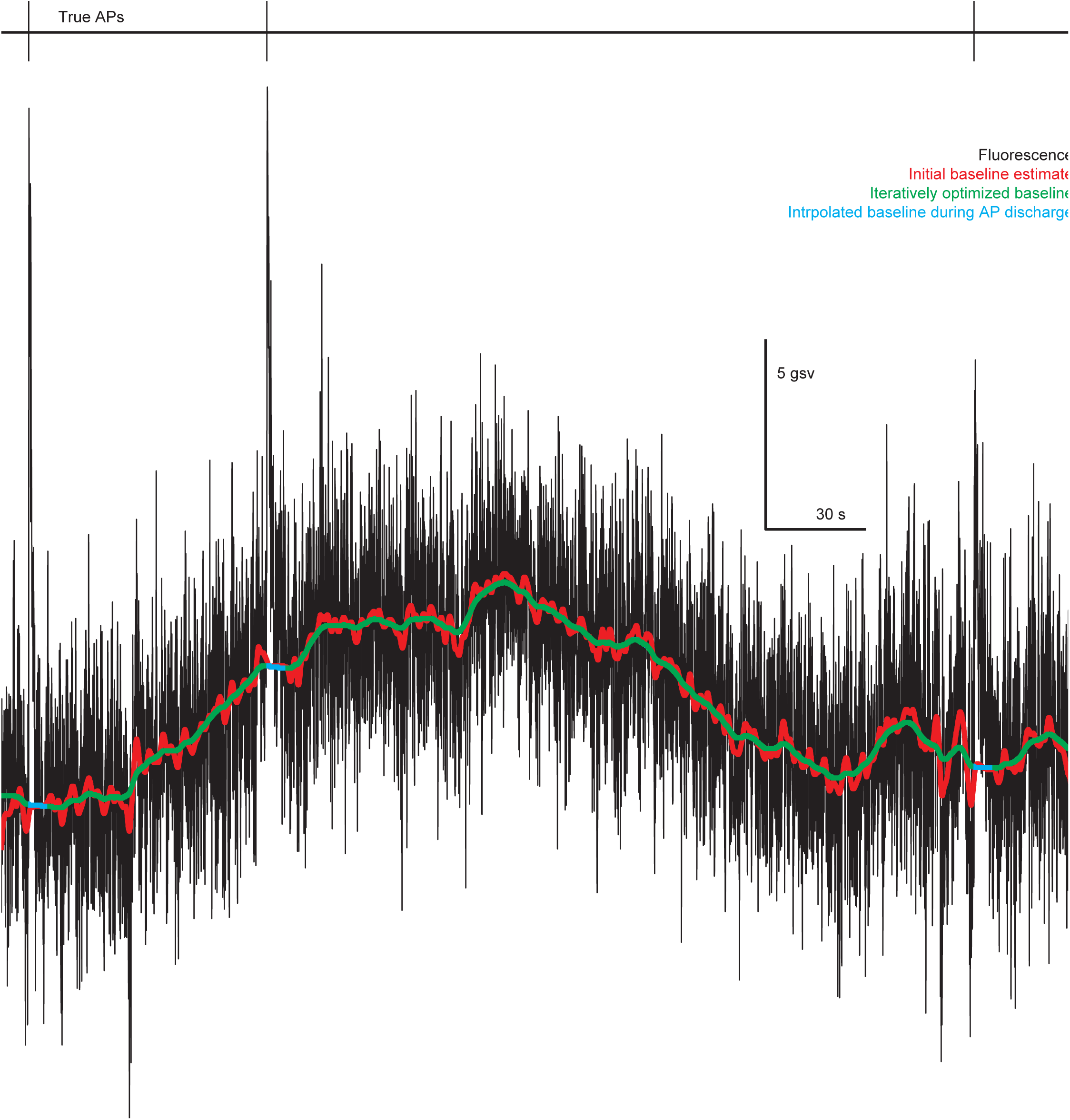
Estimation of drifting baseline fluorescence from fluorescence signals and known AP times acquired *in vivo* (full details in methods). Starting from APs (upper) and fluorescence (lower, black), an initial estimate (red) is computed by Gaussian filtering of the fluorescence data values, with fluorescence values occurring less than 5.5 seconds after an AP excluded (see methods). An iterative procedure then computes a final estimate of baseline fluorescence for periods not following an AP (green), which is interpolated to fill in the missing values following each AP (cyan).

**Figure 3–Figure supplement 2.**
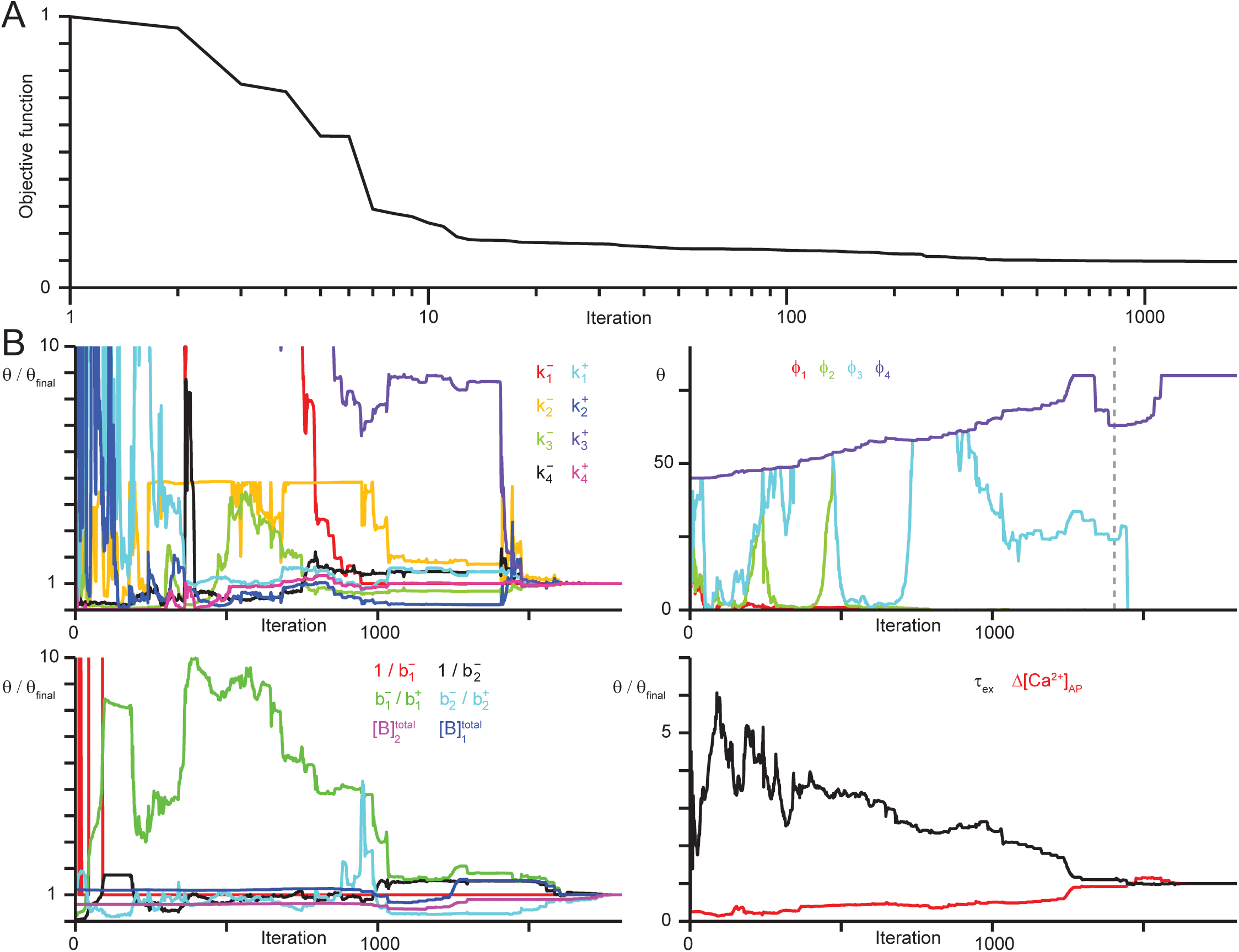
Convergence of SBM parameters while fitting *in vivo* data. **(A)** Values of the objective function *e*_vivo_ over successive iterations while fitting the SBM to full set of optical/electrical recordings (n = 22 pyramidal neurons). **(B)** Convergence of SBM parameters during optimization. *Upper left*: rate constants normalized to their final values. *Upper right*: values of *ϕ*, denoting brightness values for each binding state relative to the calcium free state. Lower left: time constants, dissociation constants and total concentrations of the two endogenous buffers, normalized to their final values. Lower right: AP-evoked calcium influx and extrusion time constant, normalized to their final values.

**Figure 3–Figure supplement 3.**
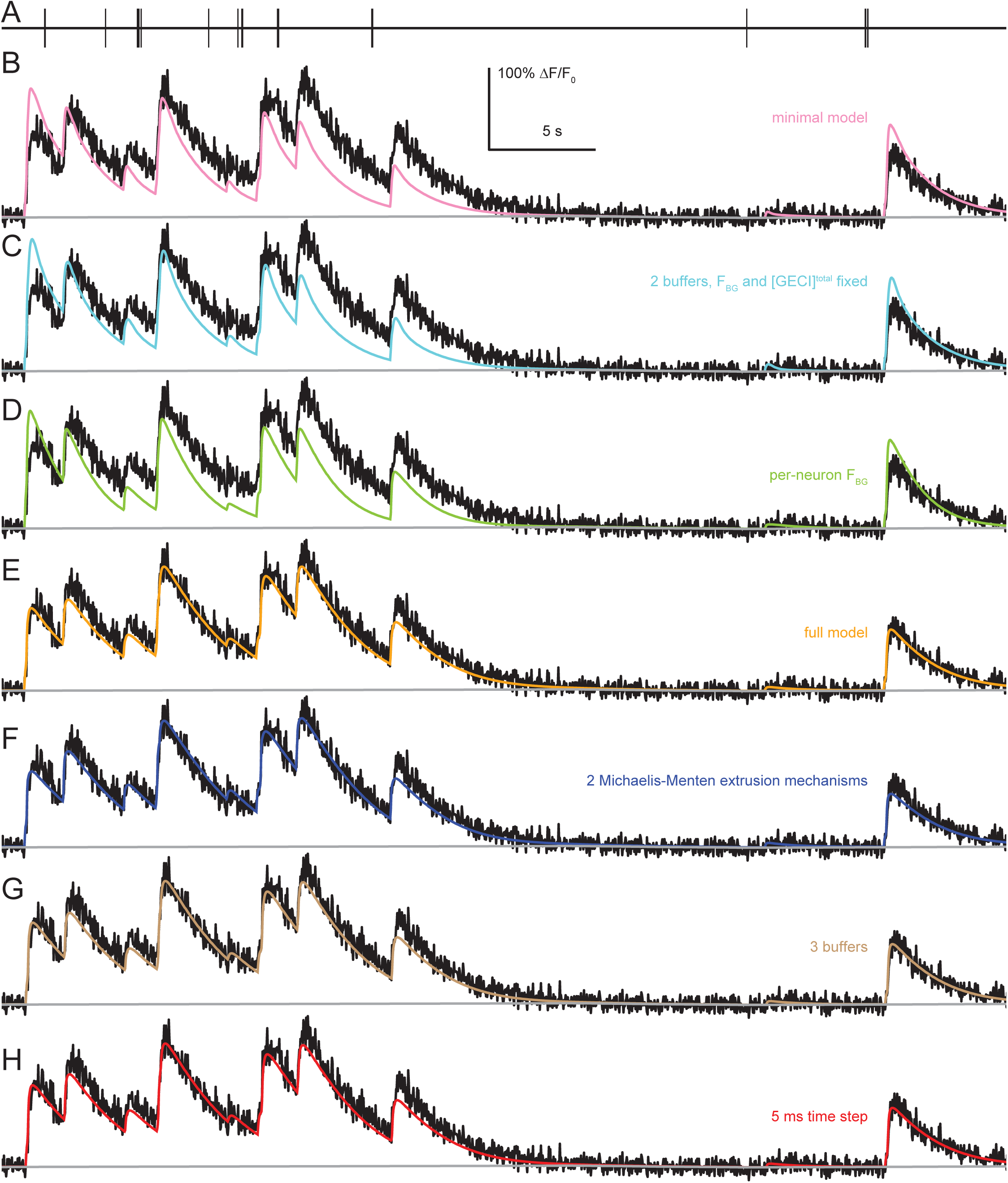
Comparison of SBM fits to data with increasing model complexity. **(A)** AP sequence recorded from a L2/3 mouse visual cortical pyramidal neuron *in vivo*. **(B)** Fluorescence (black) compared to SBM fit using a reduced minimal’ model without endogenous buffers or variation in *F*_BG_ or [GCaMP6s]^total^ over neurons. **(C)** SBM fit when including 2 endogenous calcium buffers but still without variation in *F*_BG_ or [GCaMP6s]^total^ over neurons. **(D)** SBM fit including variation in *F*_BG_ across neurons, allowing the amplitude but not the shape of AP-evoked fluorescence to be different for each neuron. **(E)** SBM fit when allowing both *F*_BG_ and [GCaMP6s]^total^ to vary over neurons; this is the SBM variant used throughout the present study unless otherwise noted. **(F)** Fit for extended SBM, consisting of the full SBM in (E) along with 2 Michaelis-Menten extrusion mechanisms, allowing for calcium-dependent modulation of extrusion rate. **(G)** Fit for extended SBM, consisting of the full SBM in (E) but with 3 endogenous buffers instead of 2. **(H)** Fit for the full SBM shown in (E), but when solving the rate equation with a maximum time step of 5 instead of 10 ms.

**Figure 3–Figure supplement 4.**
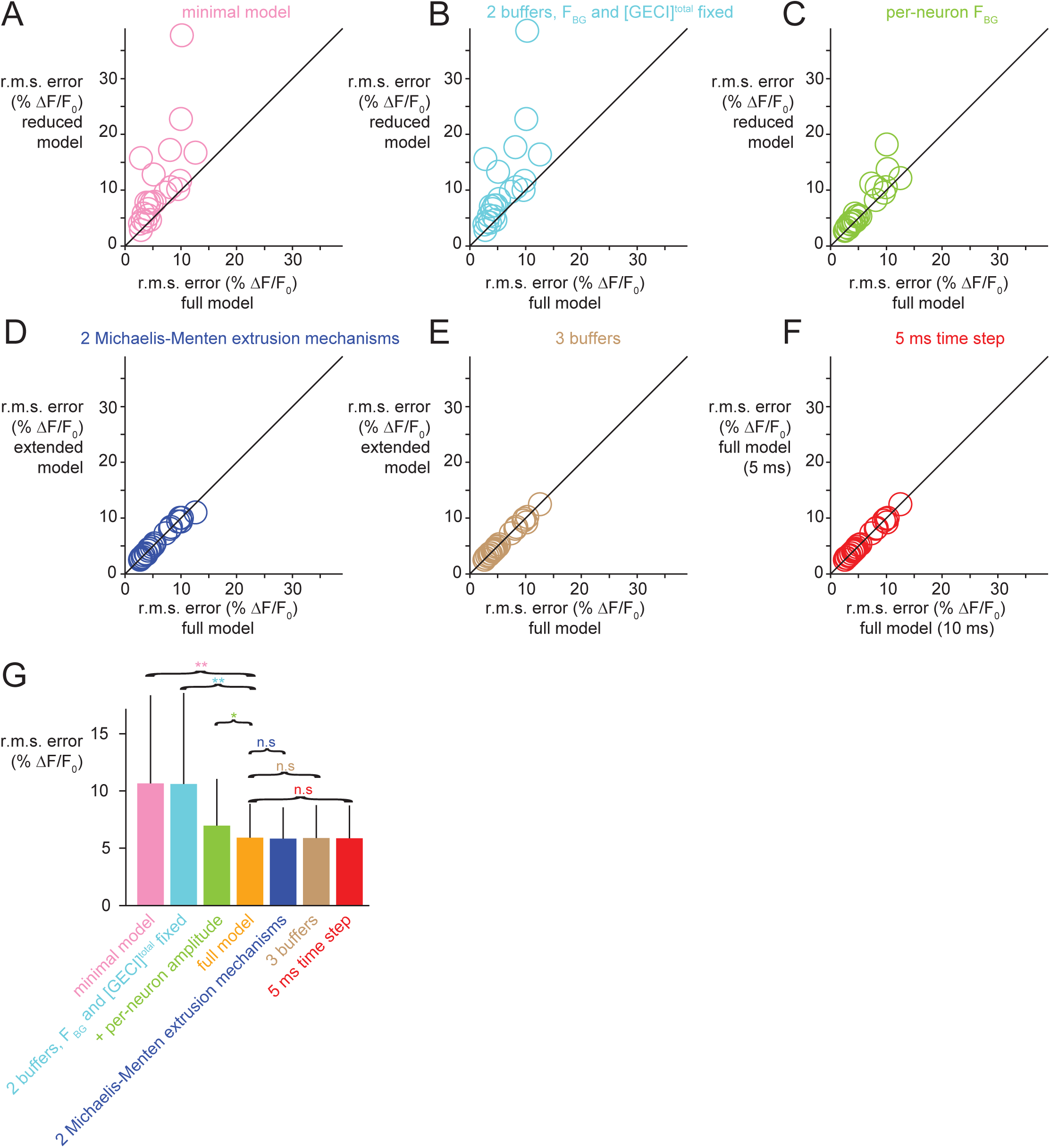
Comparison of SBM variants over pyramidal neurons (n = 22) shows that the standard SBM fits the data better than simplified versions, but using more complicated variants do not improve fit quality. **(A)** Root-mean-square error for the standard SBM compared to a reduced ‘minimal’ version without buffering or variation in *F*_BG_ or [GCaMP6s]^total^ over neurons (Figure 3 - figure supplement 2b). **(B)** Comparison of standard SBM to reduced version with 2 endogenous calcium buffers but no variation in *F*_BG_ or [GCaMP6s]^total^ over neurons (Figure 3 - figure supplement 2c). **(C)** Comparison of SBM to reduced version which also includes variation in *F*_BG_ but not [GCaMP6s]^total^ over neurons (Figure 3 - figure supplement 2d). **(D)** Comparison of standard SBM to an extended version using two Michaelis-Menten extrusion mechanisms, each with its own Michaelis constant and maximum rate (see methods). **(E)** Comparison o f standard SBM to an extended version with 3 endogenous buffers (each with its own concentration, affinity and time constant). **(F)** Comparison of standard SBMs with maximum time steps of 5 vs. 10 ms. **(G)** Mean and standard deviation of root-mean-square error over neurons for each SBM variant. Single asterisk indicates p < 0.01, double asterisks indicate p < 1e-6.

**Figure 3–Figure supplement 5.**
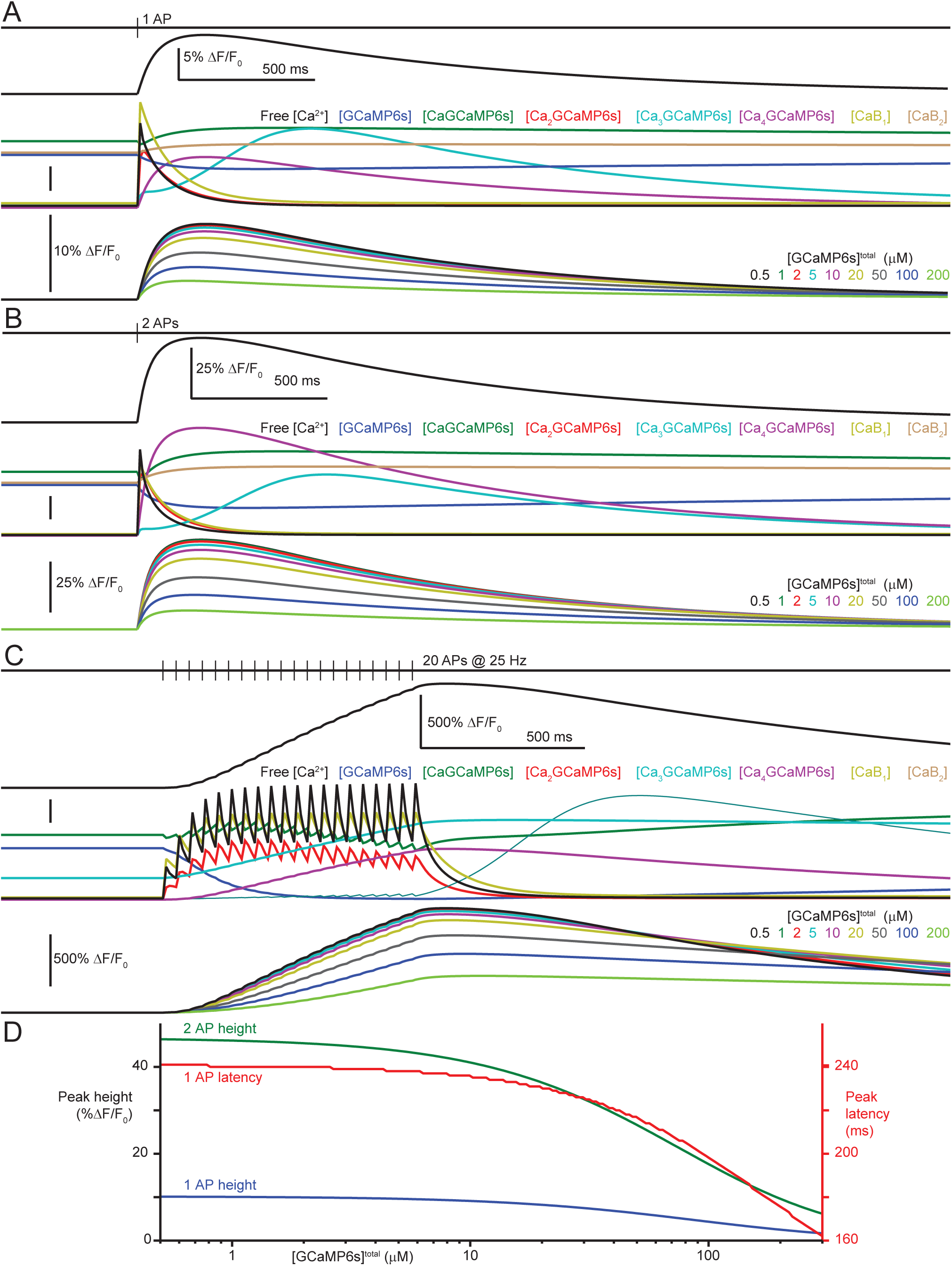
SBM simulations using parameters fit to *in vivo* data 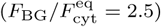. **(A)** Fluorescence response (upper) to one AP using [GCaMP6s]^total^ = 5 *µ*M, with binding state concentrations over time (middle, scalebar: 400 nM Ca^2+^, 1 *µ*M GCaMP6s, 1 *µ*M CaGCaMP6s, 100 nM Ca_2_GCaMP6s, 0.4 nM Ca_3_GCaMP6s, 10 nM Ca_4_GCaMP6s, 4 *µ*M B_1_, 4 *µ*M B_2_). Fluorescence response to one AP as a function of [GCaMP6s]^total^ (lower). **(B)** As in (A) but for a 2-AP burst (scalebar: 1 *µ*M Ca^2+^, 1 *µ*M GCaMP6s, 1 *µ*M CaGCaMP6s, 200 nM Ca_2_GCaMP6s, 2 nM Ca_3_GCaMP6s, 20 nM Ca_4_GCaMP6s, 10 *µ*M B_1_, 4 *µ*M B_2_). **(C)** As in (A-B) but for 20 APs at 25 Hz (scalebar: 1 *µ*M Ca^2+^, 1 *µ*M GCaMP6s, 1 *µ*M CaGCaMP6s, 400 nM Ca_2_GCaMP6s, 20 nM Ca_3_GCaMP6s, 1 *µ*M Ca_4_GCaMP6s, 10 *µ*M B_1_, 10 *µ*M B_2_). **(D)** Simulated peak fluorescence evoked by one (blue) and two APs (green) and single-AP peak latency (red) as functions of [GCaMP6s]^total^.

**Figure 3–Figure supplement 6.**
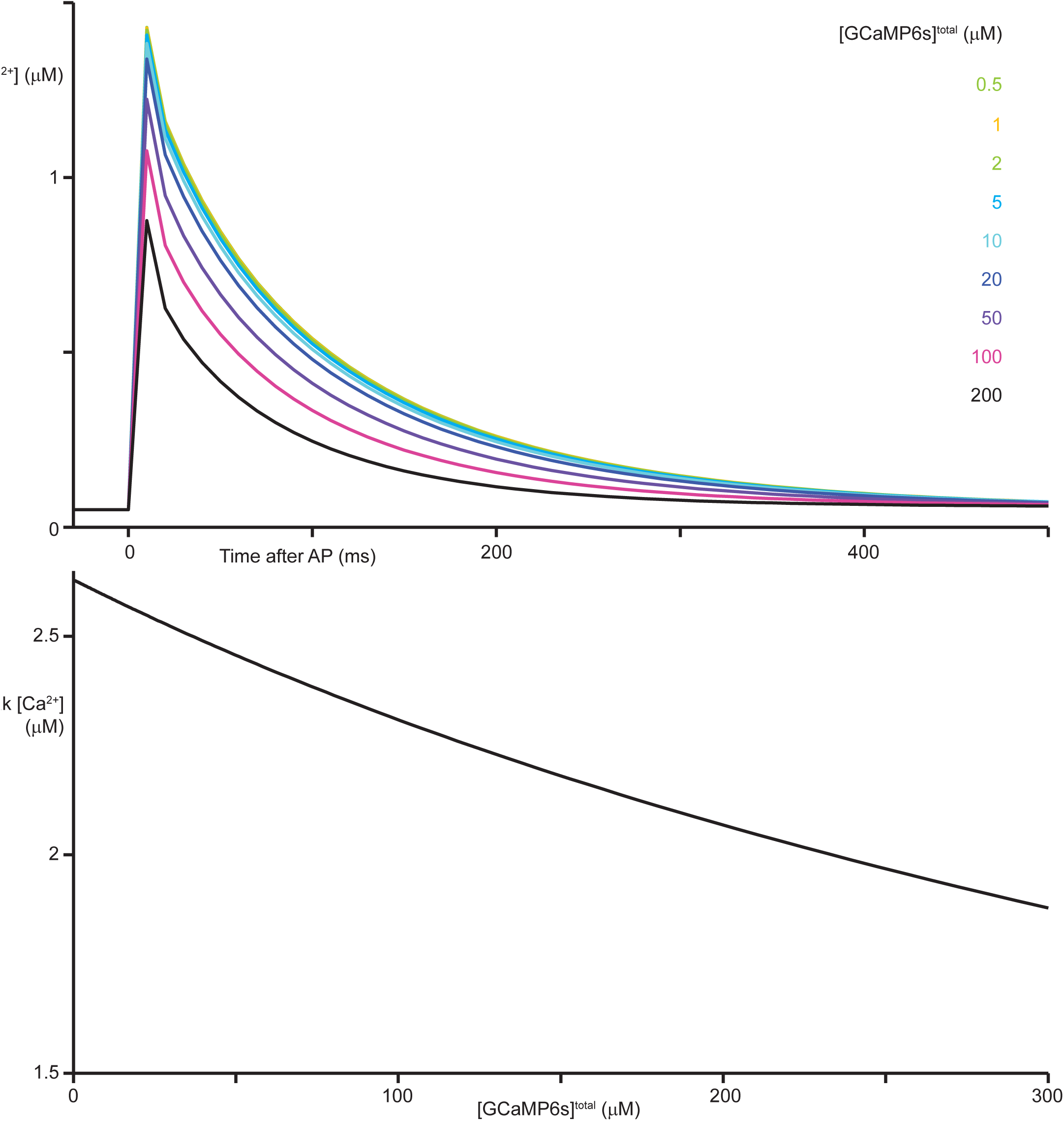
The concentration of free calcium over time after AP discharge depends on total GCaMP6s concentration in SBM simulations. **(A)** SBM simulation of calcium concentration after a single AP as a function of total GCaMP6s concentration. At low concentrations GCaMP6s does not contribute significantly to calcium buffering compared to endogenous buffers, while at higher concentrations it acts as a buffer, reducing the free calcium concentration in the cytosol. **(B)** Peak [Ca^2+^] one time step (*δ ≤* 10 ms) after a single AP as a function of total GCaMP6s concentration.

**Figure 3–Figure supplement 7.**
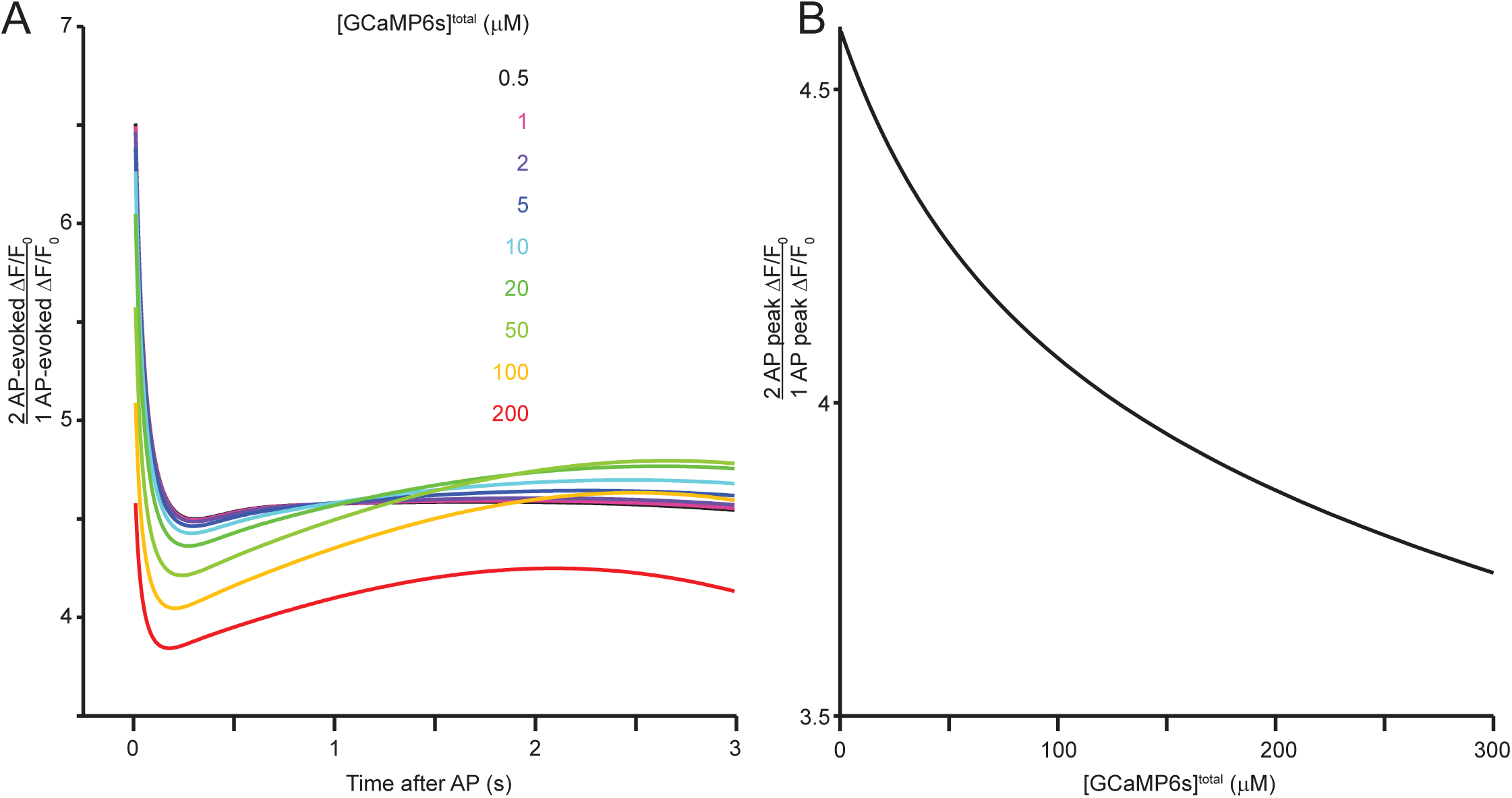
Nonlinearity of AP-evoked fluorescence depends on total GCaMP6s concentration in SBM simulations 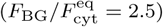. **(A)** 2-AP fluorescence response divided by 1-AP response, as a function of time after AP discharge and for a range of total GCaMP6s concentrations. Note that the ratio of the 2-AP response to the 1-AP response is not constant over time for any GCaMP6s concentration, and that the shape of the ratio as a function of time depends on the GCaMP6s concentration. **(B)** Ratio of peak 2-AP evoked fluorescence to peak 1-AP fluorescence as a function of GCaMP6s concentration.

**Figure 3–Figure supplement 8.**
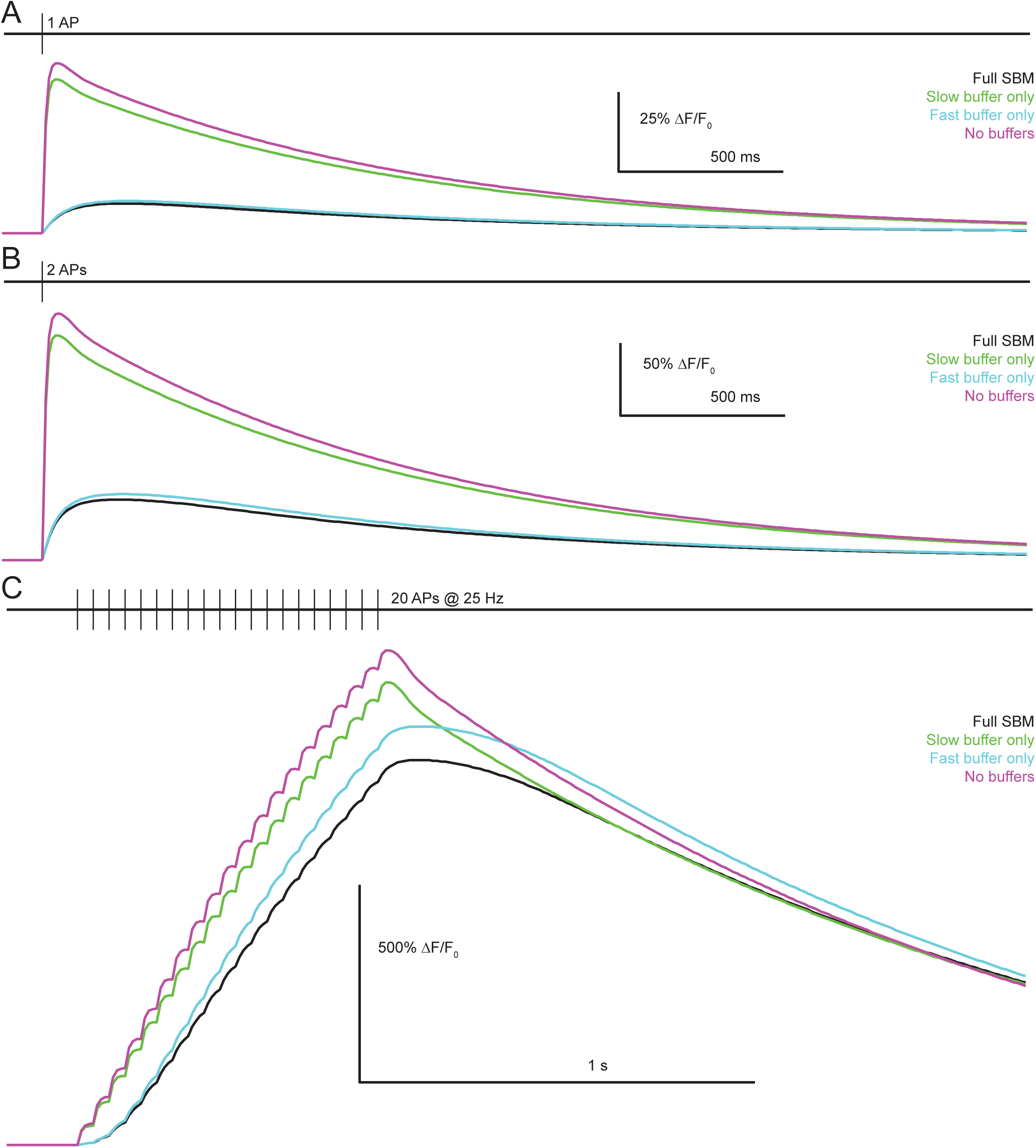
Role of endogenous buffers in shaping AP-evoked fluorescence in SBM simulations ([GCaMP6s]^total^ = 5 µM,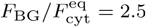. **(A)** Fluorescence arising from discharge of one AP in simulations from the full SBM (black) or when incorporating only the slow buffer (green), only the fast buffer (cyan) or neither buffer (magenta). **(B)** As in (A), but for a burst of 2 APs. **(C)** As in (A-B), but for a train of 20 APs at 50 Hz. Note that the slow buffer plays only a minor role in determining fluorescence responses to discharge of 1 or 2 APs, while the fast buffer has no effect on the decay phase following the train of 20 APs.

**Figure 3–Figure supplement 9.**
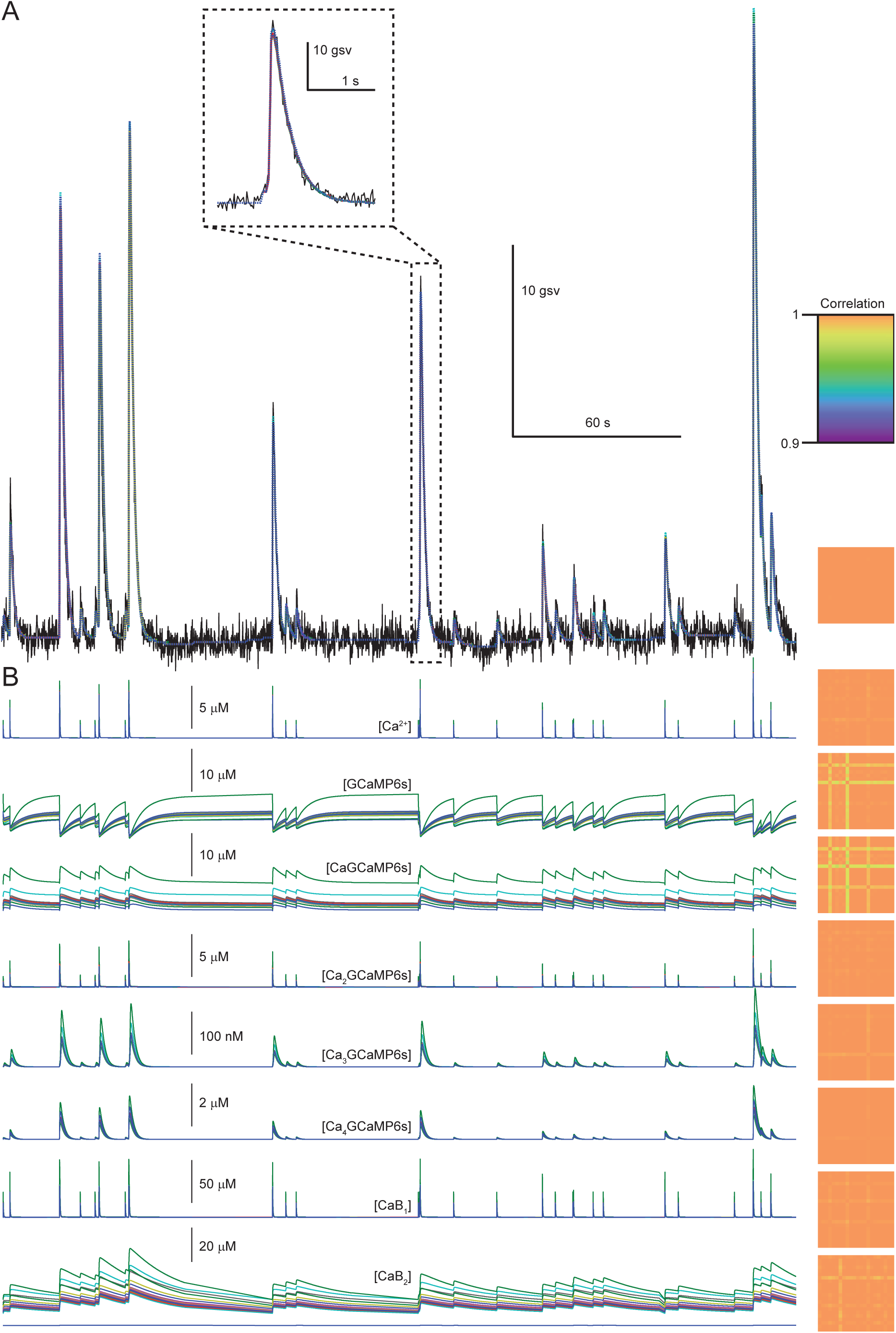
Variation of SBM fluorescence predictions over multiple parameter sets. **(A)** *Left*: Fluorescence (black, same data as Figure 4a-c) with predictions from each parameter set (colored dashed lines). Each set of SBM parameters was obtained by fitting the *in vivo* dataset while excluding a single neuron. *Right*: Correlation matrix showing correlation over time of fluorescence predictions from the 22 parameter sets. **(B)** Binding state concentrations from the fits in (A) (as in Figure 3b-c), with correlation over time of binding state concentrations from the 22 parameter sets.

**Figure 4–Figure supplement 1.**
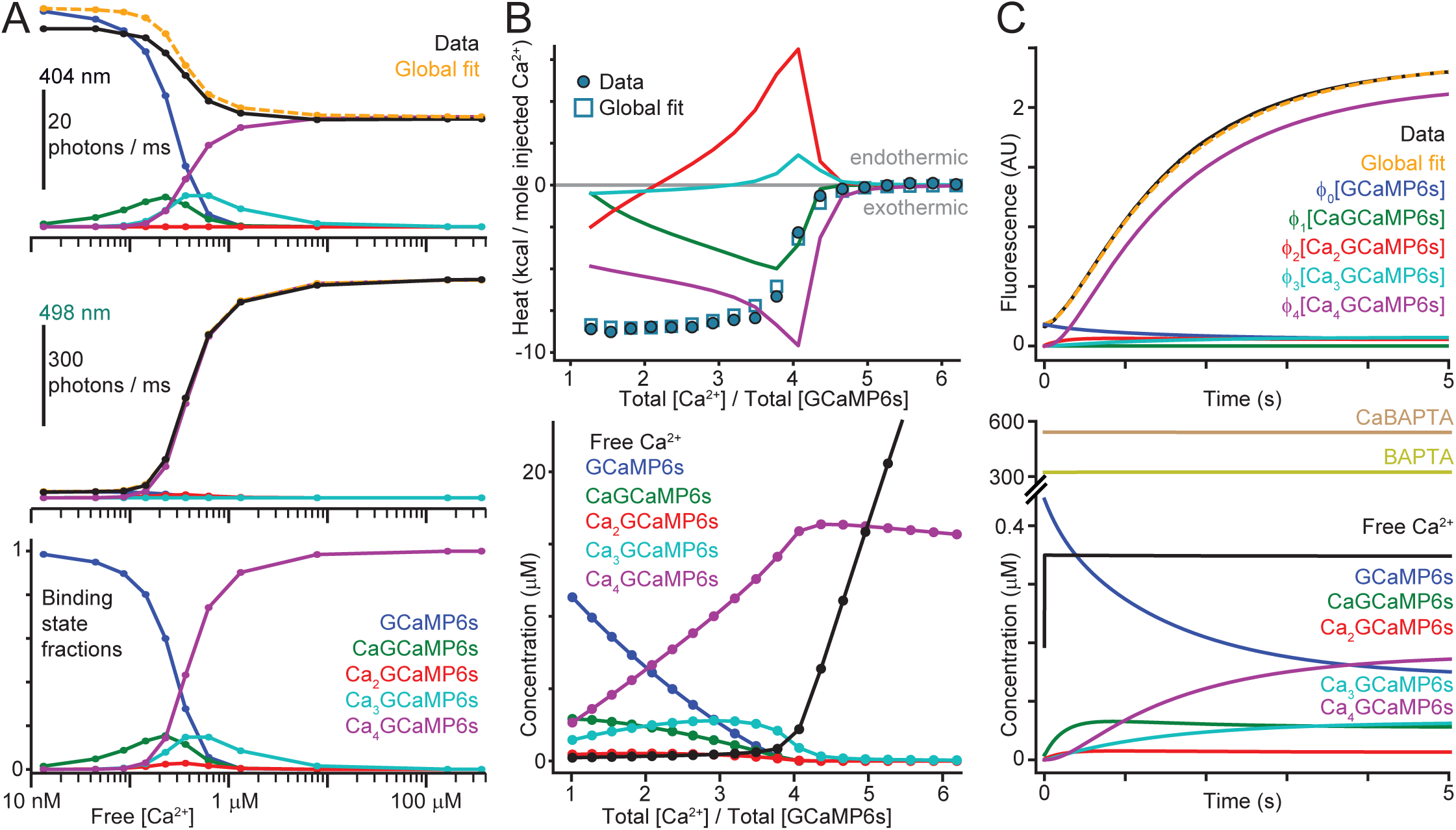
Decomposition of *in vitro* binding assay data into contributions from each GCaMP6s binding state. **(A)** Fluorescence (black) excited at 404 nm (upper) and 498 nm (middle) as a function of free calcium concentration at each step of a titration used to measure excitation spectra (Figure 4a,d), and prediction fluorescence from SBM global fit (orange). Additional colors show contributions of each binding state to total predicted fluorescence; excitation wavelengths are shown as arrowheads in Figure 4c. Modeled binding state fractions (lower) as a function of free calcium for the same data. **(B)** Integrated peak heats (circles, upper) for the data shown in turqouise in Figure 4b,d along with global fit (squares) and contributions to enthalpy changes arising from changes in each binding state concentration (colored curves). Modeled concentrations of free Ca^2+^ (black, lower) and each GCaMP6s binding state are shown for the same data. Free Ca^2+^ increases slowly until the four binding sites of GCaMP6s are saturated, after which it increases rapidly. The concentration of the saturated state Ca_4_GCaMP6s initially increases due to binding, then decreases due to dilution and perfusion. **(C)** Upper graph: Fluorescence signals acquired during stopped-flow experiment (black) and global fit (orange) for the data in Figure 4f showing binding kinetics after a transition from 17 nM to 348 nM free Ca^2+^. Additional colors show the modeled fluorescence contributions of each GCaMP6s binding state. Lower graph: modeled concentrations over time for all molecular species.

**Figure 5–Figure supplement 1.**
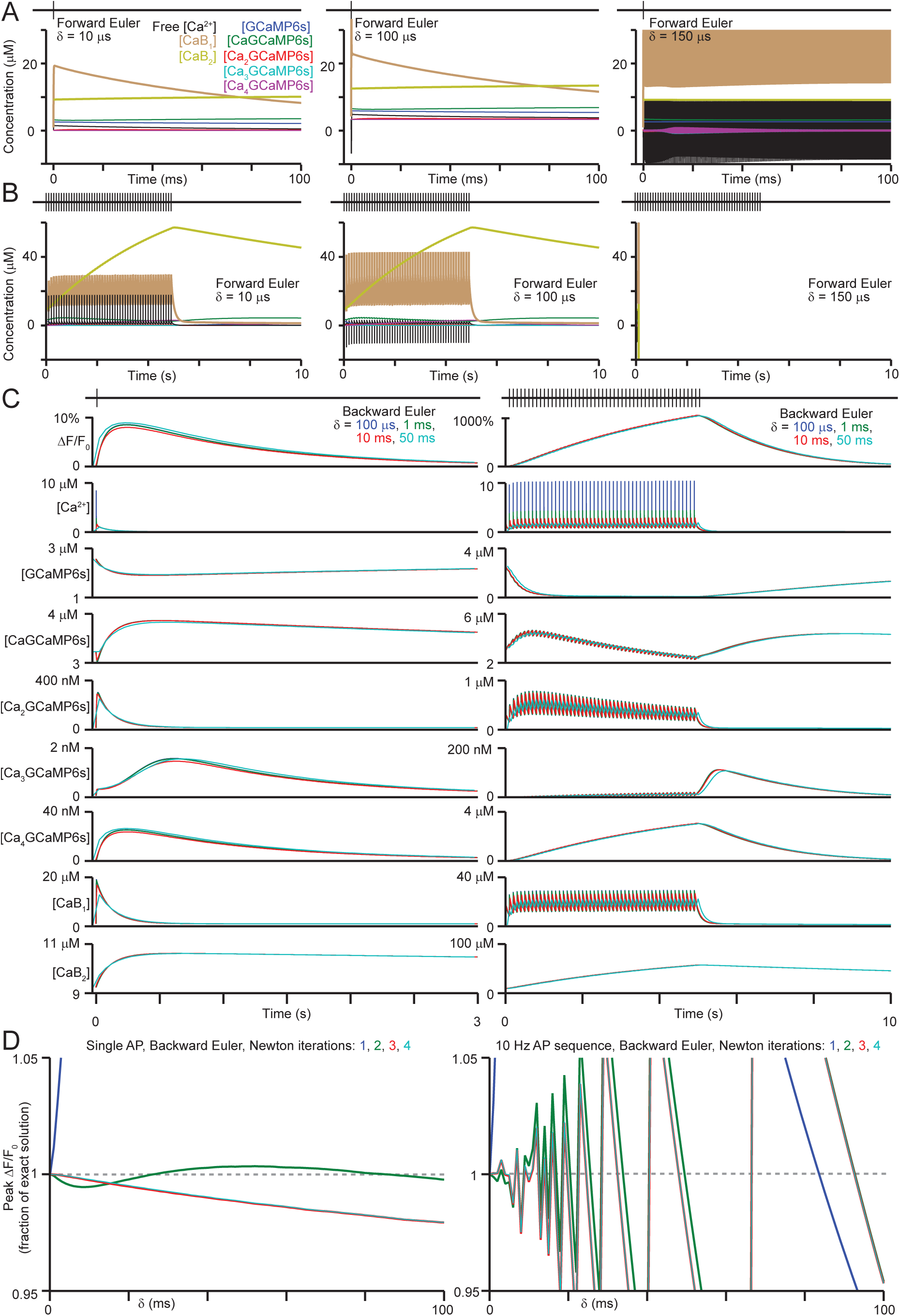
**(A)** Forward (explicit) Euler integration of the SBM rate equation for 1 AP, with δ = 10 (left), 100 (center) or 150 µ s. Steps >50 µ s resulted in negative concentrations. **(B)** As in (A), but for 50 APs at 10 Hz. Steps >100 *µ s* caused divergence to infinity. **(C)** Backward Euler integration (see methods) for the AP sequences in (A-B) with 3 Newton iterations and δ = 0.1, 1, 10 and 50 ms. Note the close agreement for δ ≤ 10 ms, and that transient differences in [Ca ^2^+] for δ ≤ 10 ms do not lead to differences in binding state concentrations or predicted fluorescence. **(D)** Peak δF/F0 values for the AP sequences in (A-C), relative to the exact solution and as a function of δ for backward Euler integration, with 1-4 Newton iterations. Note the nearly identical solutions for 3 (the default value for model fitting and AP inference) *vs*. 4Newton iterations.

**Figure 5–Figure supplement 2.**
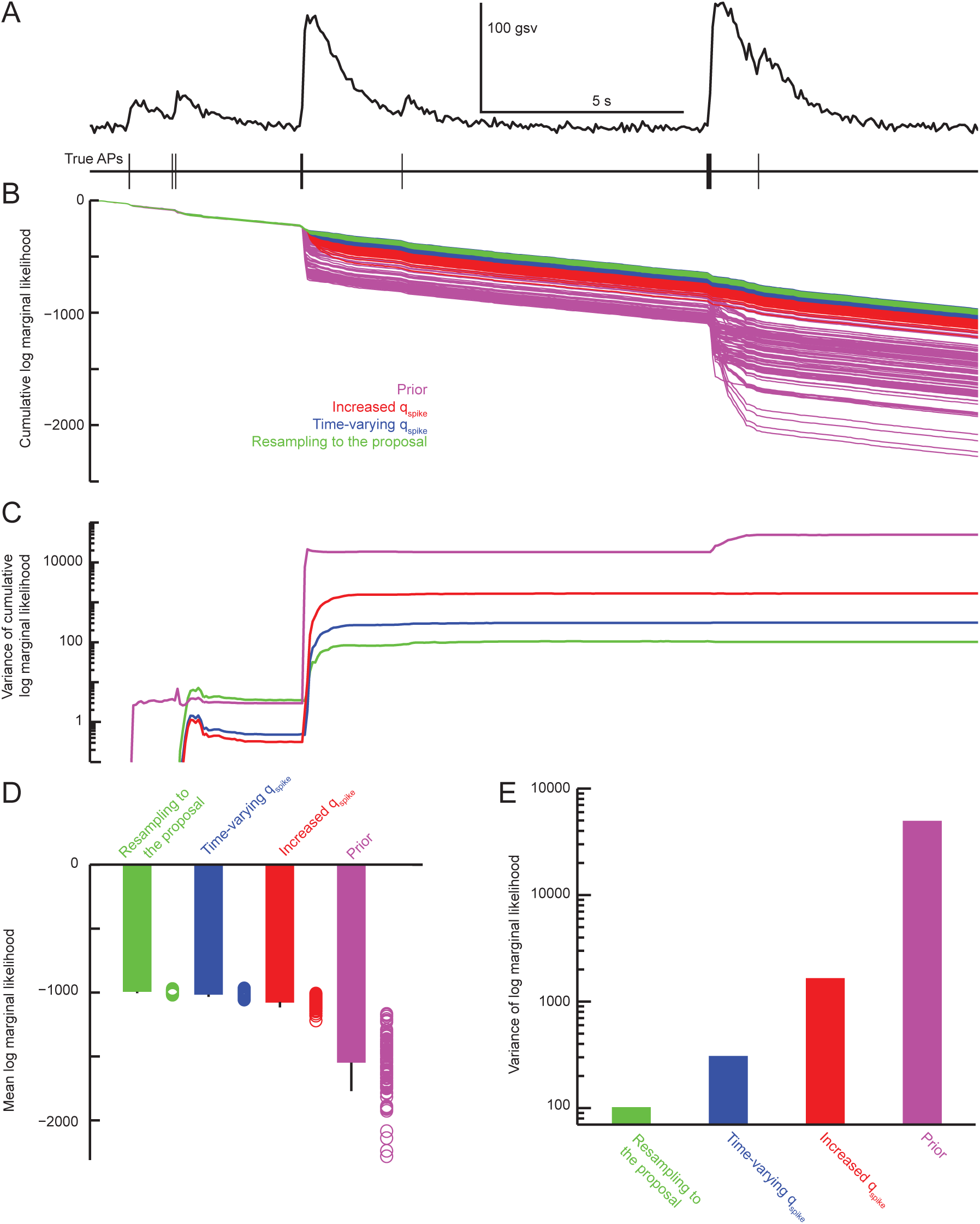
Sampling and resampling techniques for reducing the variability of the SMC algorithm’s output. **(A)** Fluorescence and APs recorded for 22 seconds from a pyramidal neuron in L2/3 mouse visual cortex. **(B)** Cumulative log marginal likelihood calculated for the data in (A). At each time point the SMC technique (see methods) is used to calculate the likelihood given the model parameters of all observed data up to the current time point. Results are shown for 100 independent runs using the same parameters and data, with sampling of AP sequences from the prior distribution (magenta), a proposal distribution with increased AP discharge probability (red), a time varying proposal in a two-round scheme (blue) or a time varying proposal with resampling to the proposal (green). **(C)** Variance over runs of the cumulative log marginal likelihoods shown in (B). **(D)** Distribution over runs of marginal likelihood at the end of the data in (A-C) (circles). Bar graph shows means and standard deviations. **(E)** Variance over runs for the distributions in (D).

**Figure 5–Figure supplement 3.**
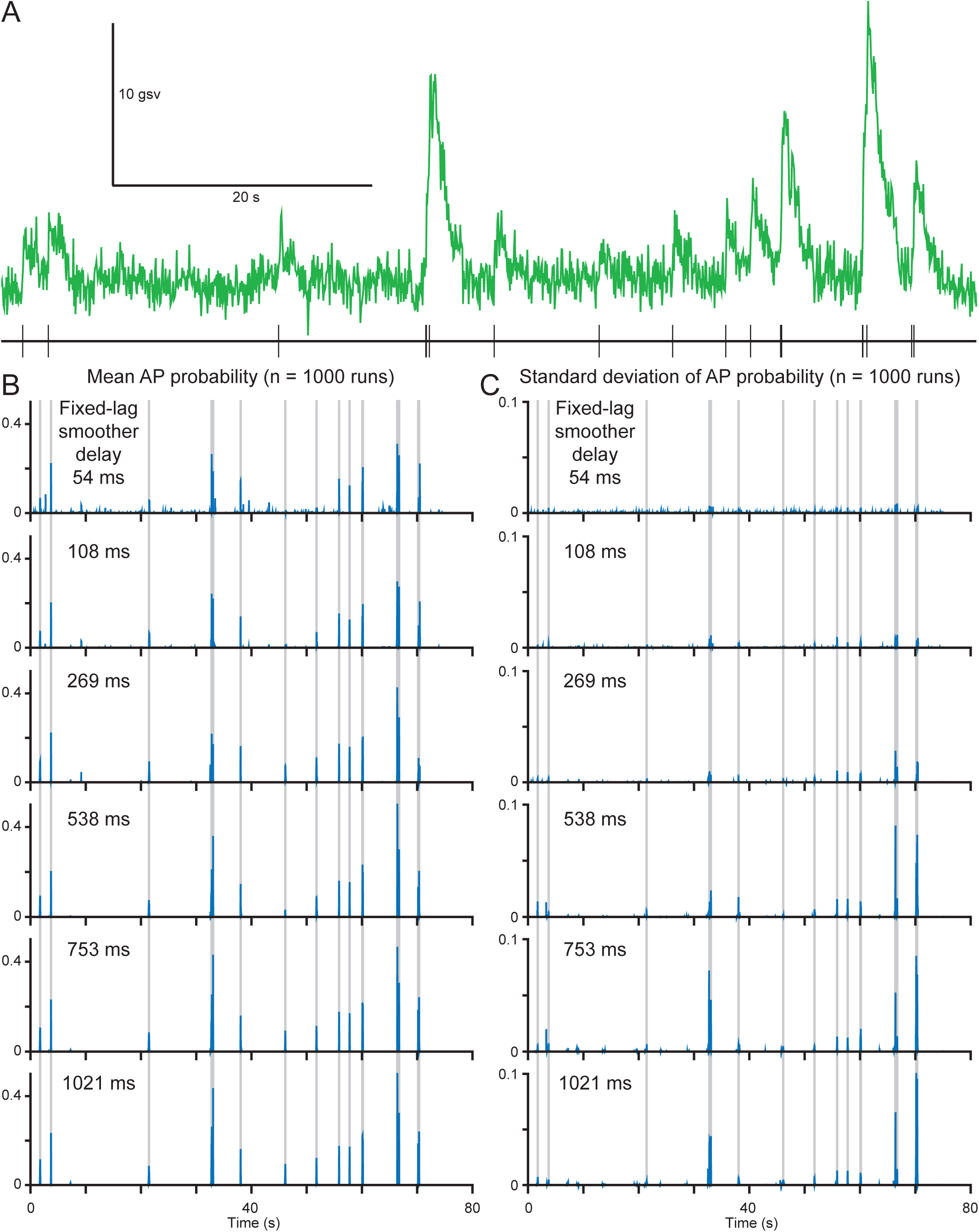
Effect of the fixed-lag smoother delay on inferred AP discharge probability using the SMC algorithm with the SBM. **(A)** Fluorescence signals (upper) and electrically recorded APs (lower) from a L2/3 mouse visual cortical pyramidal neuron. **(B)** Probability of AP discharge every 10 ms, inferred using the filter smoother and averaged over 1000 runs of the algorithm. The SMC algorithm was used with 102400 particles. Results are shown for fixed-lag smoother delays 54 (top), 108, 269, 538, 753 and 1021 ms (bottom). Same data as in (A); gray lines indicate true AP times. **(C)** As in (B), but showing the standard deviation over 1000 runs of the algorithm.

**Figure 5–Figure supplement 4.**
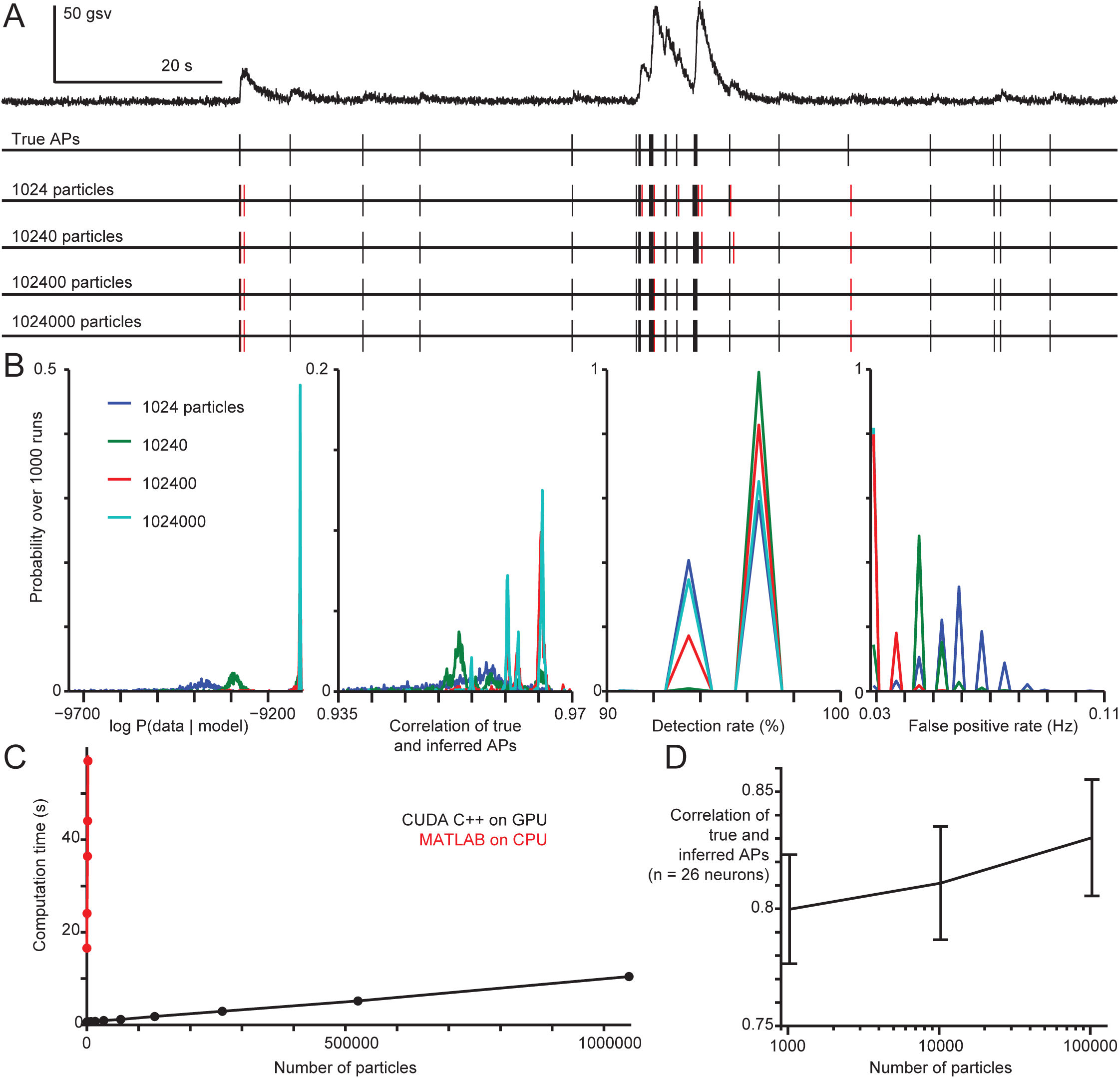
Accuracy and speed of SMC/SBM-based AP inference as a function of particle count. **(A)** 134 seconds of fluorescence signals (upper) and simultaneously recorded APs for a L2/3 mouse visual cortical pyramidal neuron. SBM/SMC-based AP inference results are shown for a single run of the algorithm with 1024, 10240, 102400 and 1024000 particles. False positives produced by the algorithm (time difference from true APs > 200 ms) are shown in red. **(B)** Probability distributions over 1000 runs for the marginal data likelihood (see methods), correlation of true and inferred AP (smoothing *σ* 200 ms), detection rate and false positive rate (timing tolerance 200 ms). Results are shown for the same particle counts and data as in (A). **(C)** Computation time for a single SMC forward pass with calculation of posterior probability of AP discharge every 10 ms, as a function of the number of particles and for the same data as in (a-b). Results are shown for a CUDA-C++ implementation running on a Geforce GTX 1080 TI (nVidia) or a Matlab implementation running on an Ryzen 7 1800X 8 core processor (AMD). **(D)** Correlation of true and inferred APs over our entire *in vivo* dataset (n = 26 neurons) as a function of particle count. Error bars show standard error of the mean over 1000 runs.

**Figure 5–Figure supplement 5.**
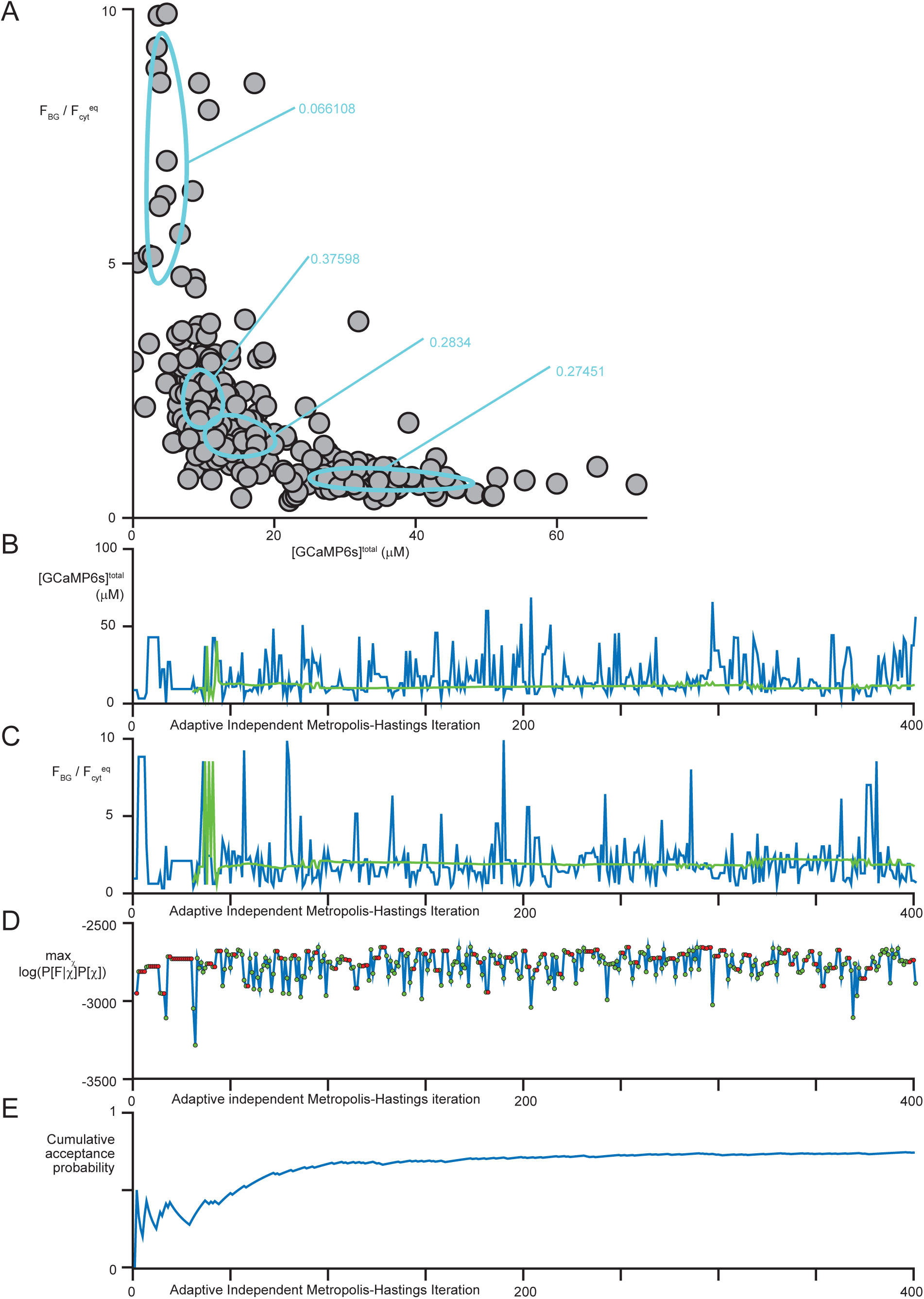
Identification of per-neuron parameters from fluorescence data alone. **(A)** Scatter plot of [GCaMP6s]^total^ vs. 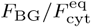 for adaptive independent Metropolis-Hasting procedure samples (see methods). Contours show 1-s.d. isoclines from each mixture cyt component; numbers indicate component probabilities. **(B)** [GCaMP6s]^total^ concentration for each sample in (A) (blue) and mode of the current iteration’s lognormal mixture fit (green). **(C)** As in (B), but for 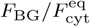. **(D)** Log marginal data likelihood times the prior probability, after optimization with respect to the firing rate, for the samples in (A-C). Green dots indicate accepted samples, while red dots indicate rejections with repetition of the previous sample. **(E)** Overall fraction of samples that have been accepted as a function of the number iterations completed.

**Figure 5–Figure supplement 6.**
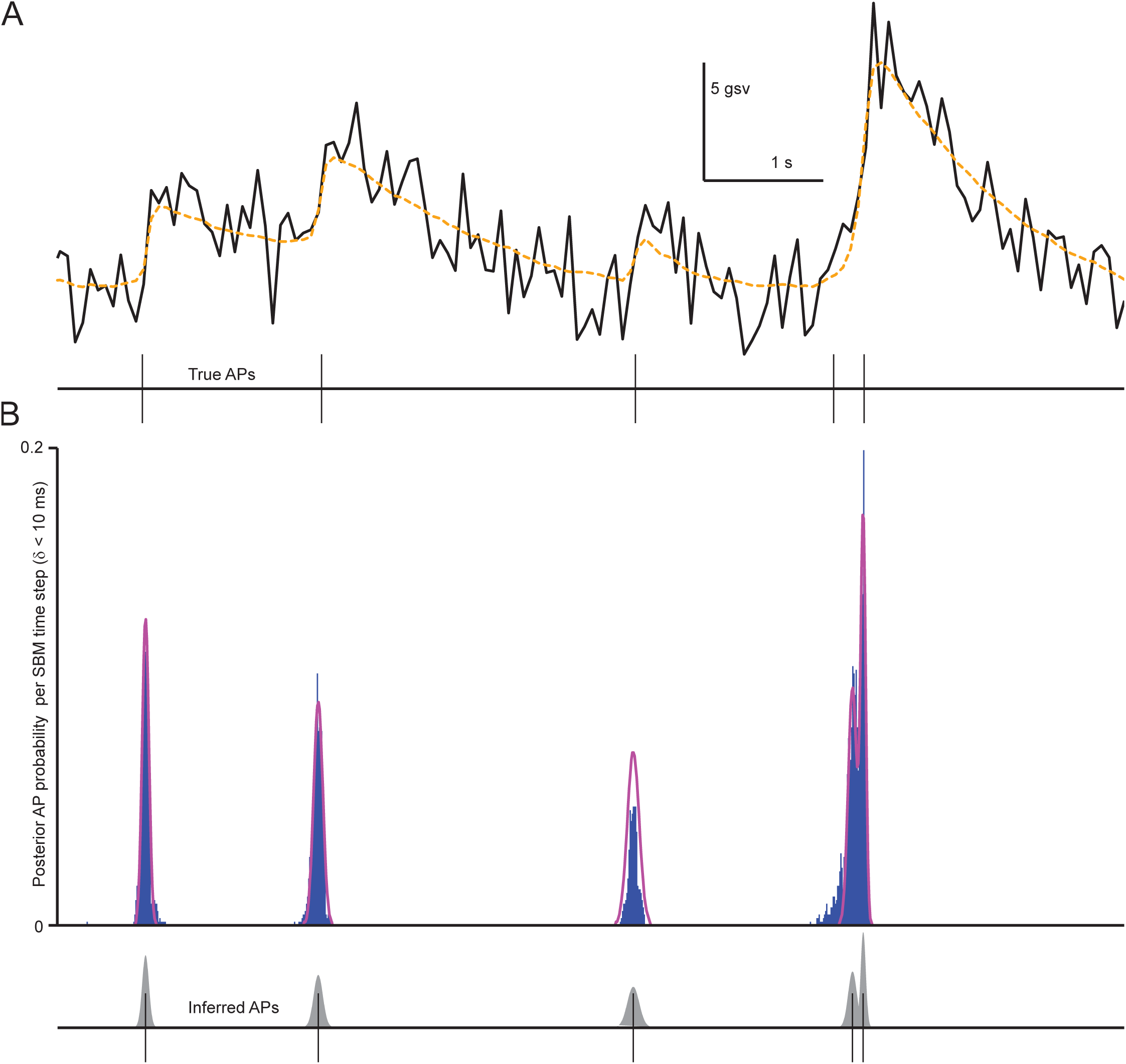
A single AP sequence is fit to posterior probability of AP discharge every time step (*δ ≤* 10 ms) given fluorescence data. **(A)** Neuronal fluorescence (upper, black) with the SBM-inferred posterior mean of denoised fluorescence (orange). Simultaneously recorded APs (lower). **(B)** Posterior probability of AP discharge every 10 ms given the fluorescence data (upper, blue) output by the SMC algorithm. Sum of Gaussian functions (magenta) fit to the posterior AP discharge probabilities. Note that the posterior probability of AP discharge never approaches 1 due to temporal uncertainty, but the total increase associated with each inferred AP sums to around one. By fitting a sum of unweighted Gaussian functions to the posterior probabilities, the SMC output is quantized, so that each period of increase in the posterior is interpreted as a discrete number of APs. The result (lower) is a set of Gaussian means that we interpret as AP times and standard deviations (gray) that we interpret as temporal uncertainties for each AP.

**Figure 6–Figure supplement 1.**
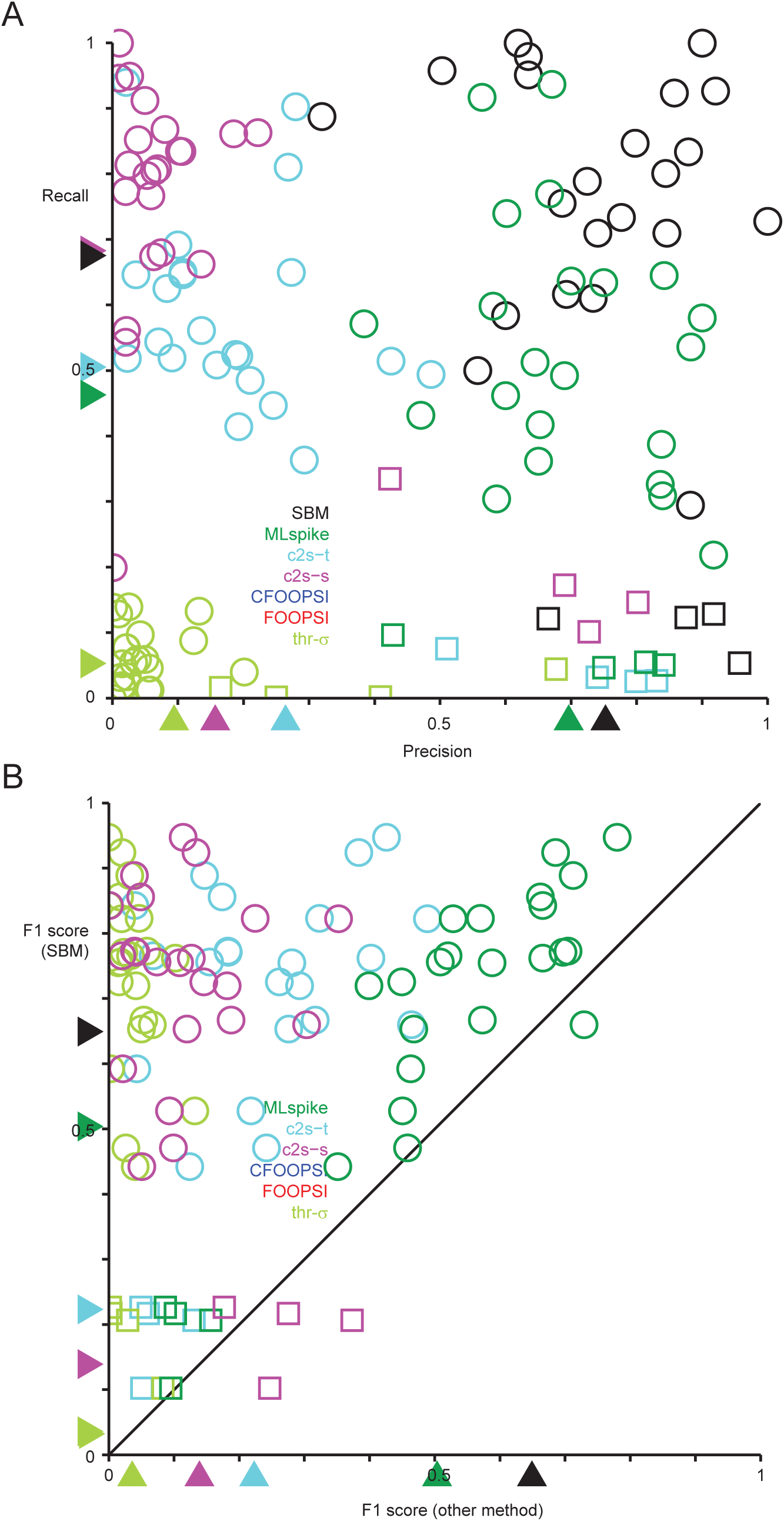
**(A)** Precision vs. recall for each neuron and AP inference algorithm. Recall is defined as the fraction of true (electrically detected) APs that are detected by the inference algorithm, and is referred to as detection rate in the main text. Precision is defined as the fraction of inferred APs that actually occurred. Both precision and recall have been computed within a maximum allowed time difference of 100 ms when matching true and inferred APs. Squares indicate interneurons. **(B)** Scatter plot comparing the performance of the SBM approach to other algorithms using the F1 score. Squares indicate interneurons. The F1 score is defined as the harmonic rate of recall and precision: 2 *·* recall *·* precision/(recall + precision).

**Figure 6–Figure supplement 2.**
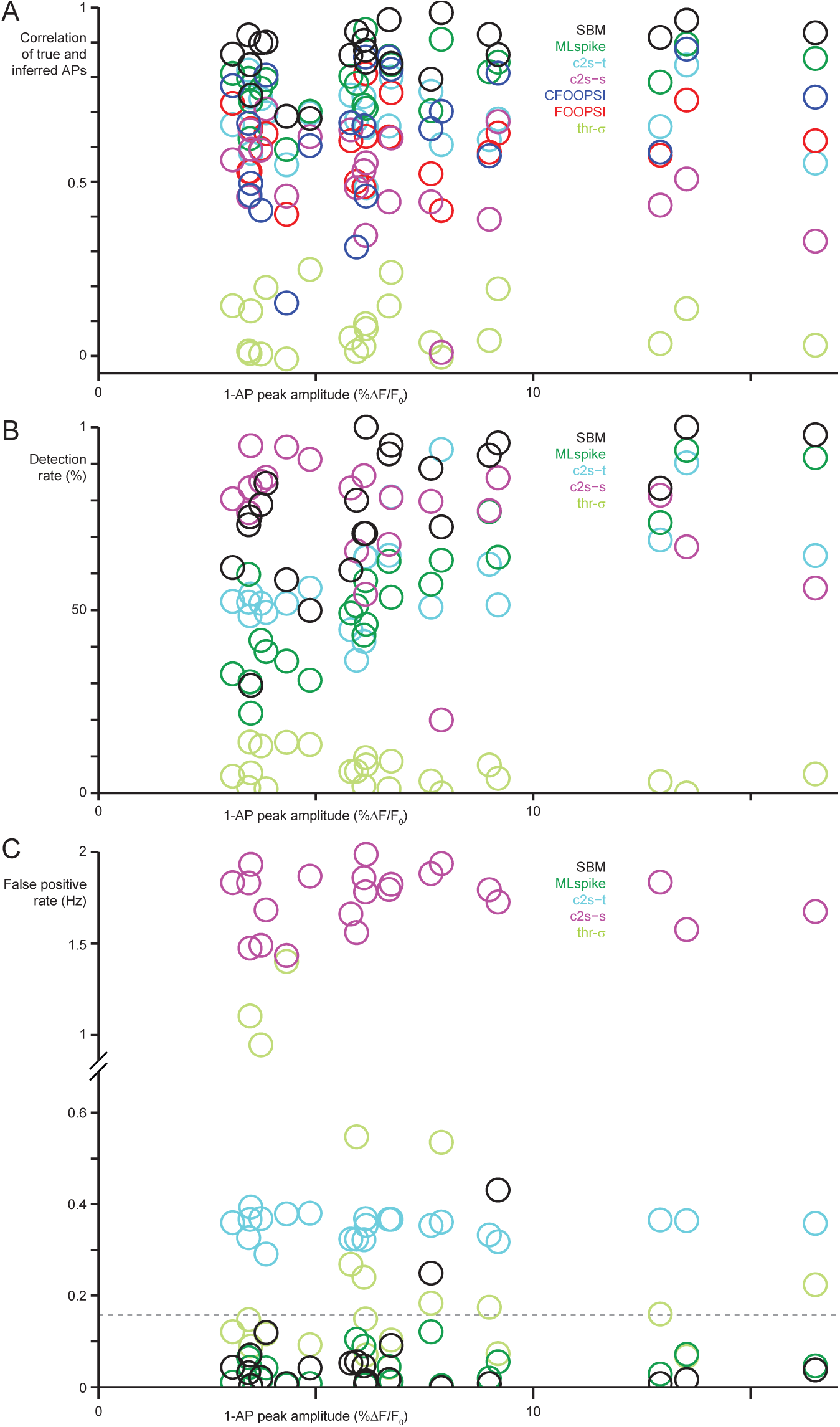
AP inference accuracy as a function of peak 1-AP fluorescence amplitude. **(A)** Peak mean fluorescence evoked by single APs in each pyramidal neuron compared to the correlation between true and inferred AP sequences for each algorithm (n = 22). **(B)** Peak mean fluorescence evoked by single APs for each neuron compared to each algorithm’s detection rate. **(C)** Peak mean fluorescence evoked by single APs for each neuron compared to each algorithm’s false positive rate. Dashed line indicates median true firing rate.

**Figure 6–Figure supplement 3.**
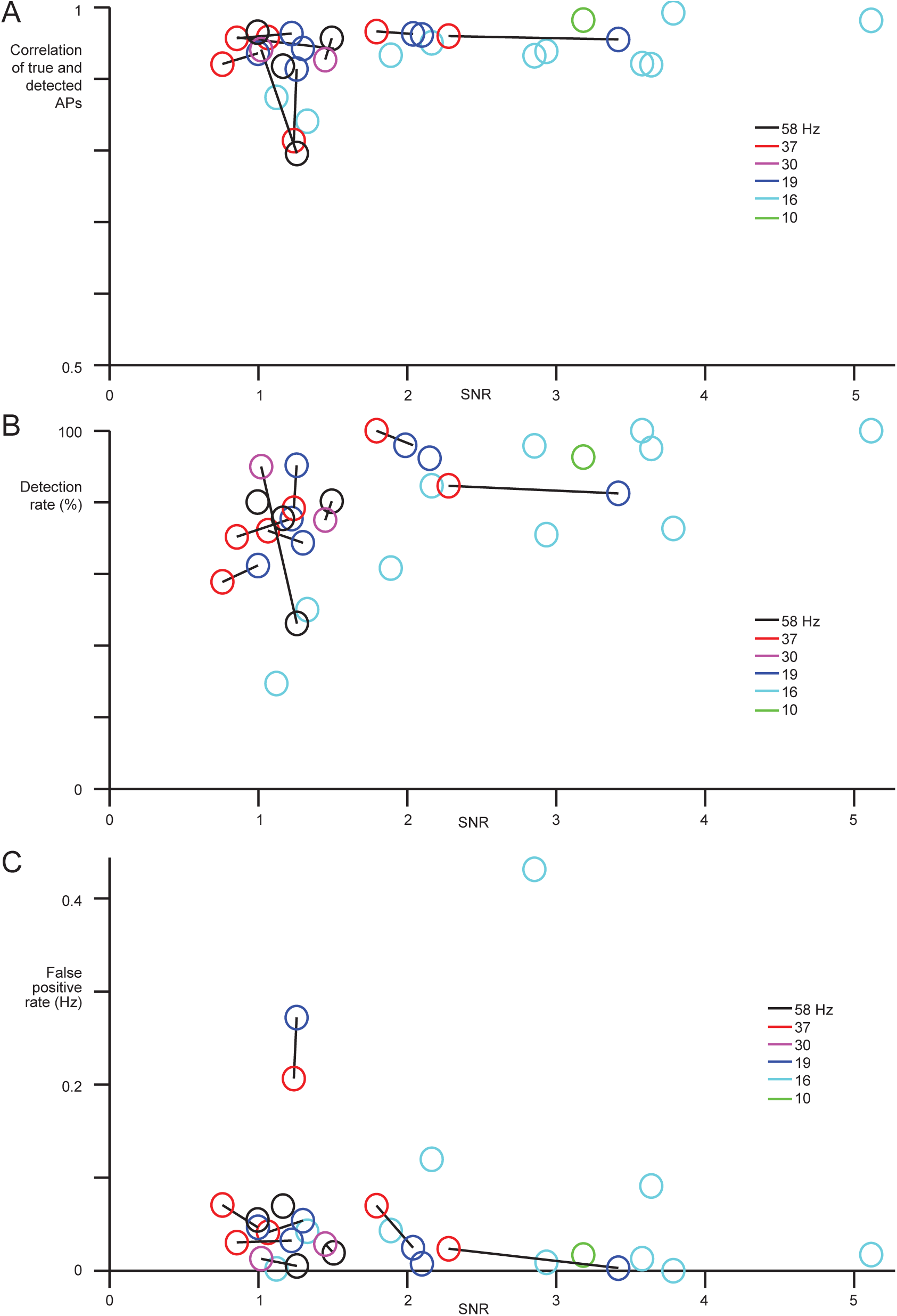
Effect of imaging frame rate and SNR on the accuracy of AP sequences inferred by the SBM. **(A)** Correlation of true and inferred APs as a function of SNR (n = 22 pyramidal neurons). Each color indicates a different imaging frame rate from 10 to 60 Hz; data from the same neuron are connected by line segments. **(B)** As in (A), for the SBM’s AP detection rate. **(C)** As in (A-B), for the rate of false positives inferred by the SBM.

**Figure 6–Figure supplement 4.**
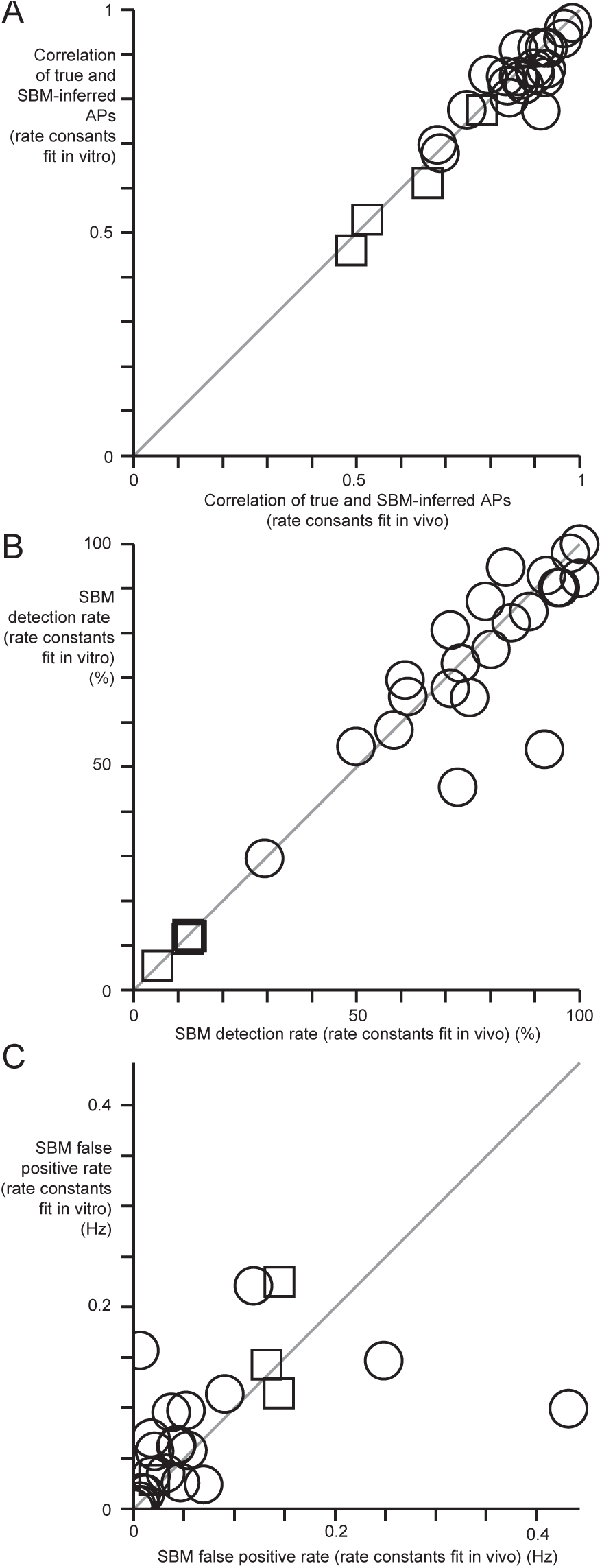
Comparison of SBM-based AP inference accuracy using rate constants fit to *in vivo* data vs. rate constants fit to *in vitro* binding assay data. **(A)** Correlation between true and inferred AP sequences for each neuron for the SBM using rate constants fit from *in vitro* binding assays (x-axis) and rate constants fit from *in vivo* data. Squares indicate interneurons. **(B)** SBM detection rate for rate constants fit *in vitro* vs. *in vivo*. **(C)** SBM false positive rate for rate constants fit *in vitro* vs. *in vivo*.

**Figure 6–Figure supplement 5.**
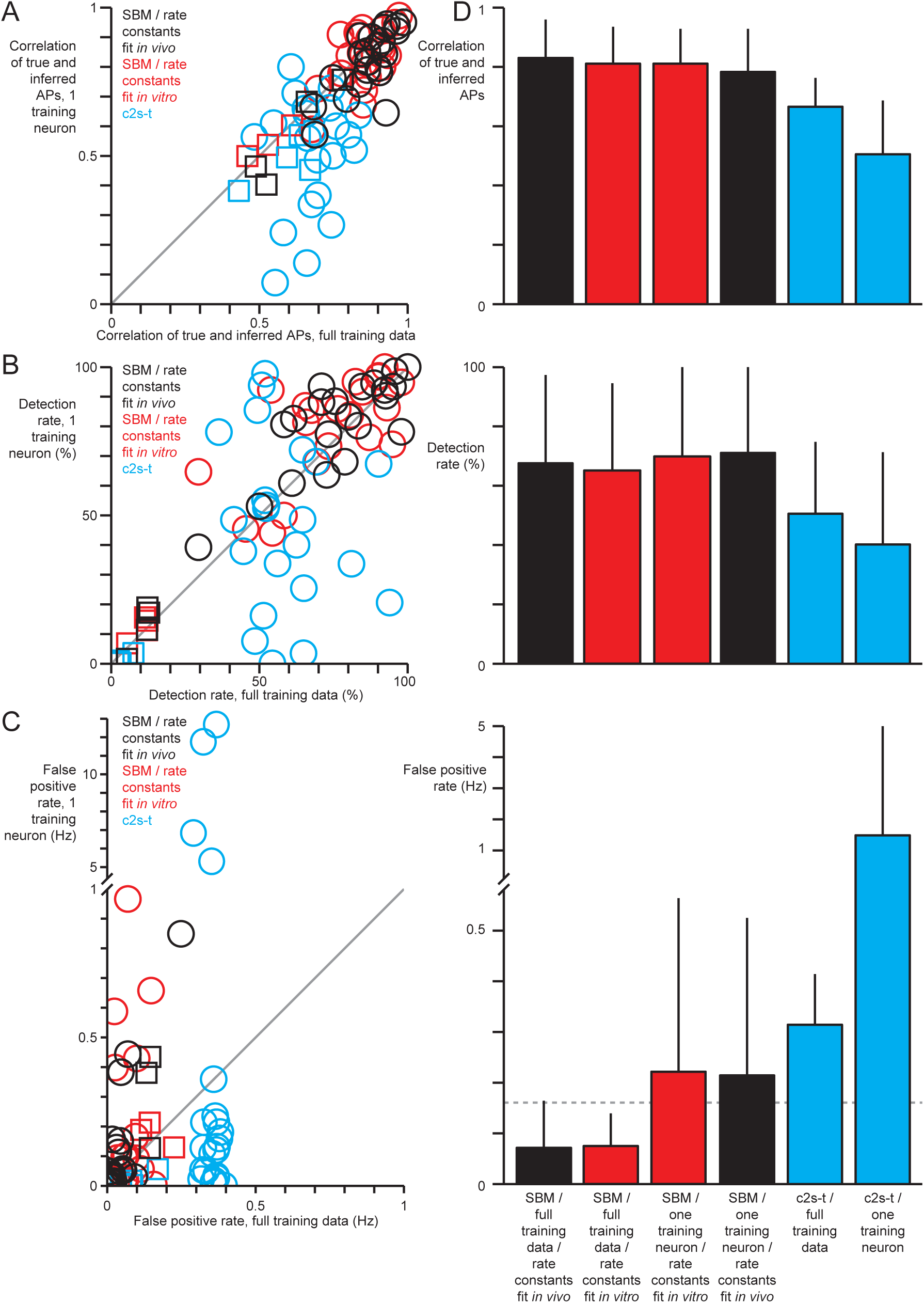
Accuracy of AP inference with one neuron of training data. **(A)** Correlation of true and inferred AP sequences for each neuron (n = 26, squares indicate interneurons) when using full training data (only one neuron at a time held out for cross-validation, x-axis) or a single training neuron (y-axis). Results are shown for the SBM using rate constants fit to *in vivo* data (black), for the SBM using rate constants fit to *in vitro* binding assays (red) and for c2s-t (cyan). **(B)** As in (A), but showing AP detection rates when using training full training data vs. a single training neuron. **(C)** As in (A-B), but showing false positive rates. **(D)** Mean and standard deviations of correlation, detection rate and false positive rate for the data shown in (A-C). Dashed line indicates true median firing rate.

**Figure 6–Figure supplement 6.**
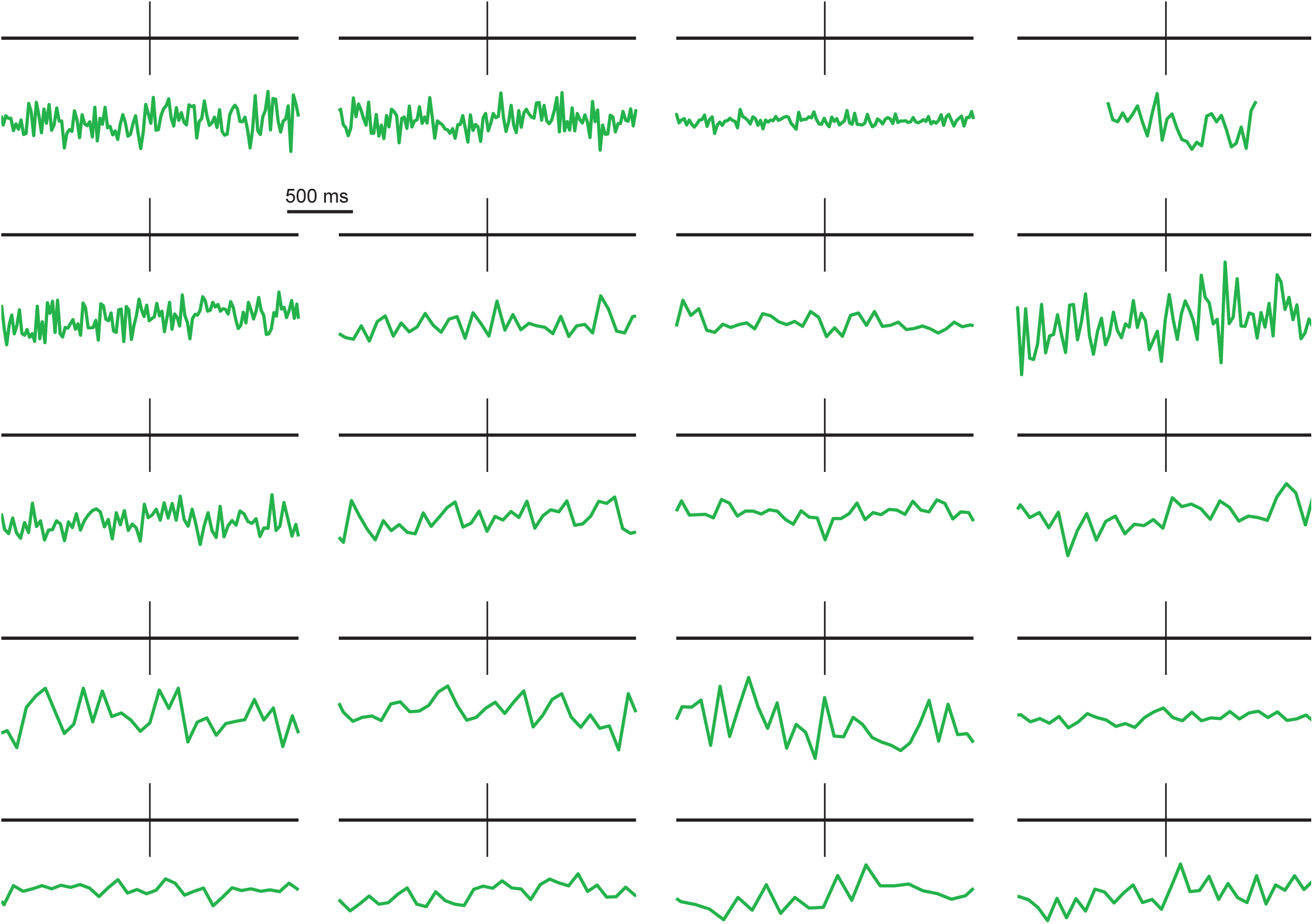
20 isolated single APs without visually apparent fluorescence increases. Each AP was recorded in a different pyramidal neuron and was chosen based on the lack of accompanying fluorescence increase. In the 2 pyramidal neurons not shown, every isolated single AP evoked a fluorescence increase that could be visually identified.

**Figure 7–Figure supplement 1.**
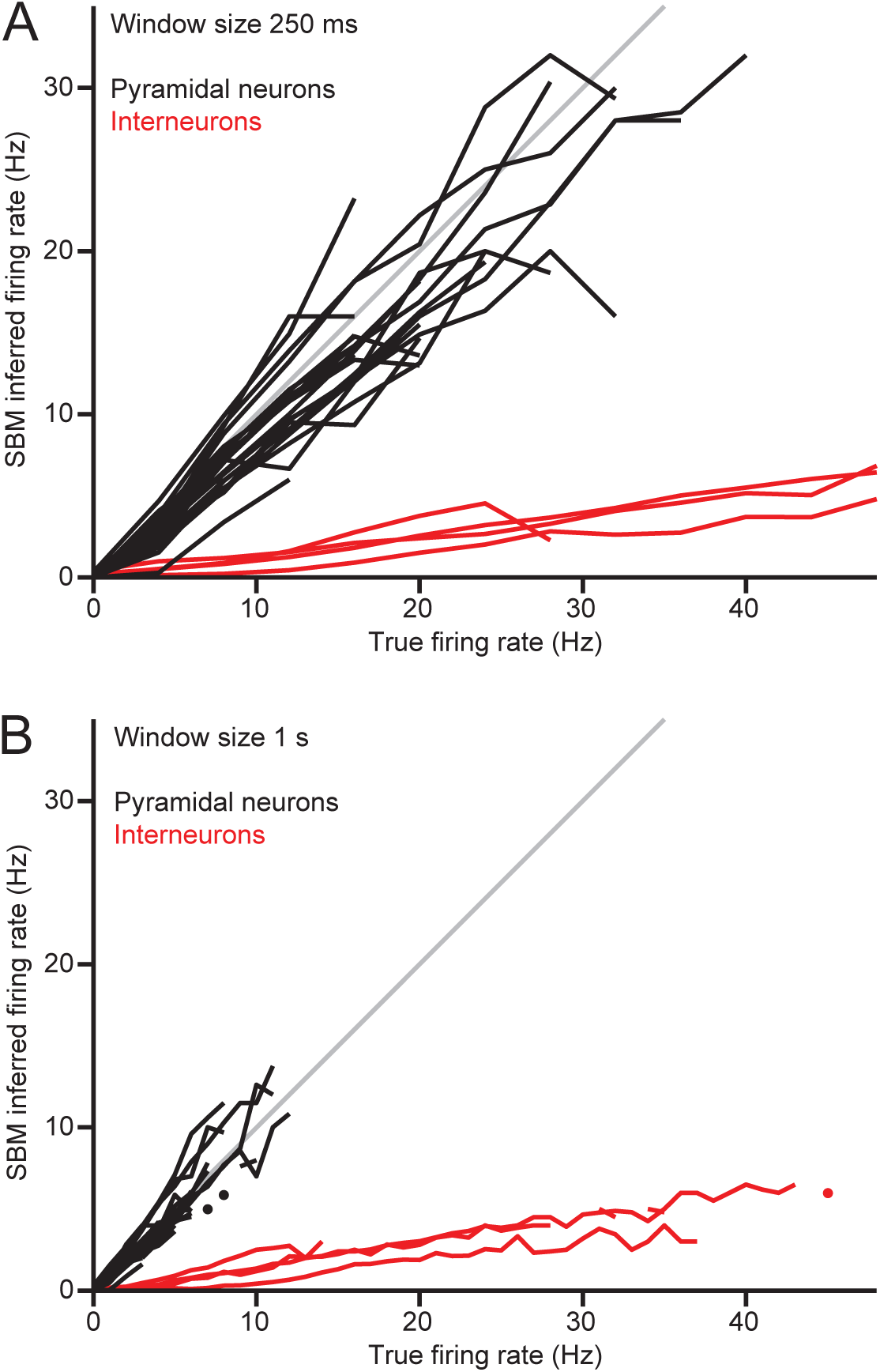
Linearity of SBM-based firing rate inference is robust to the choice of time window size. **(A-B)** As in Figure 7b, but using window sizes of 250 ms and 1s.

**Figure 7–Figure supplement 2.**
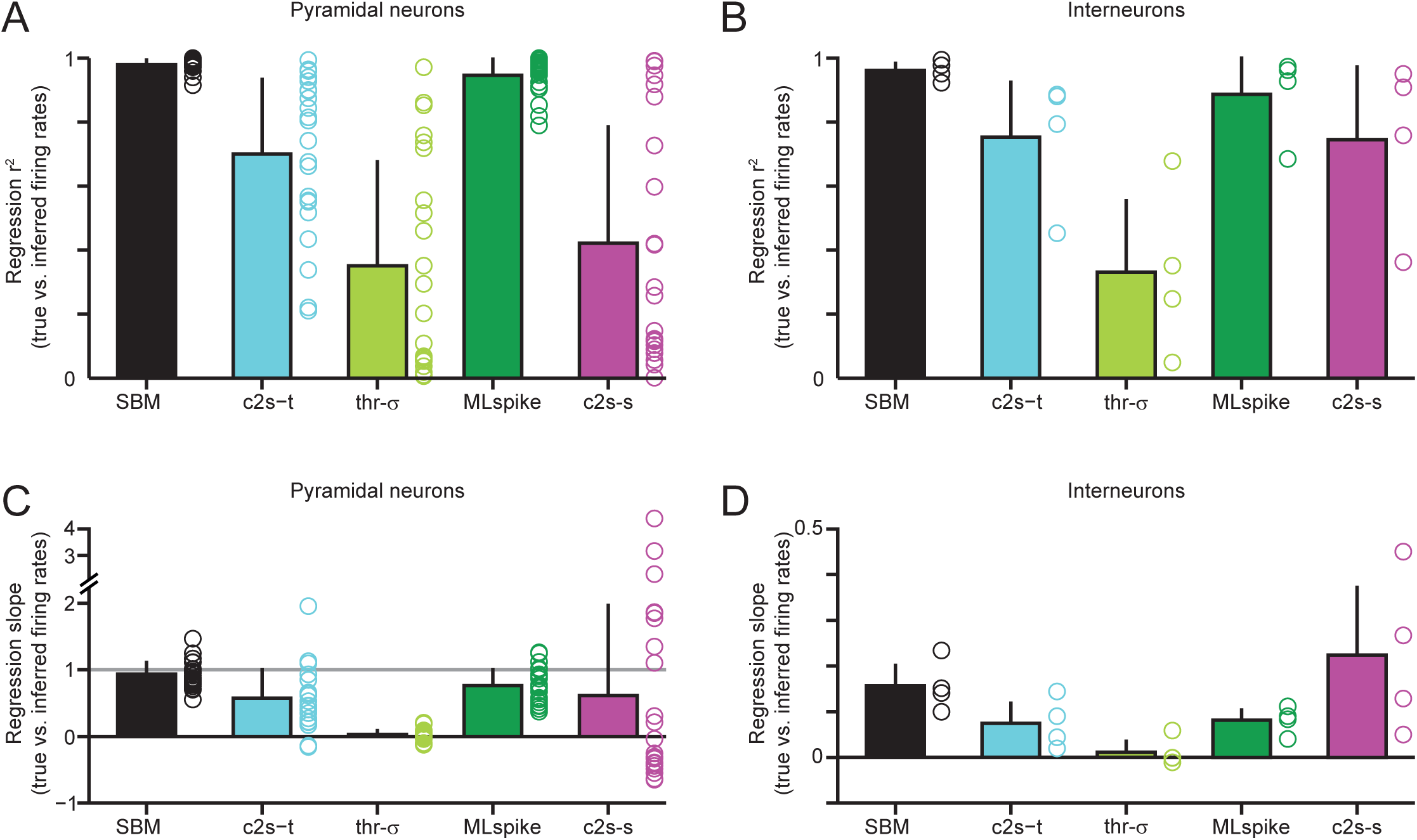
Linearity and slope of inferred firing rate as a function of true firing rate, for all inference methods (500 ms windows). **(A)** Regression *r*^2^ values for inferred firing rate as a function of true firing rate (regressions included non-zero y-intercepts). Each circle represents one pyramidal neuron, bar graph shows mean values and error bars show standard deviations. **(B)** as in (A), but for interneurons. **(C)** Linear regression slopes for inferred firing rate as a function of true firing rate, for all pyramdial neurons and each inference method. Gray line indicates a slope of 1, for which inferred firing rates have the correct units. **(D)** As in (C), but for interneurons.

**Figure 7–Figure supplement 3.**
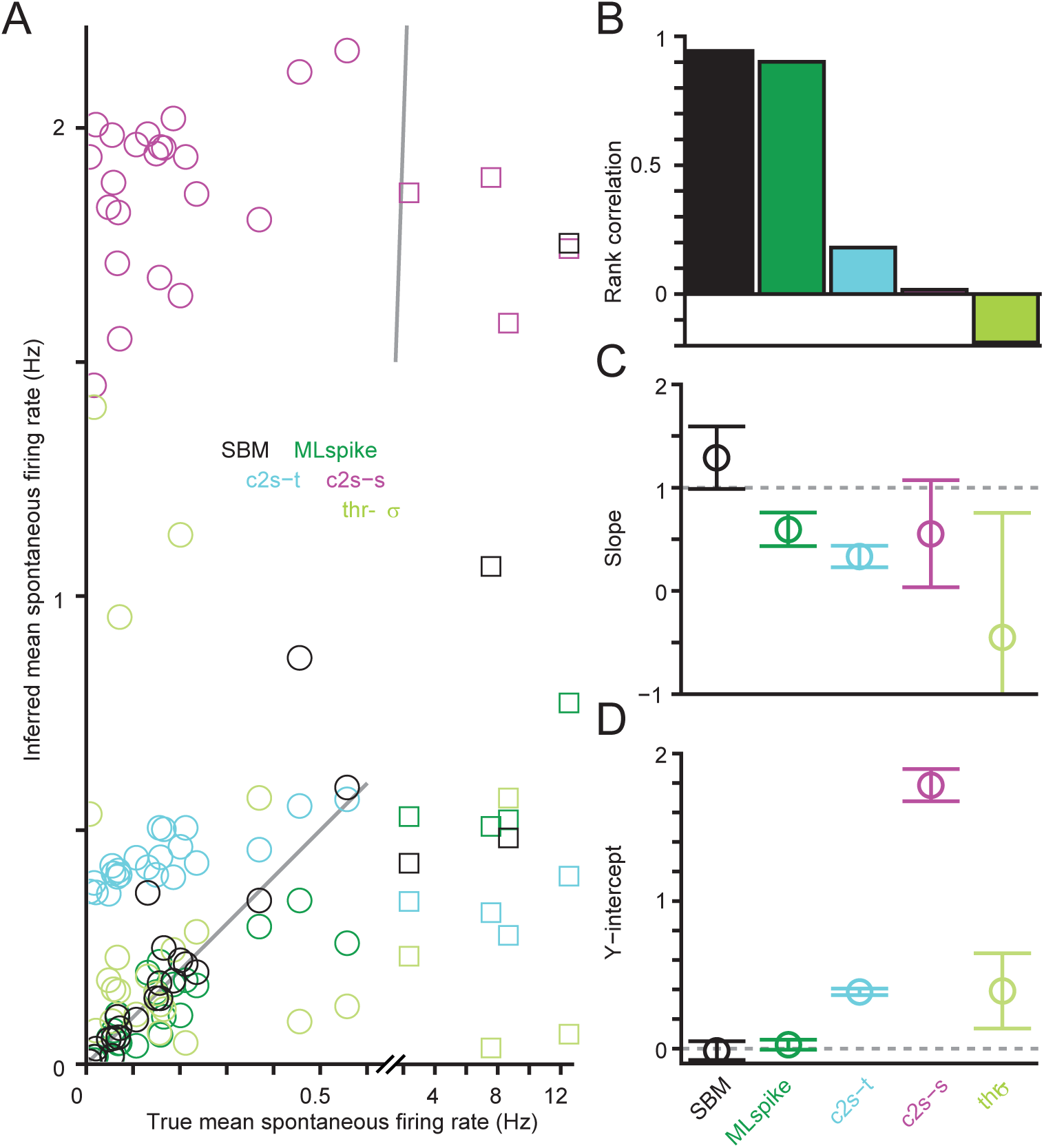
Accuracy of inferred mean spontaneous firing rates. **(A)** True vs. inferred mean spontaneous firing rates for each inference method (n = 26 neurons, squares indicate interneurons). **(B)** Rank correlation of true and estimated firing rates for each method (n= 26 neurons). **(C-D)** Slope and y-intercept of linear regressions for pyramidal neurons (n = 22). Error bars indicate 95% confidence intervals, which include both a slope of 1 and y-intercept of 0 for the SBM, but not for other methods.

**Figure 8–Figure supplement 1.**
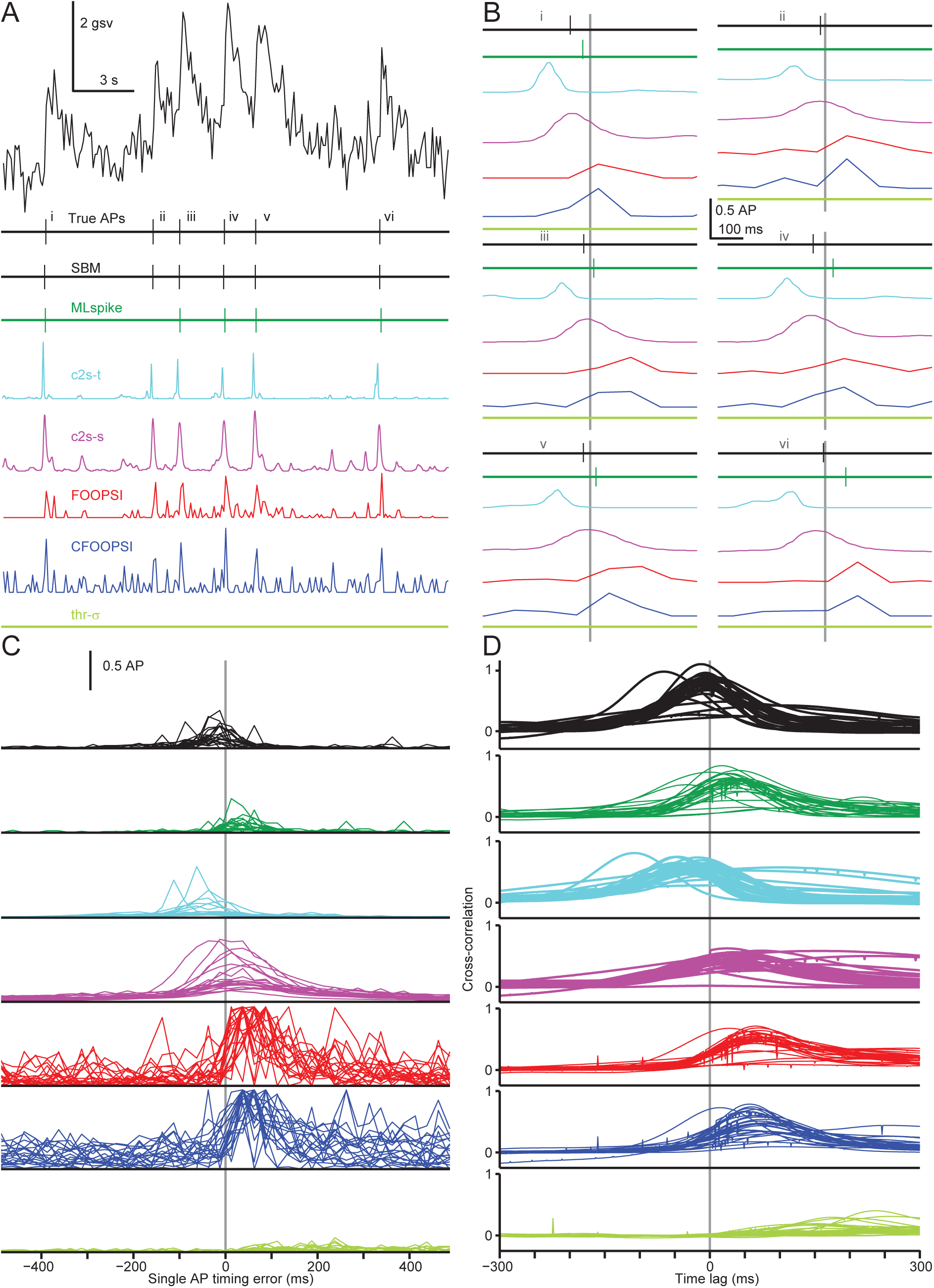
Cross-correlation and single-AP timing accuracy in individual neurons. **(A)** Six electrically recorded single APs and simultaneous fluorescence measurements (upper) with the output of AP inference algorithms (lower). Note that SBM, MLspike and thr-*σ* output an AP sequence, c2s outputs estimated spiking probabilities over time and CFOOPSI and FOOPSI have unitless outputs. **(B)** Magnified views showing AP inference results relative to electrically detected AP times (gray vertical lines) for the six single APs in (A). **(C)** Mean rates of AP inference relative to true AP times (gray vertical line) for single APs (no other APs for 1 s before and 0.5 s after), for individual neurons. Averaging these results over neurons gives the curves shown in Figure 8G. To be included in this analysis, a neuron’s data had to include at least 5 true isolated single APs, and for methods whose outputs were not unitless at least 5 total APs had to be inferred within 0.5 s of these 5 true isolated single APs. The unitless outputs of CFOOPSI and FOOPSI have been rescaled for each neuron to give a maximum average output of 1 for each neuron. **(D)** Cross-correlation between true and inferred APs for each inference method, as a function of time lag, for individual neurons. Averaging these results over neurons gives the curves shown in Figure 8D.

**Figure 8–Figure supplement 2.**
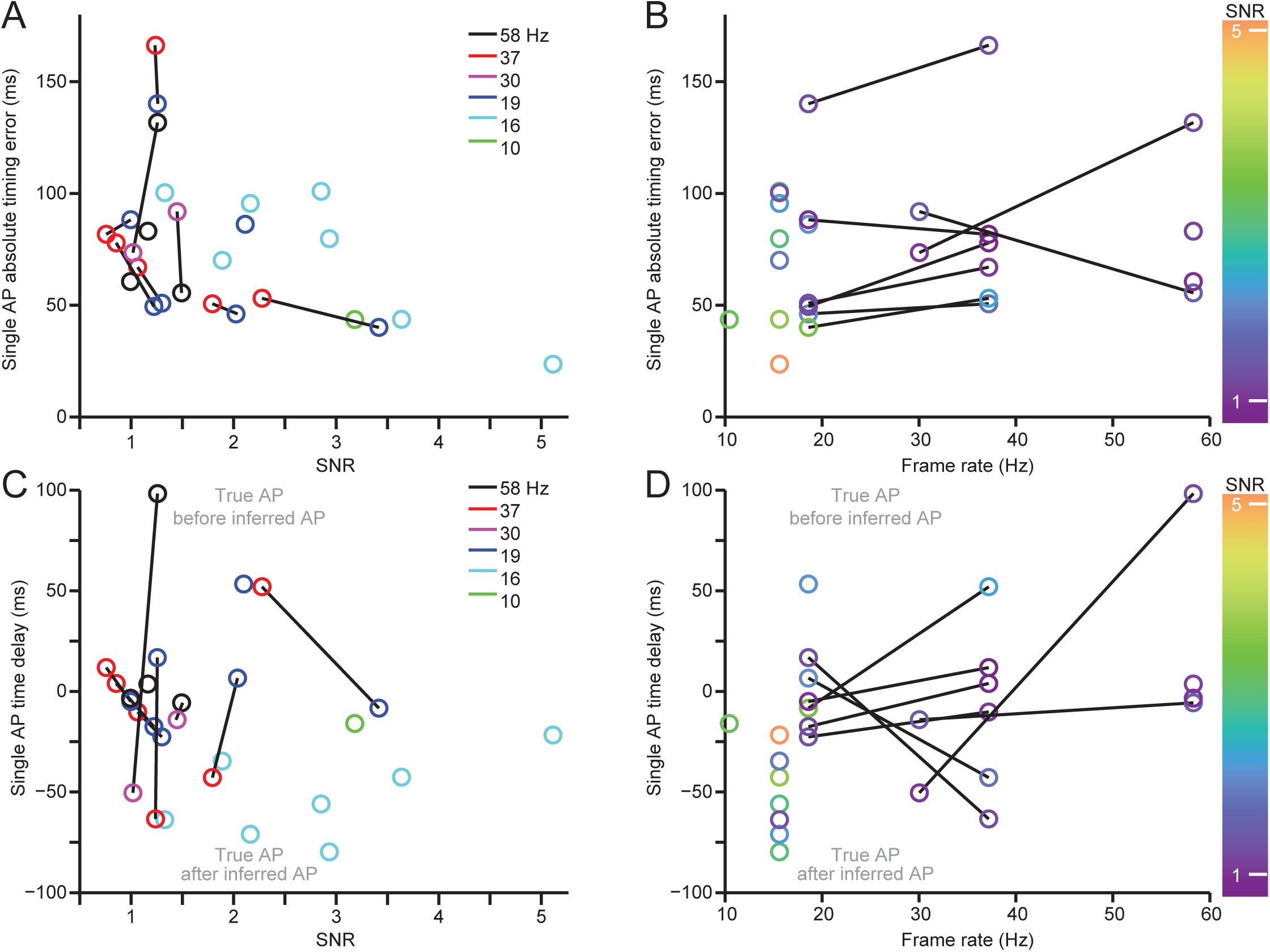
Effect of imaging frame rate and SNR on timing accuracy of APs inferred by the SBM. **(A)** Absolute timing error of isolated single APs inferred by the SBM as a function of SNR. Each color indicates a different imaging frame rate from 10 to 60 Hz; data from the same neuron are connected by line segments. Each data point corresponds to a combination of a specific pyramidal neuron with a specific imaging frame rate, and a combination was included only if it contained at least 5 true isolated single APs with at least 5 APs inferred by the SBM. **(B)** As in (A), but with imaging frame rate shown as the x-coordinate and SNR indicated by color. **(C)** Average delay from true to SBM-inferred times for isolated single APs, as a function of SNR (colors as in (A)). **(D)** As in (C), but with imaging frame rate shown as the x-coordinate and SNR indicated by color.

**Figure 8–Figure supplement 3.**
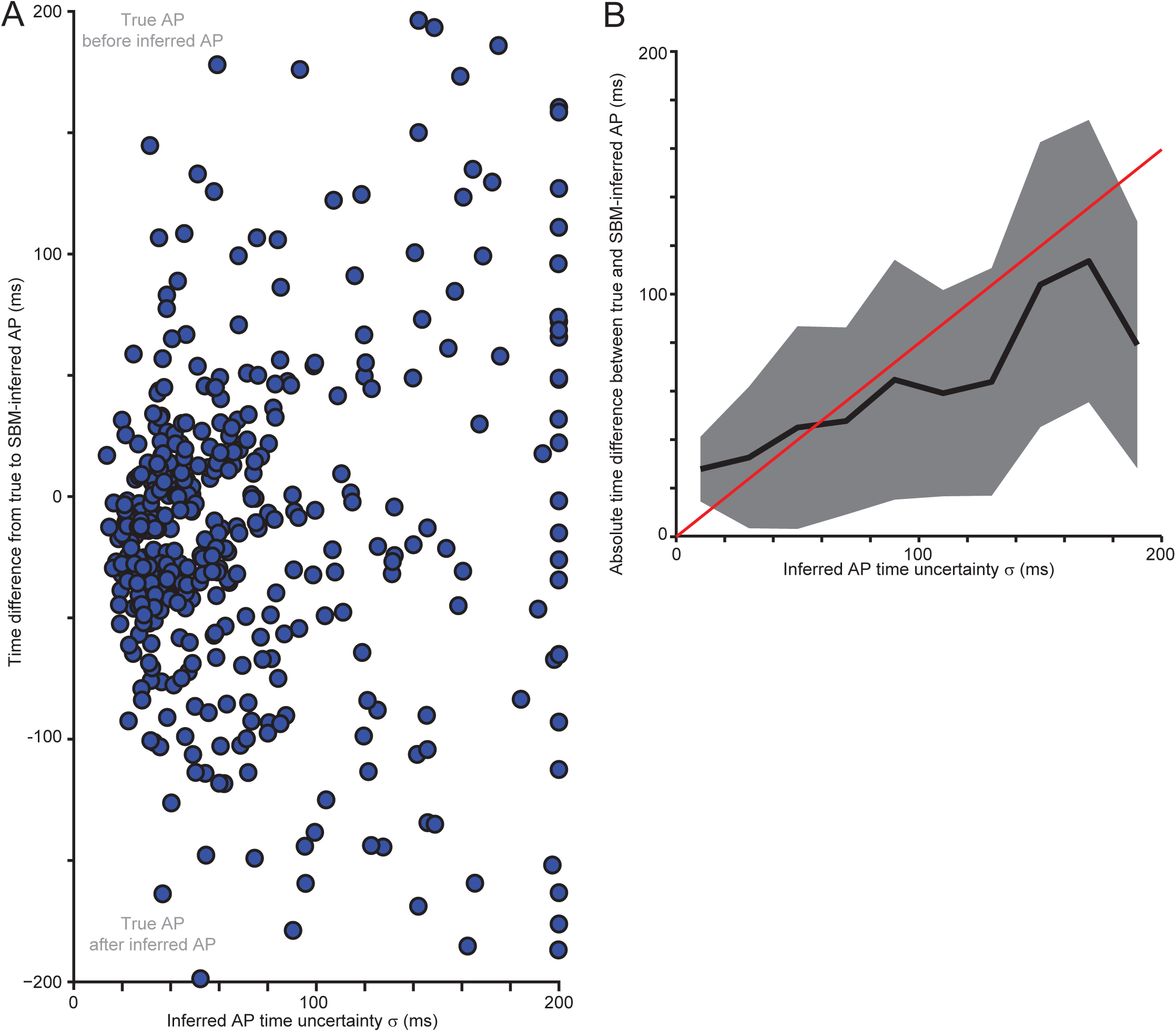
Data-based evaluation of timing uncertainty values inferred by the SBM for isolated single APs. For each AP inferred by the SBM, the algorithm also fits a standard deviation *σ* to the posterior moments inferred by the particle filter (Figure 5 - figure supplement 6), up to the maximum allowed value of 200 ms. To test whether *σ* can be validly interpreted as the width of the posterior distribution of the AP time, we compared the inferred *σ* values to the actual differences between true and inferred AP times. **(A)** Difference between true and inferred AP times as a function of inferred *σ* values (n = 399 APs from 22 neurons). APs were included in this analysis only if no other APs were present for 5 s before or 500 ms after, the SBM inferred exactly one AP within 200 ms of the true AP time and the SBM inferred no other APs within 500 ms of the true AP time. **(B)** Mean (black) and standard deviation (gray) of the absolute time difference between true and inferred AP times for the data in (A) as a function of the AP timing uncertainty output by the SBM/SMC inference algorithm. The red line shows the relation 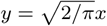, as would be expected if each *σ* value correctly describes a Gaussian posterior distribution on an AP time given the fluorescence data.

**Figure 8–Figure supplement 4.**
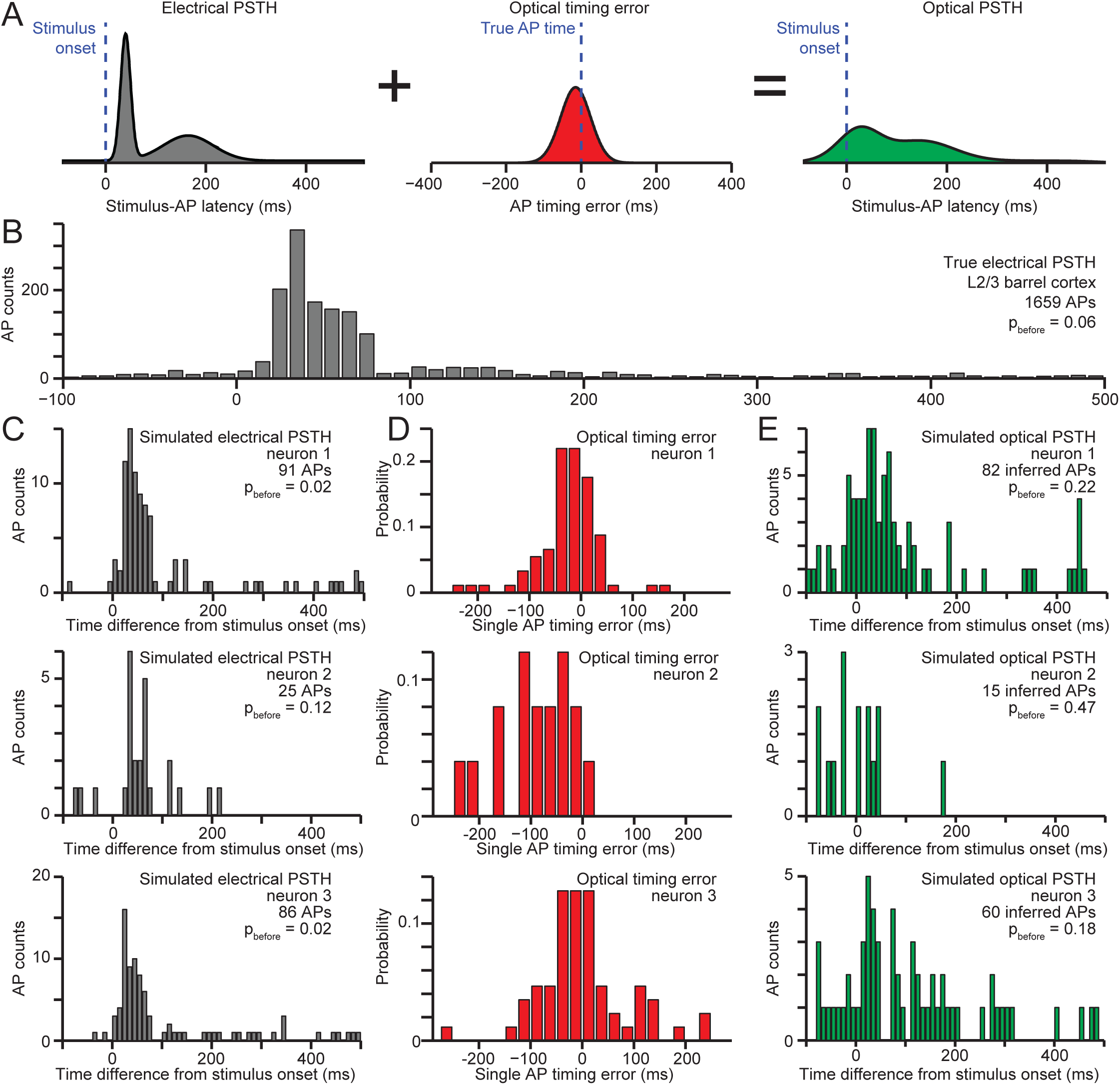
Simulations showing the effect of timing errors in optically detected APs on peri-stimulus time histograms (PSTHs). **(A)** Diagram illustrating simulation of an optical PSTH (right) by adding together stimulus→AP latencies from a true electrical PSTH (left) and timing errors for spontaneous APs detected optically in another neuron (center). **(B)** PSTH showing APs evoked by whisker deflection and recorded electrically in a L2/3 neuron in somatosensory cortex, previously published in [71]. Among APs recorded from 100 ms before the stimulus onset to 500 ms after, the fraction arising from spontaneous activity that occurred before stimulus onset was *p*_before_ = 6%. **(C)** Simulated electrical PSTHs for 3 GCaMP6s-expressing L2/3 pyramidal neurons from mouse visual cortex. For each isolated single AP recorded electrically in the neuron, an AP time relative to the simulated stimulus was drawn randomly from the PSTH shown in (B). Due to random sampling of stimulus-AP latencies and a finite number of trials, *p*_before_ ranges from 2% to 12% in these simulations. **(D)** Distribution of timing errors for isolated single APs, for the 3 neurons in (C). **(E)** Simulated optical PSTHs for the neurons shown in (C-D). For each isolated single AP recorded in the neuron, a true AP time relative to the stimulus was drawn from the PSTH shown in (B), and the results of SBM-based AP inference were used to assign optically detected AP times relative to the true AP time. Since the detection rate for single APs was less than 100%, on some stimulus trials no APs were inferred. Due to AP timing errors for the inferred APs, a greater fraction of APs were assigned to time points before the stimulus for the simulated optical PSTH than for the simulated electrical PSTH, with *p*_before_ ranging from 13% to 47%.

**Figure 8–Figure supplement 5.**
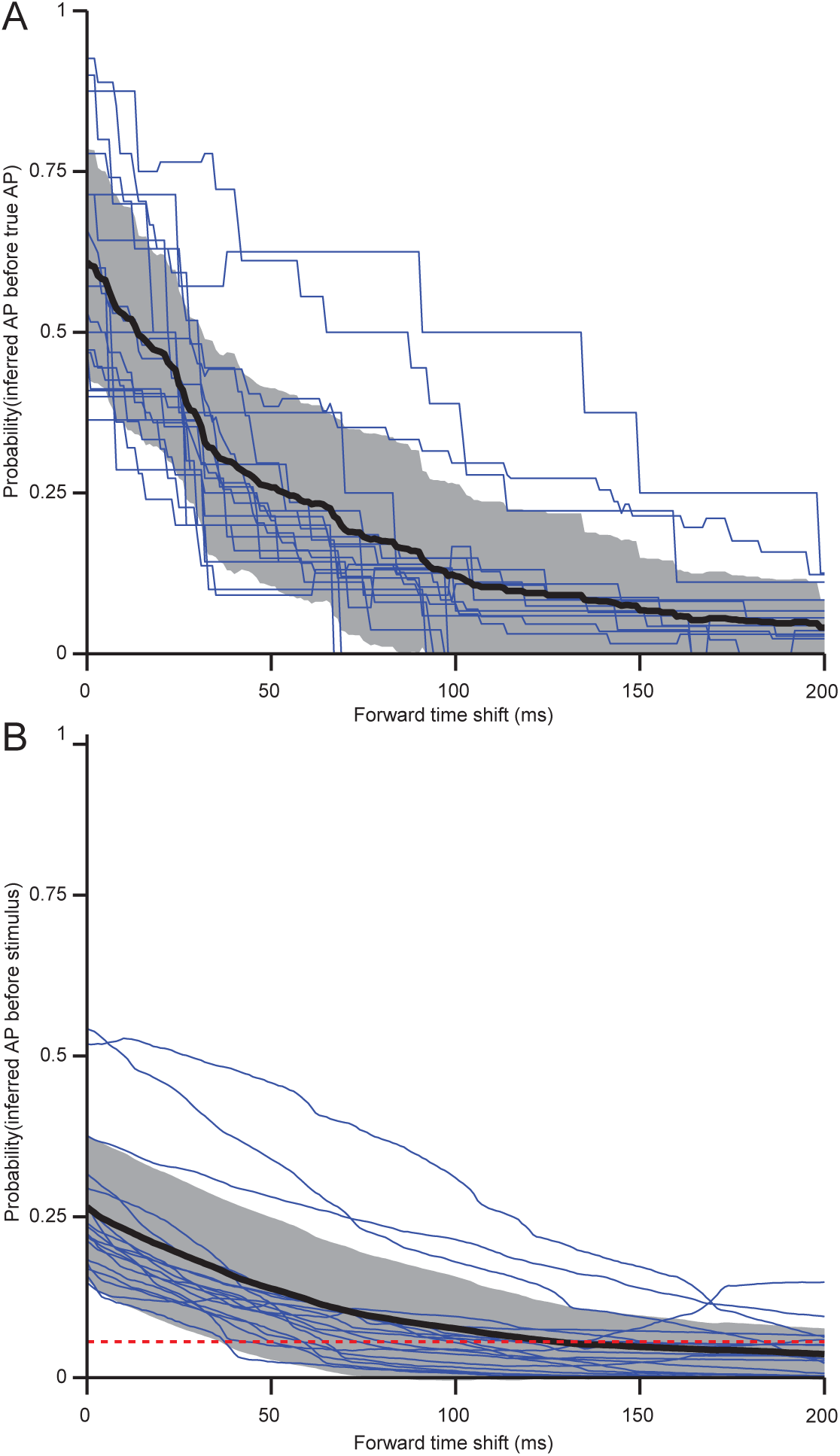
Forward time shifts limit too-early assignment of inferred APs. **(A)** Probability that SBM-inferred APs occur from 0 to 200 ms before true isolated single APs, as a function of the forward time shift applied to all inferred APs. Blue curves shows this relationship for neurons with ≥5 true isolated single APs and for which ≥5 APs were inferred within 500 ms of true single AP times (n = 18). Black curve and gray shaded region show mean and standard deviation over neurons. To limit the average probability that inferred APs occur before true APs to 0.1, a forward shift of 114 ms is required. **(B)** As in (A), but showing the probability of SBM-inferred APs occurring before a sensory stimulus for a simulated PSTH as in Figure 8 - figure supplement 4. Dashed red line indicates the true probability (0.056) that APs occurred before the stimulus in the electrically recorded PSTH used to carry out the simulations. To obtain the same probability for SBM-inferred APs on average a forward shift of 128 ms is required, while obtaining a probability of 0.1 requires a forward shift of 74 ms.

## Footnotes

1 http://commons.wikimedia.org/w/index.php?title=File:Rendered_Spectrum.png&oldid=110367311

