## Supplementary material for "Accurate action potential inference from a calcium sensor protein through biophysical modeling": data description.docx

**Accessing data**

This preliminary document gives brief instructions for accessing all data used in the manuscript. Upon publication, complete documentation of all data and metadata will be available.

For working with data files in python, the SciPy and NumPy libraries should be installed.

*In vivo* data

The file "in vivo imaging with AP times.mat" is a Matlab data file, whose contants can be loaded into a variable named "oerec" using the matlab command

**load('in vivo imaging with AP times.mat')**

or the python commands

**from scipy import io as sio**

**oerec = sio.loadmat('highfr_cascade.mat', struct_as_record=False, squeeze_me=True)['oerec']**

At this point, the data from neuron n can be accessed from Matlab as **oerec(n).** The number of separate fluoresence time series for neuron is **numel(oerec(n).data).** To access time series with index s, use

**d = oerec(n).data(s)**

Then fluorescence values can be accessed as **d.f,** the fluorescence signals removed during feature extraction as **d.f_removed**, the times when the neuron was scanned as **d.t**, and AP times as **d.spiketimes**. Additional metadeta are available in **oerec(n).info** and **oerec(n).data(s).info.**

*In vitro* data

*In vitro* data can be accessed using python using the commands

**import pickle**

**with open(‘in vitro binding assays.pickle’, 'rb') as fid:**

**experiments = pickle.load(fid)**

The variable **experiments** now contains a list of python dict objects. Import fields include ‘type’ (spectra, itc, or stopped flow) and ‘data’. Additional fields describe experimental parameters, reagent concentrations, etc.
